# *ArchaicSeeker* 3.0: A deep-learning framework for scalable, haplotype-resolved inference of archaic introgression

**DOI:** 10.64898/2026.05.05.722798

**Authors:** Baonan Wang, Chang Lei, Huanyu Lin, Shaokang Shi, Xixian Ma, Weishun Zeng, Kai Yuan, Xumin Ni, Shuhua Xu

**Author notes:** Correspondence and requests for materials should be addressed to S.X. These authors contributed equally to this work (Baonan Wang, Chang Lei, Huanyu Lin).

## Abstract

Archaic introgression has left a significant mark on human genetic diversity, but reliably identifying introgressed segments remains a major challenge, especially with complex demographic histories and limited sample sizes. Existing methods often rely on demographic assumptions or cohort-specific parameter fitting, which compromises robustness and scalability. We introduce *ArchaicSeeker 3.0* (AS3), a deep-learning framework designed for haplotype-resolved detection of archaic introgression. AS3 integrates a tract-scale sequence model with an overlap-aware reassembly approach and boundary refinement, enabling accurate, boundary-coherent reconstruction of introgressed segments across diverse genomic contexts. By leveraging a simulation-trained model, AS3 avoids inference-time recalibration, offering stable performance across unrepresented demographic scenarios and small cohorts. In extensive simulations, AS3 outperforms existing methods in precision, recall, and F1 score, while providing more continuous segments with accurate boundary localization. It demonstrates robustness in small-target regimes and varying marker densities. Applied to 3,453 genomes from 209 populations, AS3 shows strong concordance with existing introgression callers and identifies additional introgressed regions, including high-frequency AS3-specific introgressed segments supported by locus-level haplotype and phylogenetic analyses. AS3 provides a scalable, robust solution for detecting archaic introgression from single individuals to large biobank datasets, marking a significant advancement in the field of local ancestry inference and opening new possibilities for the study of human evolutionary genetics.

*ArchaicSeeker* 3.0 is available at https://github.com/Shuhua-Group/ArchaicSeeker3.0.

## Introduction

Extensive archaeological and genetic evidence suggests that around 600–700 thousand years ago, a distinct hominin population diverged from the ancestral lineages of modern humans in Africa. This group subsequently dispersed beyond Africa and, over time, differentiated into the Neanderthals in western Eurasia and the Denisovans in eastern Eurasia^1–5^. Anatomically modern humans (AMH) later arose in Africa and expanded across the continent and beyond^6,7^. During this expansion, AMH encountered archaic populations—including Neanderthals, Denisovans, and possibly other groups outside Africa. By the Late Pleistocene (e.g., Neanderthals ∼40 ka), these archaic groups had largely disappeared due to a combination of climatic, epidemiological, and competitive factors^8–10^. Ancient DNA analyses nonetheless demonstrate that introgression occurred and left a measurable legacy in present-day genomes^1^. On average, non-African populations carry ∼1–2% Neanderthal ancestry, with additional Denisovan ancestry in some groups (e.g., parts of Oceania)^1,11,12^. Functional studies indicate that this archaic legacy contributes to phenotypic diversity and adaptive traits in contemporary humans^13–16^.

To elucidate this legacy, a primary objective is to identify and characterize archaic DNA within modern genomes. This task falls under archaic local ancestry inference (LAI), which segments genomes and assigns each segment to an archaic source (or to non-archaic background). Contemporary methods largely fall into two computational paradigms, alongside a complementary IBD-based approach. LD-based methods such as S* and Sprime leverage extended haplotype structure and allele-frequency patterns that tend to persist within introgressed regions^17–21^. Probabilistic graphical models, typically Hidden Markov Models (HMM) or Conditional Random Fields (CRF), infer segment boundaries by modeling transitions along the genome, as implemented in hmmix and ArchaicSeeker 2.0^22,23^. A distinct, complementary line of work typified by IBDMix detects identity-by-descent sharing between modern haplotypes and ancient references^24^. Together these strategies emphasize different signals and have enabled increasingly fine-grained reconstructions of archaic contributions.

Despite sustained progress, several practical and methodological gaps remain. First, many current methods rely on explicit demographic assumptions or cohort-level parameter calibration, which can reduce robustness when the true history departs from the assumed model, such as under mixed archaic sources, multiple introgression pulses, or continuous gene flow. Second, an underappreciated limitation is the data regime: methods that depend on large target cohorts or population-specific fitting are difficult to apply to under-sampled populations, rare groups, or analyses with only a small number of target genomes. Third, many approaches still do not fully exploit multi-locus haplotypic context over introgressed tracts. Methods based on summary statistics, local heuristics, or simplified sequence models can miss informative multilocus linkage patterns, leading to incomplete recovery of introgressed tracts or reduced sensitivity to weak signals. Fourth, boundary placement remains a persistent bottleneck: tract calls often become fragmented or show inconsistent endpoints across tiled or window-based scans, limiting downstream interpretation of tract length, overlap structure, and locus-specific effects. More broadly, these challenges are not unique to archaic introgression, but reflect a wider class of population-genetic inference problems in which local ancestry states or segment labels must be inferred under complex and only partially specified histories. In such settings, the key difficulty is often not the absence of theoretical models, but the limited robustness of explicitly parameterized models when applied across heterogeneous demographic scenarios and target-cohort regimes. Collectively, these considerations call for methods for archaic local ancestry inference that remain robust to demographic misspecification without inference-time cohort-specific refitting, operate reliably in few-target regimes, leverage multi-locus haplotypic context efficiently, and produce coherent and stable segment boundaries.

Deep learning provides a flexible route for inferring ancestry patterns directly from genetic sequences. Transformer-based and convolutional models have shown utility in regulatory genomics, local ancestry inference, and adaptive introgression tasks^16,25,26^. For this class of problems, deep learning is valuable not simply because it replaces one model family with another, but because it can learn discriminative sequence-level rules from diverse simulated histories without requiring the full inference procedure to be re-parameterized for each new dataset. For archaic local ancestry inference, however, two practical requirements remain difficult to satisfy simultaneously: modeling sufficiently long SNP contexts to capture extended haplotypic information across introgressed tracts, and producing stable site-wise predictions with coherent segment boundaries under windowed inference. These considerations motivate efficient sequence architectures that can model broader haplotypic context while maintaining practical scalability; the specific backbone design is described in the Methods.

Here we present ArchaicSeeker 3.0 (AS3), a simulation-trained, two-stage deep-learning framework for haplotype-resolved inference of archaic introgression from SNP sequences using African outgroup and archaic references. Although developed and validated here for archaic introgression, AS3 is motivated by a broader simulation-to-inference perspective for population-genetic problems that can be formulated as local sequence- or tract-level labeling tasks. AS3 is designed to address four practical challenges simultaneously: capturing multi-locus haplotypic context, mitigating window-boundary artifacts, maintaining robustness to demographic-model misspecification without inference-time demographic cohort-specific fitting or recalibration, and supporting few-target analyses. The framework combines a sequence backbone for within-window inference with an overlap-aware chromosome reassembly procedure and a lightweight refinement module to improve cross-window coherence and boundary localization. Across heterogeneous demographic simulations, AS3 preserves a favorable precision–recall balance and achieves the strongest archaic-seeking F1 among compared methods, while producing tract calls with improved continuity and tighter boundary placement. Because tract length distributions encode information about the timing, multiplicity, and structure of introgression events, improved boundary localization is important not only for cleaner segment calls but also for downstream historical interpretation. Notably, AS3 remains stable in low-target settings (including *n_tgt_* ≤ 10) and can be applied without cohort-specific parameter fitting or recalibration at inference time. Together, these results position AS3 not only as a practical framework for archaic introgression inference, but also as an example of how simulation-trained deep-learning models can generalize across demographic mismatch in haplotype-level population-genetic sequence inference.

## Materials and Methods

### Model architecture

ArchaicSeeker 3.0 is a neural network-based algorithm designed for the detection of archaic introgression events. The model processes genomic data from African (AFR), Neanderthal (NEAN), Denisovan (DEN), and the query sequence (the input admixed sequence), subsequently identifying potential archaic introgressed segments. It is composed of two primary components: a base model followed by a smoother layer (Fig 1). The base model aggregates information within windows and generates preliminary predictions regarding whether each site within the window is an archaic introgression site. The smoother layer, on the other hand, incorporates information across windows to refine these initial predictions. ArchaicSeeker 3.0 demonstrates significant generalization capabilities, being applicable to a broad range of demographic models upon completion of training in a specific setting, without the necessity for retraining or adjustments.

**Fig. 1.**
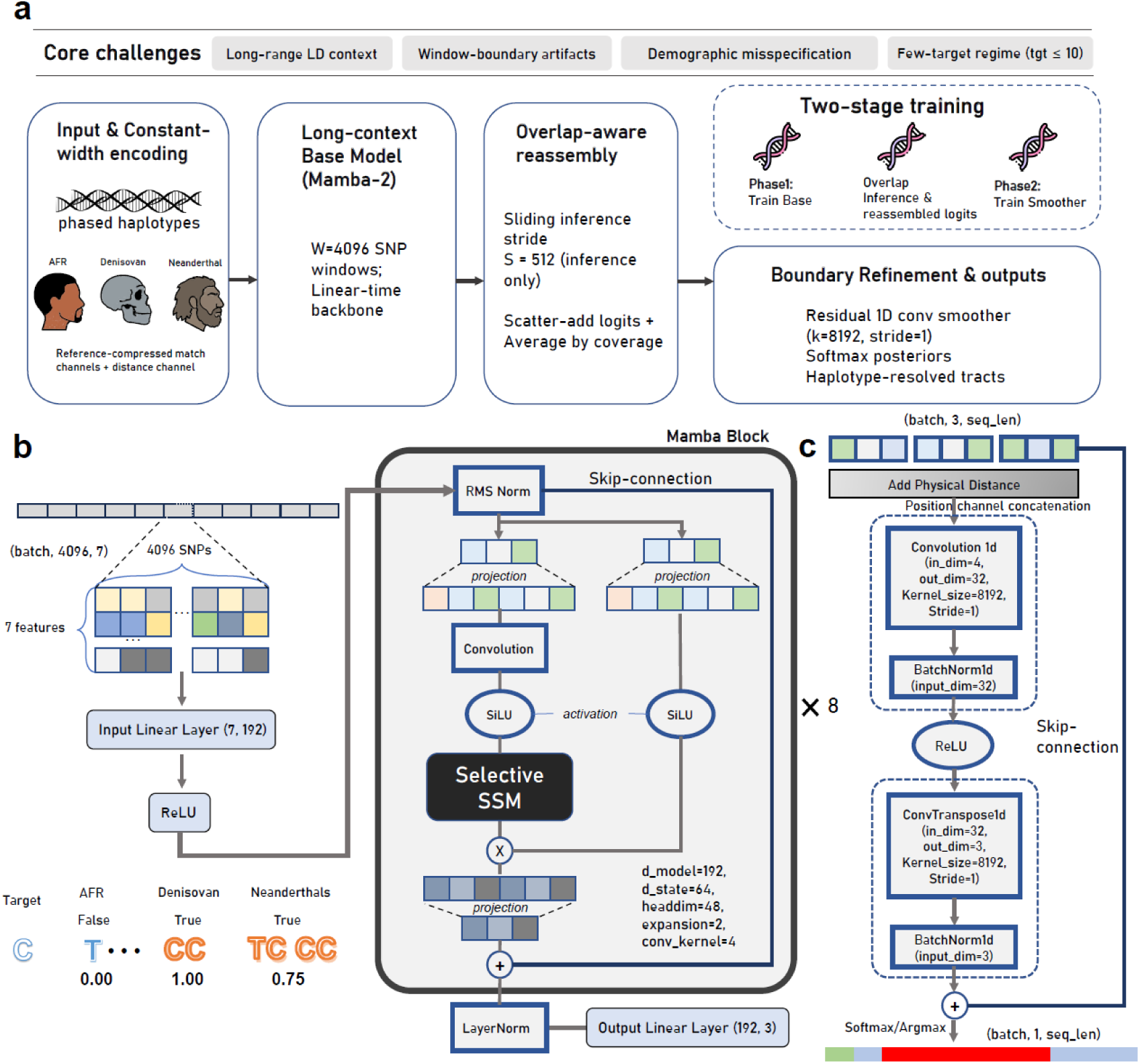
Overview of the ArchaicSeeker 3.0 framework. (a) System overview and design targets. ArchaicSeeker 3.0 (AS3) is a two-stage framework for haplotype-resolved archaic introgression inference designed to address four practical challenges: capturing long-range LD context, mitigating window-boundary artifacts, remaining robust to demographic model misspecification without inference-time demographic re-fitting, and operating in few-target regimes (*n_tgt_* ≤ 10). AS3 takes phased target haplotypes together with African outgroup, Denisovan and Neanderthal references, constructs a constant-width, reference-compressed per-site encoding plus a distance/position channel, and applies a long-context Base model to fixed windows (*W* = 4,096 SNPs) to produce per-site logits. Chromosome-wide inference uses overlapping windows with stride *S* = 512 (inference only); per-window logits are scatter-added and averaged by coverage to yield a single reassembled logit track aligned to the original SNP axis. A lightweight residual 1D convolutional Smoother then refines boundary coherence, after which Softmax posteriors (and argmax labels) are converted into haplotype-resolved tracts by downstream tract decoding controlled by merge hyperparameter (*δ*). Training proceeds in two stages: Phase I trains the Base model on simulations spanning multiple demographic scenarios; Phase II trains the Smoother using reassembled logits generated by overlap inference. (b) Base model architecture (Mamba-2). Within each 4,096-SNP window, the input tensor has shape (batch, 4,096, 7): six reference-compressed match channels (max/mean over AFR, DEN and NEAN) plus one distance-derived channel. The 7-channel sequence is projected to the model dimension (192) and processed by a stack of Mamba blocks (×8) that combine normalization, gated projections, local convolutional mixing, and selective state-space dynamics to propagate long-range context while preserving sequence length. The model outputs per-site logits over three ancestry states (non-archaic, Denisovan, Neanderthal) via a final normalization and an output linear layer. (c) Boundary refinement module (Smoother). The Smoother operates on the reassembled chromosome-length logits by concatenating a position/distance channel to form a 4-channel 1D input. A wide-kernel Conv1d (*k* = 8,192, stride = 1) with BatchNorm and ReLU integrates cross-window context; a matched ConvTranspose1d (*k* = 8,192, stride = 1) restores the three logit channels, and a residual connection adds the original logits to the refined output. Final per-site posterior probabilities are obtained via Softmax (and discrete labels via argmax), and tract decoding converts per-site labels into segments using *δ*.

### Base model architecture

The base model uses Mamba-2, a selective state-space sequence architecture, as the long-context backbone for within-window inference. Selective state-space models propagate information through the sequence using input-dependent state updates, allowing the model to retain and prioritize context relevant to the current site while avoiding the quadratic cost of full self-attention^27^. We chose Mamba-2 because it provides an efficient and practically stable implementation for long one-dimensional sequences, with improved parallelization and hardware efficiency relative to the earlier formulation^28^. These properties make it well suited to SNP-window modeling in archaic local ancestry inference, where informative haplotypic patterns can extend across long genomic intervals but site-wise ancestry logits must still be produced for downstream chromosome reassembly and boundary refinement. For each window of 4096 SNPs, we construct an input matrix 𝑋 ∈ ℝ^4096×7^. This window length serves as a practical working configuration in the present framework rather than a universally optimal constant. It was chosen to balance the need for sufficiently long haplotypic context for tract-scale inference, computational efficiency, and compatibility with the downstream overlap-aware reassembly and boundary-refinement design. In particular, the smoother uses a kernel size of 8,192 so that its receptive field spans exactly two adjacent 4,096-SNP windows, enabling information to be integrated across neighboring windows during refinement while avoiding padding-induced boundary artifacts. At each site, we compute the max and mean of the binary match indicators between the query haplotype and each of three reference sets, African (AFR), Denisovan (DEN), and Neanderthal (NEAN), yielding 6 channels, and append a relative genetic-distance channel. Relative distance is obtained by converting physical to genetic distance using 1 cM/Mb = 0.01 Morgan/Mb and subtracting the within-window minimum:

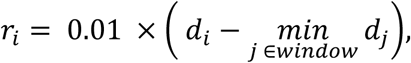

where 𝑑_𝑖_ is the physical distance in Mb; when empirical recombination maps are available, they are used to derive genetic distances. This encoding provides (i) site-level archaic distinctiveness (max), (ii) allele-frequency context (mean), and (iii) positional anchoring and linkage information (relative distance), while compressing an arbitrary number of reference haplotypes into a fixed-width representation.

The network maps 7 features to 𝑑_𝑚𝑜𝑑𝑒𝑙_=192 via a linear layer with ReLU, followed by 8 stacked Mamba-2 blocks (each with 𝑑_𝑠𝑡𝑎𝑡𝑒_ = 64, expansion = 2, depthwise conv kernel = 4, residual connections, and LayerNorm). A final linear layer outputs per-site logits over non-archaic, Denisovan, Neanderthal with shape 4096×3. Sequence length is preserved throughout and no explicit positional encodings are required (order is captured by the state-space dynamics, while the relative-distance channel provides local anchoring). The computation is *O(W)* in window length *W*, enabling efficient scaling to long contexts.

By combining reference compression with long windows, the base model reduces dependence on reference sample size without sacrificing accuracy and maintains a stable I/O contract for the downstream multi-offset sliding-window smoother, which consolidates overlapping predictions into coherent segment calls.

### Windowed inference with overlap-aware reassembly

At inference, windows of *W* = 4096 SNPs are slid with stride *S* = 512, yielding overlapping contexts. Per-window logits are projected back to the global SNP coordinate axis and aggregated across coverage: logits from all covering windows are scatter-added at each index and divided by the coverage count. Minimal right-padding makes the sequence length a multiple of S before windowing; padding is removed after reassembly. This produces a contiguous per-site logit series aligned to the original SNP coordinates while retaining the benefits of long-context modeling within windows.

### Boundary Smoother architecture

After reassembly, we apply a one-dimensional convolutional smoother to improve boundary coherence across neighboring windows (Fig. 1). The smoother operates on reassembled site-level logits rather than on raw SNP features: specifically, it takes 𝑌_𝑏𝑎𝑠𝑒_ ∈ ℝ^𝐵×3×𝐿^ and concatenates a Morgan-scaled absolute-position channel 𝑝 ∈ ℝ^𝐵×1×𝐿^ , forming an input tensor in ℝ^𝐵×4×𝐿^ . The network consists of a 1D convolution (kernel size *k*=8192, stride 1, no padding) followed by batch normalization and ReLU, and a 1D transpose convolution (kernel size *k*=8192, stride 1, no padding) followed by a second batch-normalization layer. A residual connection^29^ then adds the original logits back to the smoothed output, producing 𝑌_𝑠𝑚𝑜𝑜𝑡ℎ_ ∈ ℝ^𝐵×3×L^. The residual is applied only to the three logit channels, whereas the position channel is used only to guide smoothing. In our implementation, *k*=8192 spans two adjacent 4,096-SNP windows when *W*=4096, allowing the smoother to incorporate cross-window context during refinement. Under this convolution–transpose-convolution configuration, the final output length matches the input length while avoiding padding-induced boundary artifacts. Channel-wise softmax is applied downstream in the inference pipeline to obtain per-site probabilities.

### Phase-wise training

We train the system in two phases. Phase I (base model): the base network is optimized on simulated data using non-overlapping windows of *W*=4096 SNPs. Inputs comprise six reference channels (AFR/DEN/NEAN max/mean) plus a relative distance channel obtained by converting physical distance to Morgans and subtracting the within-window minimum; the model outputs per-site logits over non-archaic, Denisovan, Neanderthal. Phase II (smoother): the base model is frozen. We run overlapping windows with (*W*, *S*) = (4096, 512), reassemble per-window logits onto the global SNP coordinate axis by scatter-adding logits from all covering windows and dividing by the coverage count, and then train a 1D convolutional smoother on the reassembled logits. The smoother consumes [*B*, 3, *L*] logits concatenated with a Morgan-scaled absolute position channel [*B*, 1, *L*] and uses a 1D convolution (kernel size *k* = 8192, stride 1, no padding) with batch normalization and ReLU, followed by a 1D transpose convolution (kernel size *k* = 8192, stride 1, no padding) with batch normalization, and a residual addition to the three logit channels, yielding 𝑌_𝑠𝑚𝑜𝑜𝑡ℎ_ ∈ ℝ^𝐵×3×L^. The loss is computed on logits (prior to Softmax). Targets are canonicalized before loss computation: per-site labels are encoded as 𝑦 ∈ {0,1,2} for non-archaic, Denisovan and Neanderthal, respectively; in single-archaic settings, the unused archaic class is collapsed into the present one. For model selection we track validation *F1* and select the checkpoint achieving the best validation *F1*; training is terminated once validation *F1* does not improve for multiple consecutive epochs. All main results are reported with this two-phase regimen. A practical advantage of this design is memory efficiency: overlap reassembly is performed as a precomputed intermediate between phases (without backpropagation), so overlap-aware signals are retained while avoiding the substantial GPU-memory overhead that would arise if overlap aggregation were embedded inside end-to-end gradient updates.

### Data augmentation

We use two stochastic perturbations during training. (i) Genotype-flip noise: with probability 0.5 we flip allele states in both the query and reference channels at a maximum per-site rate of 0.02 to mimic sequencing/alignment errors. (ii) Offset jitter (Phase I): because the base model is trained on non-overlapping windows (no stride) whereas inference uses overlapping windows with (*W*, *S*) = (4096,512), we introduce a random offset before windowing to randomize the alignment between window boundaries and genomic coordinates. Concretely, we apply a random padding of up to *W* = 4096 SNPs prior to window extraction and remove this padding after prediction. This procedure exposes the base model to all window phases, encourages shift-robust per-site predictions, and mitigates train–test phase mismatch that would otherwise arise from the fixed stride used at inference. No stochastic augmentation is applied during Phase II (smoother training) or at inference.

### Optimization

We train the model in a supervised multi-class setting and optimize parameters using stochastic gradient-based methods^30^. To reduce the dominance of easy negatives under the strong imbalance between archaic and non-archaic sites, we use a three-class focal loss^31^. Let *y*_c_ denote the one-hot target for class *c*, and let *p_c_* denote the predicted probability for class *c* after softmax, with *C*=3. The loss is defined as

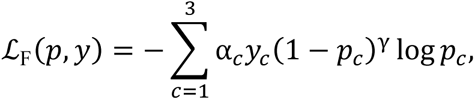

where we follow the original focal-loss experiments and use γ=2 and a uniform *α_c_*=0.25 for all three classes^31^. In this setting, the focusing effect is introduced primarily through the (1 − 𝑝_𝑐_)^𝛾^ term rather than through class-specific weighting. The loss is computed on logits (prior to softmax) using a numerically stable log-softmax formulation, consistent with the two-phase training regimen described above. Targets are canonicalized before loss computation as specified in the Phase-wise training section. Optimization uses Adam^32^ with learning rate 10^−3^ and ( 𝛽_1_, 𝛽_2_ )= (0.9,0.999) , together with a 400-step linear warm-up from 10^−5^ to the target rate, and learning-rate decay is disabled in the released configuration. Training uses batch size 1 and no gradient accumulation in the released settings. Training runs for up to 100 epochs; we select the checkpoint that achieves the best validation F1 for inference, and stop training once validation F1 shows no further improvement for multiple consecutive epochs. Implementation uses PyTorch 2.2^33^ and Python 3.8.

### Simulation framework and demographic models

Msprime 1.0^34,35^ was used to generate an extensive dataset for both training and evaluation, based on six demographic models (Fig. S2.1, Fig. S2.5, Tables S1–S6). The first three models represent single archaic introgression scenarios: a Bonobo–Ghost model^36^, a Human–Neanderthal model^16^, and a Human–Denisovan model^22^. The subsequent three models simulate dual archaic introgression scenarios: a Chimpanzee–Ghost–Bonobo model^21^, a Papuan–Neanderthal–Denisovan model^37,38^, and a Europe–Neanderthal–Denisovan model^23^.

### Generation, augmentation, and partitioning of simulated data

Training data were generated from all demographic models except the Human–Denisovan model. Parameters within some models were adjusted to broaden dataset diversity (Table S7), with unmentioned parameters defaulting to those specified in Tables S1–S6. For each parameter set within the training dataset, we performed ten replicate simulations and applied a per-replicate filter to the number of loci. Specifically, across the ten replicates we retained 0.10, 0.13, 0.17, 0.22, 0.28, 0.36, 0.46, 0.60, 0.77, 1.00 of the original number of loci, thereby simulating scenarios of missing loci. Unless specifically indicated, simulations assumed a constant recombination rate. When variability was required, we utilized the HapMap recombination maps for chromosome 22^39^ to ascertain recombination rates.

In total, 147,000 haplotypes of 50 Mb length were generated for the training dataset, with 80% allocated for training and 20% designated for validation. Furthermore, an additional set of 4,400 haplotypes, each 50 Mb in length and based on all demographic models above, was generated for use as a test dataset.

### Evaluation

#### Decoding and post-processing: from per-site posteriors to segments

The model outputs per-site posteriors over a class set that contains non-archaic and the archaic classes under consideration.

We first take the argmax class per site and retain sites labeled as an archaic class. For each haplotype, candidate segments are then constructed in a deterministic pass. A maximum gap *G* is applied so that archaic-labeled SNP clusters separated by a large physical interval are not treated as a single contiguous tract. This is particularly useful across marker-sparse or low-information intervals, including regions such as centromeres or other long genomic stretches with sparse retained archaic-labeled sites, where physically distant sites may otherwise become adjacent in the retained archaic-site list. The resulting archaic sub-blocks are scanned in genomic order, and adjacent sub-blocks separated by less than a user-specified merge distance *δ* are merged. The default value of *δ* was guided by simulation-based sensitivity analyses (Supplementary Information, Fig. S3.7), which evaluated archaic-tract detection performance across a range of merge distances and identified 10 kb as a stable working value under the setting considered here. Sub-blocks with fewer than two SNPs are discarded. Sensitivity analyses on simulated data showed that filtering short sub-blocks before merging helps control false positives. After merging, score–length filtering is applied using the default settings derived from the analyses in Fig. S3.7 and Fig. S3.8. Finally, source composition is annotated at the segment level after tract construction and post-processing. Mosaic is not a model output class during training; instead, it is used only as a post hoc annotation for merged segments that contain support from more than one archaic class. In this setting, a mosaic threshold *τ* determines whether the minority-source contribution is large enough for the segment to be reported as mosaic rather than assigned exclusively to the majority source. When support from only one archaic class is present, the segment is assigned directly to that class. The default hyperparameters and decoding thresholds used in AS3 are summarized in Table S15.

### Benchmarking on simulations

We evaluate tract-detection performance using length-weighted precision, recall and F1 computed from base-pair overlap between inferred and ground-truth archaic tracts; given the pronounced class imbalance, metrics are macro-averaged across archaic classes, and we additionally report an inferred/true length ratio to quantify over- or under-estimation. Beyond length-based metrics, we assess segment-level recovery by considering a ground-truth segment successfully recovered if at least 80% of its length is covered by any predicted segment. Parameter sensitivity analyses and robustness experiments (including SNP-density downsampling and variant-ascertainment filters) follow the Supplementary Information.

### Empirical concordance and computational profiling

To assess empirical concordance, we run ArchaicSeeker 3.0 on 1000 Genomes Phase 3 (KGP2504)^40,41^ non-African samples and compare results to representative published callsets after lightweight harmonization, reporting overlap-based concordance summaries. To evaluate additional AS3-specific introgressed segments, we further identified high-frequency segments detected only by AS3, filtered them by population frequency, source-specific archaic-reference match rate and the number of comparable archaic-informative variants, and assessed representative loci using haplotype-state and maximum-likelihood phylogenetic analyses. We also profile wall-clock runtime, peak RSS and peak GPU memory under varying cohort sizes, reference-panel sizes, SNP densities and genome lengths to characterize computational scaling; full protocols are provided in the Supplementary Information.

### Segment confidence

For a segment labeled as class 𝑐̂, confidence is defined as the mean posterior restricted to sites of that class within the segment:

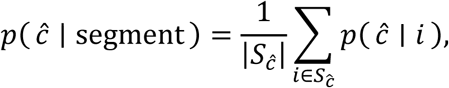

where 𝑆_𝑐̂_ = { 𝑖 ∈ segment: label(𝑖) = 𝑐̂ }.

### Precision, recall and macro-averages

Given the class imbalance, we report macro-averaged precision, recall, and 𝐹_1_ across the archaic classes. For class *c*,

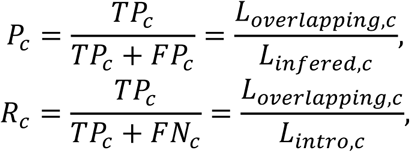

where 𝑇𝑃_𝑐_, 𝐹𝑃_𝑐_ and 𝐹𝑁_𝑐_ denote true positives, false positives, and false negatives for class 𝑐 ; 𝐿*_overlap_*_,𝑐_ is the total base-pair overlap between predicted and true segments of class 𝐿_𝑖𝑛𝑓𝑒𝑟𝑒𝑑,𝑐_ is the total predicted length; and 𝐿_𝑖𝑛𝑡𝑟𝑜,𝑐_ is the total ground-truth length. Macro-averaged metrics are the arithmetic mean across classes:

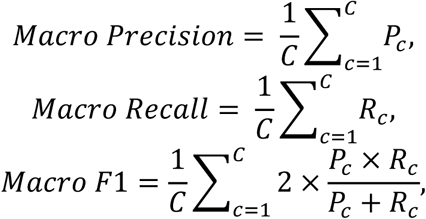

with 𝐶 the number of archaic classes considered (typically *C*=2 in dual-source settings). When only one archaic source is considered, source-wise macro-averaging is not applicable and the corresponding single-source precision, recall, and F1 are reported. For model selection we use validation *F1* (macro-averaged over archaic classes) and select the checkpoint with the best validation *F1* once it plateaus for multiple epochs. Primary reporting on simulated and real genomes is at the segment level, averaging metrics across chromosomes and samples.

### Empirical cohorts and preprocessing overview

We evaluated AS3 on phased modern human genomes together with high-coverage archaic references. The modern panel was assembled from the 1000 Genomes Project (KGP)^40,41^, the Human Genome Diversity Project (HGDP)^42^, the Simons Genome Diversity Project (SGDP)^43,44^, the Estonian Genome Diversity Panel (EGDP)^45^, and the Human Origins dataset (HO)^46,47^, yielding 3,453 individuals from 209 populations after harmonization (Table S14). Across cohorts, variants were lifted/converted to a unified reference build, restricted to high-confidence biallelic SNPs, and subjected to consistent quality control and masking prior to inference. To ensure cross-dataset comparability, we applied a uniform processing protocol for sample inclusion, SNP-level filtering, phasing/imputation harmonization, and archaic-reference matching before running AS3. This design enabled joint landscape analysis across both densely sampled and sparsely sampled populations without population-specific parameter fitting. Full details of reference design, archaic genome preprocessing, dataset-specific QC rules, and final inclusion masks are provided in the Supplementary Information.

## Results

### Overview of the ArchaicSeeker 3.0 framework

ArchaicSeeker 3.0 (AS3) is a two-stage neural framework for haplotype-resolved archaic introgression inference that separates within-window evidence extraction from chromosome-scale tract reconstruction (Fig. 1). Rather than treating tiled window predictions as final outputs, AS3 first generates site-wise ancestry logits from long-context SNP windows and then reconciles overlapping predictions into a chromosome-aligned logit track for downstream boundary refinement and tract decoding. This design decouples local sequence modeling from cross-window consolidation, providing a coherent basis for haplotype-level segment inference across the chromosome.

Given phased target haplotypes together with African outgroup, Denisovan and Neanderthal references, AS3 constructs a constant-width per-site representation that compresses reference information and appends a distance/position-derived channel (Fig. 1b; Methods). This compact encoding allows the model to ingest heterogeneous reference panels while keeping the input dimensionality fixed, enabling consistent training and deployment across datasets.

The Base model applies a long-context backbone based on selective state-space modeling (Mamba-2) to fixed-length windows of W = 4,096 SNPs, producing per-site logits over three ancestry states (non-archaic, Denisovan and Neanderthal) for each window (Fig. 1b; Methods). In contrast to attention-based backbones with quadratic scaling in window length, the state-space formulation scales approximately linearly, making it feasible to model multi-kilobase SNP contexts under standard hardware budgets while preserving site-wise outputs required for downstream refinement.

To obtain chromosome-wide predictions, AS3 runs the Base model in an overlapping sliding-window mode at inference time and reassembles a single contiguous logit track aligned to the original SNP axis by aggregating predictions from overlapping windows (Fig. 1a; Methods). Using an overlap stride of S = 512 SNPs, this procedure mitigates discontinuities at window edges and provides a globally consistent per-site representation for refinement.

AS3 then applies a lightweight residual 1D convolutional Smoother to enforce cross-window boundary coherence on the reassembled logit track (Fig. 1c). The Smoother operates on the reassembled logits augmented with a position/distance channel and uses a wide effective receptive field (default *k* = 8,192) to integrate context spanning adjacent windows, yielding refined per-site logits from which posterior probabilities are obtained via Softmax and discrete labels via argmax (Methods). Finally, haplotype-resolved archaic tracts are derived downstream by converting per-site labels into contiguous segments under gap/merge/mosaic criteria.

Training follows the same two-stage structure as the inference pipeline (Fig. 1a): Phase I trains the Base model on simulations spanning diverse demographic scenarios to promote robustness to demographic misspecification; Phase II trains the Smoother using reassembled logits generated by overlapping inference. Once trained, AS3 can be applied to new cohorts without cohort-specific demographic parameter estimation or re-fitting, supporting deployment in settings with limited target sample sizes while maintaining tract-level boundary stability.

### Generalizes across heterogeneous demographic histories without inference-time demographic fitting

A central obstacle in archaic local ancestry inference is demographic model misspecification: methods with explicit demographic assumptions or cohort-level parameter estimation can degrade when the true history deviates from the assumed model, particularly under multi-pulse events, mixed archaic sources, or continuous gene flow. To stress-test robustness under diverse and intentionally heterogeneous histories, we benchmarked ArchaicSeeker 3.0 (AS3) against representative baselines (ArchaicSeeker 2.0, IBDmix, hmmix, DAIseg and Sprime) on an independent msprime-based simulation suite spanning single- and dual-lineage introgression and a range of temporal modes (single pulse, multiple pulses and continuous gene flow). Performance was evaluated at the segment level based on base-pair overlap between predicted and ground-truth tracts; given the pronounced class imbalance between introgressed and non-introgressed sequence, we report macro-averaged precision, recall and F1 across archaic classes. This benchmark was designed not simply to compare callers under matched conditions, but to test whether a simulation-trained sequence model could retain stable performance when applied across demographically heterogeneous scenarios without inference-time cohort-specific fitting or recalibration.

Across target/reference size configurations, AS3 consistently occupies a favorable region of the precision–recall plane, maintaining high precision while avoiding the pronounced recall loss observed for more conservative callers (Fig. 2a). ArchaicSeeker 2.0 typically operates at a conservative point (higher precision but reduced recall), whereas Sprime shows the opposite trade-off (higher recall with reduced precision). Other baselines display larger dispersion across settings, indicating greater sensitivity to sample-size configuration and demographic complexity (Fig. 2a). Stratifying by demographic scenario further shows that AS3 attains the strongest archaic-seeking F1 across all tested histories, including challenging settings such as multi-pulse events and mixed-ancestry configurations (Fig. 2b; Fig. S2.2–S2.4). Together, these results indicate that AS3’s window-based multi-locus modeling, trained on diverse simulated histories, maintains stable detection accuracy under demographic heterogeneity rather than being restricted to a narrow family of matched training scenarios. More broadly, they suggest that simulation-trained sequence inference can generalize across demographic mismatch without requiring the training distribution to exhaustively enumerate all evaluation-time models (Fig. 2b; Fig. S2.7).

**Fig. 2.**
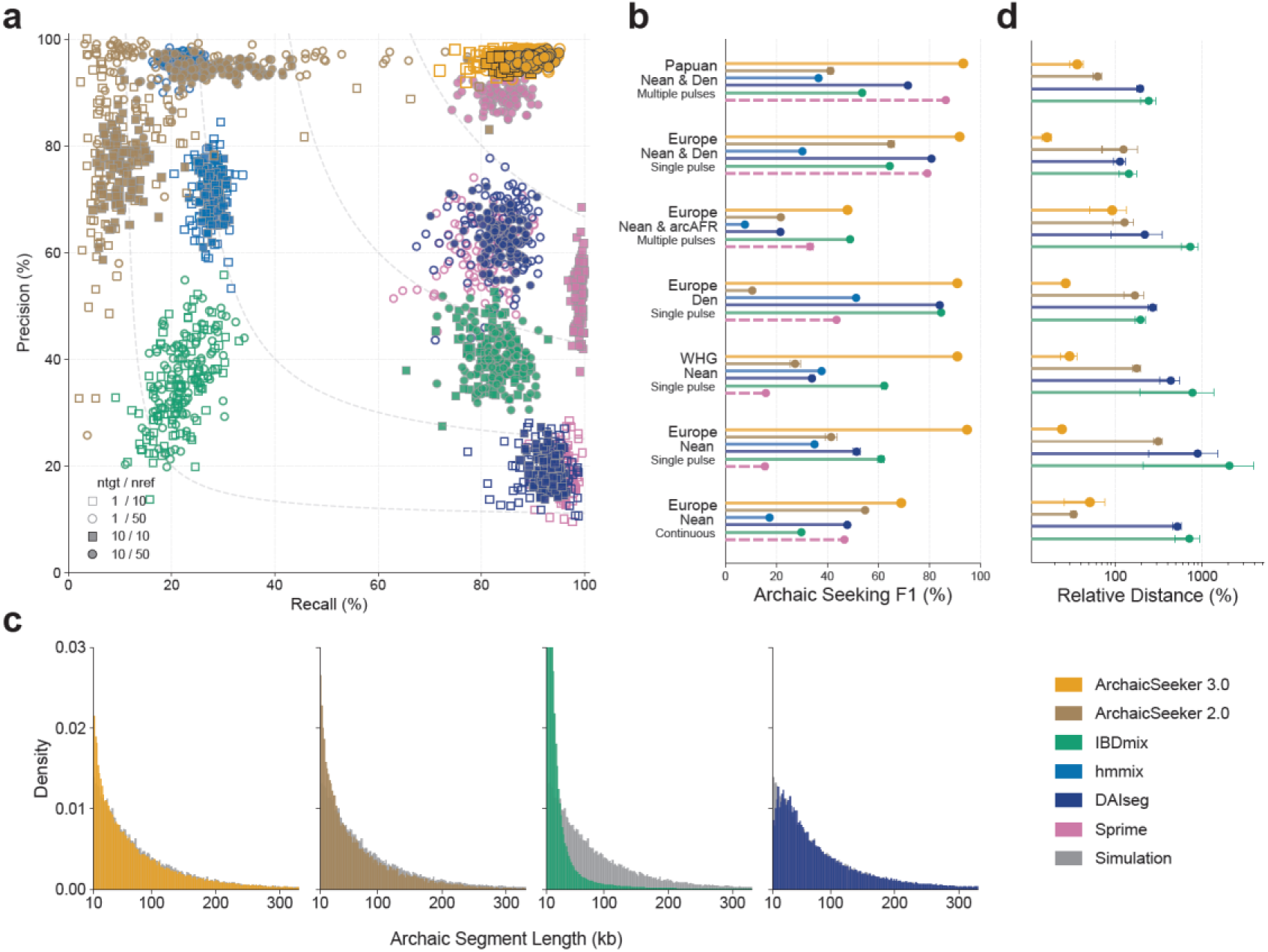
Simulation benchmarks of archaic introgression detection. (a) Segment-level precision versus recall across simulation settings. Marker shape encodes target/reference sample sizes (*n_tgt_*/*n_ref_*): open square, 1/10; open circle, 1/50; filled square, 10/10; filled circle, 10/50. Grey dashed curves indicate iso-F1 contours. (b) Archaic-seeking F1 (%) across demographic scenarios: Papuan Nean & Den (multiple pulses); Europe Nean & Den (single pulse); Europe Nean & arcAFE (multiple pulses); Europe Den (single pulse); WHG Nean (single pulse); Europe Nean (single pulse); Europe Nean (continuous). Points summarize performance across replicates; error bars indicate variability across replicates (see Methods). (c) Distributions of inferred archaic segment lengths (kb) for methods that directly output haplotype-level tracts, compared with the simulated truth (grey). hmmix and Sprime are not shown in this panel because they do not yield tract lengths in a directly comparable format: Sprime reports introgressed regions rather than haplotype-resolved tracts, and hmmix outputs calls in fixed 1,000-bp windows that require additional post-processing to derive contiguous segments. (d) Boundary accuracy measured by relative distance (%) between inferred and true segment boundaries (log-scaled x-axis; see Methods). Points summarize performance across replicates; error bars indicate variability across replicates.

AS3’s robustness is also reflected in tract-level call structure. The inferred segment-length spectrum more closely follows the ground-truth distribution, consistent with reduced fragmentation into short segments and improved tract coherence (Fig. 2c; Fig. S2.6). Segment-length distributions are shown only for methods that natively output haplotype-resolved contiguous tracts; Sprime reports region-level candidates rather than haplotype-resolved tracts, and hmmix outputs fixed 1-kb window calls that require additional, user-defined merging to derive tract lengths (Methods), precluding a direct like-for-like comparison in this panel. Because tract length distributions encode information about the timing and structure of introgression, improved tract coherence and boundary fidelity are relevant not only for detection accuracy but also for downstream historical interpretation.

### Training diversity, rather than raw sample volume, drives cross-model generalization

To verify that the observed robustness stems from our training strategy, we conducted a systematic ablation study on training data composition (Table S8). We hypothesized that the diversity of evolutionary histories seen during training is critical for generalization. To test this, we trained variants of the model on subsets of data representing varying degrees of demographic complexity while keeping the validation set constant. This analysis was intended to distinguish two possible sources of robustness: increased exposure to more training haplotypes within a single model, versus broader exposure to qualitatively different introgression histories.

We observed that while increasing sample size within a single demographic model provided marginal gains, the inclusion of diverse demographic scenarios was the primary determinant of out-of-distribution performance. Models trained on a limited variety of demographic histories showed substantially weaker generalization to held-out demographic settings, regardless of the total number of training haplotypes. Conversely, training with a strategically diverse regimen—spanning both single- and multi-archaic introgression pulses—allowed the model to learn invariant features of introgression, stabilizing performance across distinct demographic models (Table S8). These findings indicate that, for simulation-trained population-genetic inference, the breadth of evolutionary regimes represented in the training simulations is a more important determinant of robustness than raw sample volume within any single demographic model. In this setting, training diversity appears to matter not simply because it increases data quantity, but because it exposes the model to a wider range of tract-generating processes from which more transferable sequence-level decision rules can be learned.

### Accurate boundary localization supports tract-scale historical interpretation

Beyond overall detection, boundary placement is critical because tract boundaries encode recombination breakpoints and therefore constrain inference on introgression timing and downstream locus-level interpretation. We quantified boundary accuracy using the relative-distance metric, measuring the deviation between inferred and true tract boundaries (smaller values indicate more accurate localization; Fig. 2d; Supplementary Information). Across demographic scenarios, AS3 yields consistently smaller relative distances than competing methods, indicating tighter localization of both start and end coordinates and reduced boundary inflation/overextension (Fig. 2d). In contrast, several baselines exhibit scenario-dependent boundary errors—either over-merging nearby signals into longer tracts or producing irregular boundaries that drift from true breakpoints—leading to substantially larger relative distances. Together, these results show that AS3 improves detection performance while simultaneously enhancing tract coherence and boundary precision across heterogeneous introgression histories.

### Haplotype-wise inference enables few-target analyses

A practical limitation of cohort-based archaic LAI approaches is their reliance on sufficiently large target cohorts to stably estimate cohort-specific parameters (e.g., transition rates or demographic summaries), which can be prohibitive for under-sampled populations and study designs constrained by sequencing budgets. In contrast, ArchaicSeeker 3.0 performs inference per haplotype using a fixed, learned decision rule and does not require cohort-level parameter fitting or re-calibration at inference time. This design avoids failure modes induced by unstable cohort-level estimates and makes the method applicable when only a small number of target individuals are available.

Consistent with this design, AS3 remains effective in a small-target regime. We evaluated performance as a function of the number of target haplotypes available at inference and found that both archaic-seeking *F1* and archaic-source classification accuracy remain stable across target cohort sizes, including when *n_tgt_* ≤ 10 (Fig. 3a; Fig. S3.1). These results indicate that AS3’s reference-compressed representation and window-based inference do not depend on large target cohorts for calibration, enabling robust tract inference for rare populations, pilot datasets, or scenarios where only a handful of individuals can be deeply sequenced or confidently phased.

**Fig. 3.**
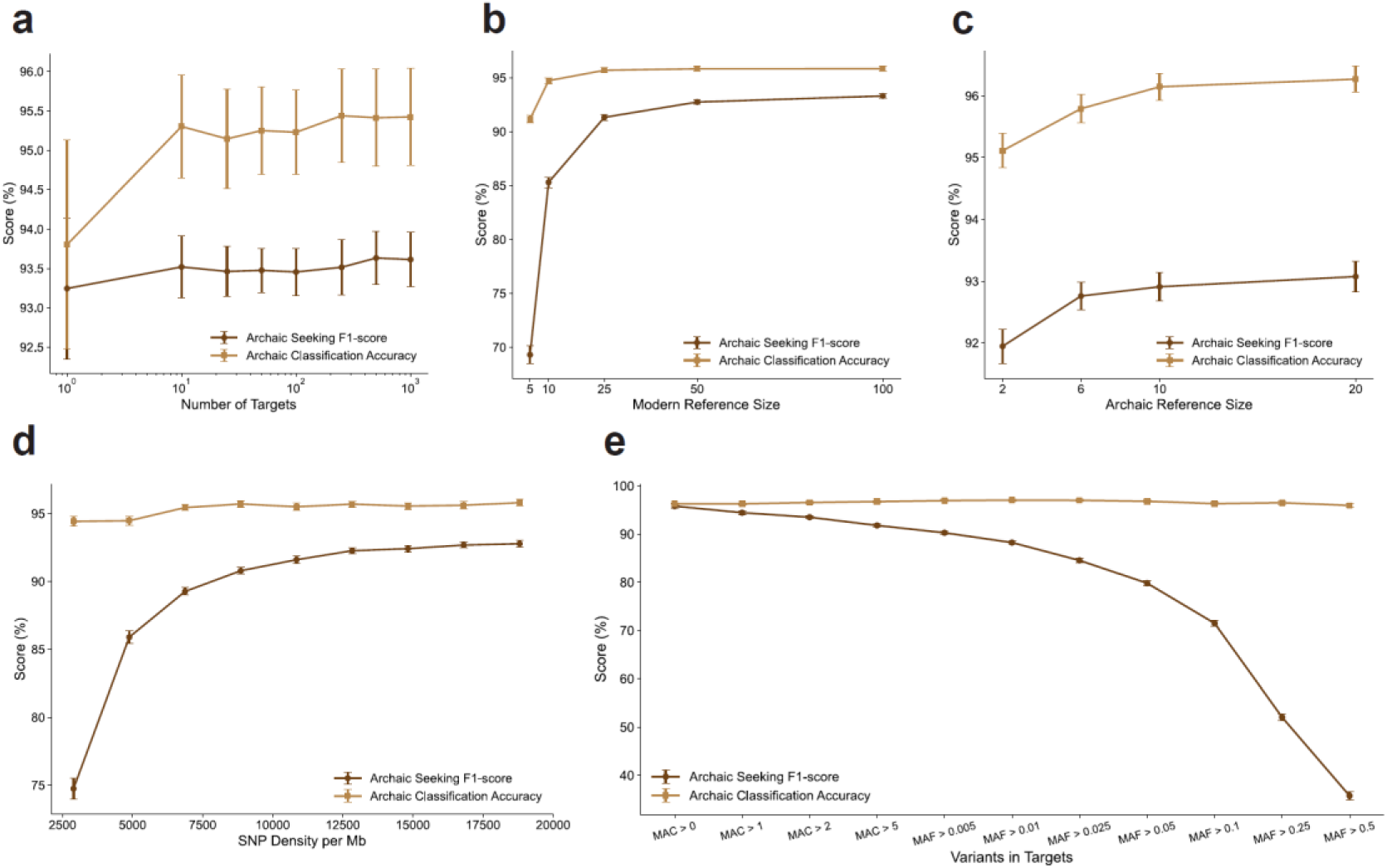
Operating regimes of ArchaicSeeker 3.0 across cohort size, reference information and marker properties. (a) Archaic-seeking *F1*-score and archaic classification accuracy as a function of the number of target haplotypes (log-scaled x-axis). (b) Archaic-seeking *F1*-score and archaic classification accuracy as a function of modern (African) reference panel size. (c) Archaic-seeking *F1*-score and archaic classification accuracy as a function of archaic reference panel size. (d) Archaic-seeking *F1*-score and archaic classification accuracy as a function of SNP density (per Mb) in the target data. (e) Archaic-seeking *F1*-score and archaic classification accuracy under increasingly stringent variant ascertainment in targets, using minor allele count (MAC) or minor allele frequency (MAF) thresholds as indicated. Scores are reported as percentages. Points summarize performance across replicates; error bars indicate variability across replicates (see Methods). Archaic-seeking F1 is computed at the segment level and macro-averaged across archaic classes; archaic classification accuracy reflects correct assignment of archaic source labels (Denisovan versus Neanderthal) (see Methods).

### Reference information requirements

To clarify how much reference information is required for stable tract calling, we varied the sizes of (i) the modern outgroup panel (Sub-Saharan African haplotypes) and (ii) the archaic reference panels while holding the inference pipeline fixed.

Increasing the modern (African) reference size produced the largest gains when the panel was extremely small: archaic-seeking F1 improved sharply from minimal panel sizes and then approached a plateau as additional African haplotypes were added (Fig. 3b; Fig. S3.2). In contrast, expanding the archaic reference size yielded a more modest but consistent improvement that saturated quickly, with diminishing returns beyond intermediate panel sizes (Fig. 3c; Fig. S3.3). Across these settings, archaic-source classification accuracy remained comparatively stable, whereas archaic-seeking *F1* was more sensitive to the amount of reference information available (Fig. 3b–c; Fig. S3.4). Together, these results indicate that AS3 does not require large archaic panels to function reliably, while avoiding overly small modern outgroup panels is more important for sensitive detection.

### Post-processing parameterization and operating-point trade-offs

We next optimized the post-processing stage used to convert window-level probabilities into tract calls. Before merging, these candidates were well separated by both score and length: overlap-positive windows were enriched at high scores (∼0.85–0.95), whereas overlap-negative windows concentrated around lower scores (∼0.40–0.65) and short segments (<10 kb) (Fig. S3.7a, b, e). A grid search identified a broad optimum at pre-merge minimum length ∼5 kb and merge distance ∼8–12 kb (peak archaic-seeking F1 ∼92.8–92.9%); compared with merge-first decoding, pre-filtering before merge increased median F1 (∼92.8% vs ∼91.6%) and precision (∼95.3% vs ∼90.5%) (Fig. S3.7c, d, f–h). Under post-merge filtering, F1 remained near plateau (∼92.7–92.9%) for min score 0.4–0.7 when minimum length was ≤10 kb, while stricter length thresholds increased precision (to ∼97%) and segment precision (to ∼99–100%) at the cost of recall (∼91% to ∼79%) and inferred archaic ratio (∼97% to ∼82%); source-classification accuracy varied within a narrower band (∼95.0–97.1%) (Fig. S3.8).

### Robustness to marker properties and recombination features

We next examined robustness to practical properties of the target data that can act as information bottlenecks for tract discovery, focusing on marker density, variant ascertainment, and recombination-related feature construction. Reducing SNP density led to a pronounced decline in archaic-seeking F1 (Fig. 3d), consistent with a broader loss of informative variation and local haplotypic patterns that support tract detection and boundary localization; in contrast, archaic-source classification accuracy changed relatively little across densities, suggesting that when segments are detected, source discrimination remains stable.

Variant ascertainment had an even stronger impact on detection: progressively filtering the target data to retain only common variants caused a marked deterioration of archaic-seeking F1, while classification accuracy remained largely stable (Fig. 3e; Fig. S3.6). This pattern indicates that low-frequency and private-allele information in the target haplotypes contributes disproportionately to identifying introgressed tracts, whereas the Denisovan-versus-Neanderthal label is comparatively robust once an archaic segment is supported. As an orthogonal ascertainment analysis, we also applied allele-frequency filtering to variants in the modern reference panel, and observed corresponding shifts in tract-detection and source-assignment metrics (Fig. S3.5).

Finally, we evaluated sensitivity to recombination-related choices used to construct distance/position features. Using a constant average recombination rate to convert physical distance to Morgans performed comparably to, and in this benchmark no worse than, supplying a fine-scale rate map (Fig. S3.9a). Likewise, using genetic distance versus physical distance for the distance/position features did not materially change performance (Fig. S3.9b). These results support a practical operating regime in which AS3 remains reliable without requiring high-resolution genetic maps, but benefits strongly from maintaining sufficient marker density and retaining informative variation (including low-frequency sites) for sensitive tract discovery.

### Computational efficiency, usability and biobank-scale feasibility

We profiled inference-time computational scaling of ArchaicSeeker 3.0 (AS3) across cohort size, reference-panel size, marker density and genomic span (Table 1). AS3 supports high-throughput batched inference with a compact memory footprint: runtime per haplotype decreases and then saturates with increasing batch size, consistent with improved accelerator utilization under batched execution, while peak GPU memory and peak host RSS remain within a few GiB for typical chunk sizes (Table 1). Operationally, AS3 is designed to be simple to run with a minimal input contract: inference requires only phased, haplotype-resolved VCFs for (i) the Phased Target cohort and (ii) a phased reference set comprising sub-Saharan African haplotypes and archaic haplotypes (or pseudo-haplotypes, when applicable). Phasing is required because AS3 operates at haplotype resolution, and the two haplotypes of each diploid individual are treated as independent inference units, enabling straightforward batching and parallelization. This input design supports a practical deployment pattern in which genomes are processed as independent genomic blocks (for example, per chromosome or fixed Mb chunks) while target haplotypes are batched to maximize throughput, making end-to-end wall time largely governed by job granularity, I/O bandwidth and scheduler availability rather than algorithmic constraints.

**Table 1.** Computational efficiency and resource usage of ArchaicSeeker 3.0 under varying dataset configurations. Performance metrics (Wall-clock time, Peak RSS, and Peak GPU memory) were measured on an NVIDIA 4090 GPU. Values represent the mean ± s.d. over 10 independent runs. Unless variable, fixed parameters were set to: Target size n=10 haplotypes (except n=1 for Genome Length experiments), Reference size m=50, Sequence length L=10 Mb, and SNP density ∼ 19,500 SNPs/Mb (except ∼ 13,300 SNPs/Mb for Genome Length experiments).

| Variable |  | Wall-clock Time (s) | Peak RSS (GiB) | Peak GPU Memory Usage (GiB) |
| --- | --- | --- | --- | --- |
| # Target | 1 | 17.37 ± 0.26 | 1.46 ± 0.00 | 2.70 ± 0.17 |
|  | 10 | 52.81 ± 0.54 | 1.50 ± 0.00 | 3.03 ± 0.02 |
|  | 25 | 111.70 ± 1.28 | 1.52 ± 0.00 | 3.06 ± 0.01 |
|  | 50 | 210.40 ± 2.54 | 1.55 ± 0.00 | 3.06 ± 0.00 |
|  | 100 | 408.27 ± 5.07 | 1.61 ± 0.01 | 3.07 ± 0.00 |
|  | 250 | 1006.53 ± 12.70 | 1.79 ± 0.03 | 3.07 ± 0.00 |
|  | 500 | 2003.16 ± 29.95 | 2.10 ± 0.05 | 3.07 ± 0.01 |
|  | 1000 | 4019.53 ± 54.68 | 2.70 ± 0.10 | 3.07 ± 0.01 |
| # Reference | 5 | 53.13 ± 0.28 | 1.32 ± 0.00 | 2.93 ± 0.00 |
|  | 10 | 50.78 ± 0.26 | 1.32 ± 0.00 | 2.90 ± 0.00 |
|  | 25 | 50.86 ± 0.31 | 1.38 ± 0.00 | 2.94 ± 0.00 |
|  | 50 | 53.11 ± 0.29 | 1.50 ± 0.00 | 3.02 ± 0.02 |
|  | 100 | 59.03 ± 0.34 | 1.86 ± 0.00 | 3.08 ± 0.04 |
| SNP Density per Mb | 1950 ± 4 | 18.54 ± 0.51 | 1.06 ± 0.00 | 1.76 ± 0.11 |
|  | 3896 ± 8 | 25.61 ± 0.52 | 1.11 ± 0.00 | 2.66 ± 0.05 |
|  | 5849 ± 11 | 28.50 ± 0.29 | 1.16 ± 0.00 | 2.62 ± 0.06 |
|  | 7791 ± 10 | 33.06 ± 0.26 | 1.19 ± 0.00 | 2.72 ± 0.05 |
|  | 9758 ± 12 | 35.92 ± 0.29 | 1.26 ± 0.00 | 2.80 ± 0.05 |
|  | 11707 ± 18 | 38.88 ± 0.40 | 1.30 ± 0.00 | 2.77 ± 0.08 |
|  | 13648 ± 18 | 42.69 ± 0.27 | 1.33 ± 0.00 | 2.83 ± 0.05 |
|  | 15614 ± 19 | 45.96 ± 0.28 | 1.36 ± 0.00 | 2.94 ± 0.04 |
|  | 17548 ± 21 | 49.73 ± 0.32 | 1.47 ± 0.00 | 2.99 ± 0.02 |
|  | 19505 ± 24 | 52.95 ± 0.32 | 1.50 ± 0.00 | 3.02 ± 0.02 |
| Genome Length (Mb) | 10 | 15.30 ± 0.20 | 1.31 ± 0.00 | 2.32 ± 0.03 |
|  | 20 | 19.00 ± 0.16 | 1.67 ± 0.00 | 2.52 ± 0.08 |
|  | 50 | 30.51 ± 0.23 | 2.59 ± 0.01 | 3.33 ± 0.12 |
|  | 75 | 41.33 ± 0.35 | 3.52 ± 0.00 | 3.66 ± 0.04 |
|  | 100 | 51.60 ± 0.31 | 4.21 ± 0.01 | 4.18 ± 0.12 |
|  | 150 | 70.43 ± 1.10 | 6.08 ± 0.00 | 4.96 ± 0.08 |
|  | 200 | 87.79 ± 0.48 | 7.50 ± 0.00 | 6.50 ± 0.03 |
|  | 250 | 107.65 ± 0.60 | 9.90 ± 0.00 | 8.00 ± 0.00 |

### Projected compute requirements for UK Biobank–scale cohorts

Using the throughput measured in Table 1, we extrapolated the inference-only compute required for UK Biobank–scale^48^ processing (Table 2). Assuming ∼3,000 Mb of autosomal sequence per individual and haplotype-resolved inference, a cohort of 500,000 individuals (1,000,000 haplotypes) corresponds to ∼3×10^9 Mb of haplotype sequence. Under marker densities comparable to dense variant panels, the projected compute is ∼3.35×10^5 GPU-hours for the full cohort (Table 2), corresponding to ∼14 days on 1,000 GPUs, ∼28 days on 500 GPUs or ∼140 days on 100 GPUs under idealized full parallelism. At lower marker density, projected compute decreases proportionally (Table 2). These results indicate that AS3 is scalable to biobank-scale studies in large computing environments, with turnaround time determined primarily by available accelerator parallelism rather than changes to the core inference procedure.

**Table 2.** Projected inference-only compute for UK Biobank–scale processing based on Table 1 throughput. Estimates assume a cohort of 500,000 individuals (1,000,000 haplotypes). Assumptions: ∼3,000 Mb autosomal span per individual; haplotype-resolved inference; batched inference; estimates exclude preprocessing and I/O; realized wall time depends on job granularity, I/O bandwidth, and scheduler/queue dynamics. Compute derivation: Total GPU-hours = (runtime per haplotype per 10 Mb from Table 1) × (3,000 Mb / 10 Mb) × (1,000,000 haplotypes), converted to hours.

| SNP Density<br>(per Mb) | Runtime<br>(min/haplotype) | Total Effort<br>(GPU-hours) | Projected Turnaround Time<br>(days) |  |  |
| --- | --- | --- | --- | --- | --- |
|  |  |  | 100 GPUs | 500 GPUs | 1,000 GPUs |
| ~19,505 | ~20.10 | ~335,000 | ~139.6 | ~27.9 | ~14.0 |
| ~6,713 | ~13.04 | ~217,000 | ~90.5 | ~18.1 | ~9.1 |
| ~1,950 | ~9.27 | ~154,500 | ~64.4 | ~12.9 | ~6.4 |

### Orthogonal concordance supports the reliability of ArchaicSeeker 3.0

We evaluated the concordance of ArchaicSeeker 3.0 (AS3) with representative introgression callers on 1000 Genomes European samples (CEU, TSI, FIN, GBR and IBS), including ArchaicSeeker 2.0(AS2)^23^, hmmix^22^, IBDmix^24^, and Sprime^19^(Fig. 4). Across populations, AS3 identifies substantial archaic sequence, and a large proportion of the inferred tracts is supported by at least one other method; a consistent core is supported by two or more independent callers (Fig. 4a).

**Fig. 4.**
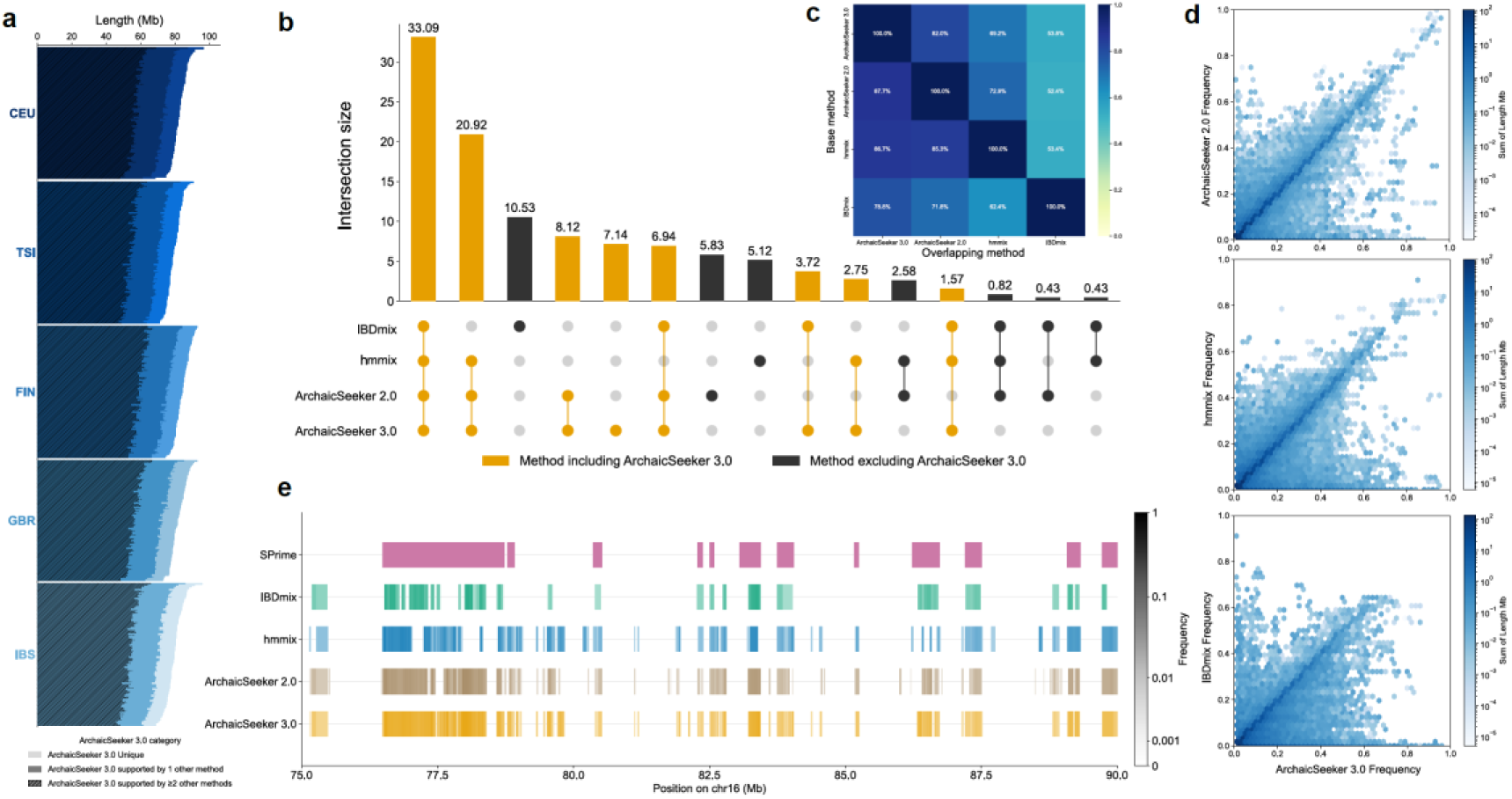
Empirical concordance of ArchaicSeeker 3.0 with established introgression callers. (a) Per-individual inferred archaic sequence length across 1000 Genomes European populations (CEU, TSI, FIN, GBR and IBS). AS3 calls are stratified by support level: AS3-unique, supported by one other method, or supported by two or more other methods; individuals are ordered by total inferred length within each population. (b) UpSet plot summarizing intersections of inferred archaic sequence among ArchaicSeeker 3.0, ArchaicSeeker 2.0, hmmix and IBDmix. Bars indicate intersection size (Mb); bar color distinguishes intersections that include AS3 from those that do not. (c) Pairwise overlap matrix among callers. Each cell reports the fraction of the base method’s inferred sequence that is overlapped by another method. (d) Concordance of segment frequency estimates. Frequencies inferred by ArchaicSeeker 2.0, hmmix and IBDmix are plotted against AS3 frequency; point density reflects the summed segment length per frequency bin. (e) Example region on chromosome 16 (75–90 Mb) comparing calls from Sprime and segment callers. Tracks are colored by segment frequency; AS3 tracts are annotated by support category, illustrating shared intervals and additional AS3-resolved candidates.

Intersection analysis shows that AS3 retains the shared signal while expanding the call set. The largest intersection (33.09 Mb) is jointly detected by AS3, AS2, hmmix and IBDmix, and an additional major component (20.92 Mb) is shared among AS3, AS2 and hmmix (Fig. 4b). Beyond these, AS3 contributes substantial sequence supported by subsets of baselines (for example, AS3+AS2, 8.12 Mb; AS3+AS2+IBDmix, 6.94 Mb), and also includes an AS3-specific component (7.14 Mb) that is comparable in magnitude to method-specific components observed for other callers (for example, IBDmix-only 10.53 Mb; AS2-only 5.83 Mb; hmmix-only 5.12 Mb) (Fig. 4b).

Pairwise overlap quantification further indicates high concordance between AS3 and established callers while preserving additional sensitivity. Using the fraction of the base method’s sequence overlapped by another method, AS3 overlaps most of the sequence inferred by AS2 (87.7%), hmmix (86.7%) and IBDmix (78.8%); in the reverse direction, AS2 and hmmix overlap 82.0% and 69.2% of AS3 calls, respectively, and IBDmix overlaps 53.8% (Fig. 4c). At the population level, AS3 segment-frequency estimates are highly concordant with those from AS2 and hmmix, and remain positively associated with IBDmix despite greater dispersion (Fig. 4d). A representative region on chromosome 16 further illustrates that AS3 recapitulates major candidate intervals highlighted by Sprime and other callers, while additionally resolving smaller or partially supported tracts (Fig. 4e). Together, these results support the reliability of AS3 on empirical genomes and its improved sensitivity relative to existing approaches.

To evaluate real-data behavior in a Denisovan-enriched population, we performed a Papuan-focused cross-method comparison. Frequency–frequency maps showed consistent diagonal enrichment for AS3 versus both AS2 and hmmix, indicating concordant population-level Denisovan frequency structure across methods (Fig. S4.2 and S4.3). Directed overlap was substantial but asymmetric: 57.8% of AS2 calls and 62.4% of hmmix calls were recovered by AS3, whereas 25.8% of AS3 calls overlapped AS2 and 36.5% overlapped hmmix; reciprocal overlap between AS2 and hmmix was 54.3% and 41.4%, respectively (Supplementary Fig. S4.4). UpSet decomposition further quantified total inferred Denisovan sequence of 84.56 Mb (AS3), 70.99 Mb (hmmix), and 47.28 Mb (AS2), including a 20.44 Mb three-way core and method-specific components (AS3-only, 25.94 Mb; hmmix-only, 26.44 Mb; AS2-only, 7.63 Mb) (Fig. S4.5). Together with the observed segment-length spread in Papuans (Fig. S4.1), these results support a shared Denisovan backbone across methods while indicating substantial non-redundant signal captured by AS3.

### Global landscape of archaic introgression in present-day populations

To systematically characterize archaic introgression at worldwide scale, we integrated modern genomes from KGP^40,41^, HGDP^42^, SGDP^43,44^, EGDP^45^ and HO^46,47^, yielding 3,453 individuals from 209 populations (Table S14). Because AS3 performs haplotype-level inference without population-specific parameter fitting, we were able to analyze both large cohorts and sparsely sampled populations within a unified framework.

Global mapping revealed distinct large-scale patterns for Neanderthal and Denisovan ancestry (Fig. 5a, b). Inferred Neanderthal ancestry proportions were comparatively constrained across populations (0.88–1.37%), whereas Denisovan proportions showed substantially broader heterogeneity (0.05–1.24%), with the highest levels concentrated in Oceania-related groups. These patterns indicate a widely shared Neanderthal background together with more geographically localized Denisovan enrichment.

**Fig. 5.**
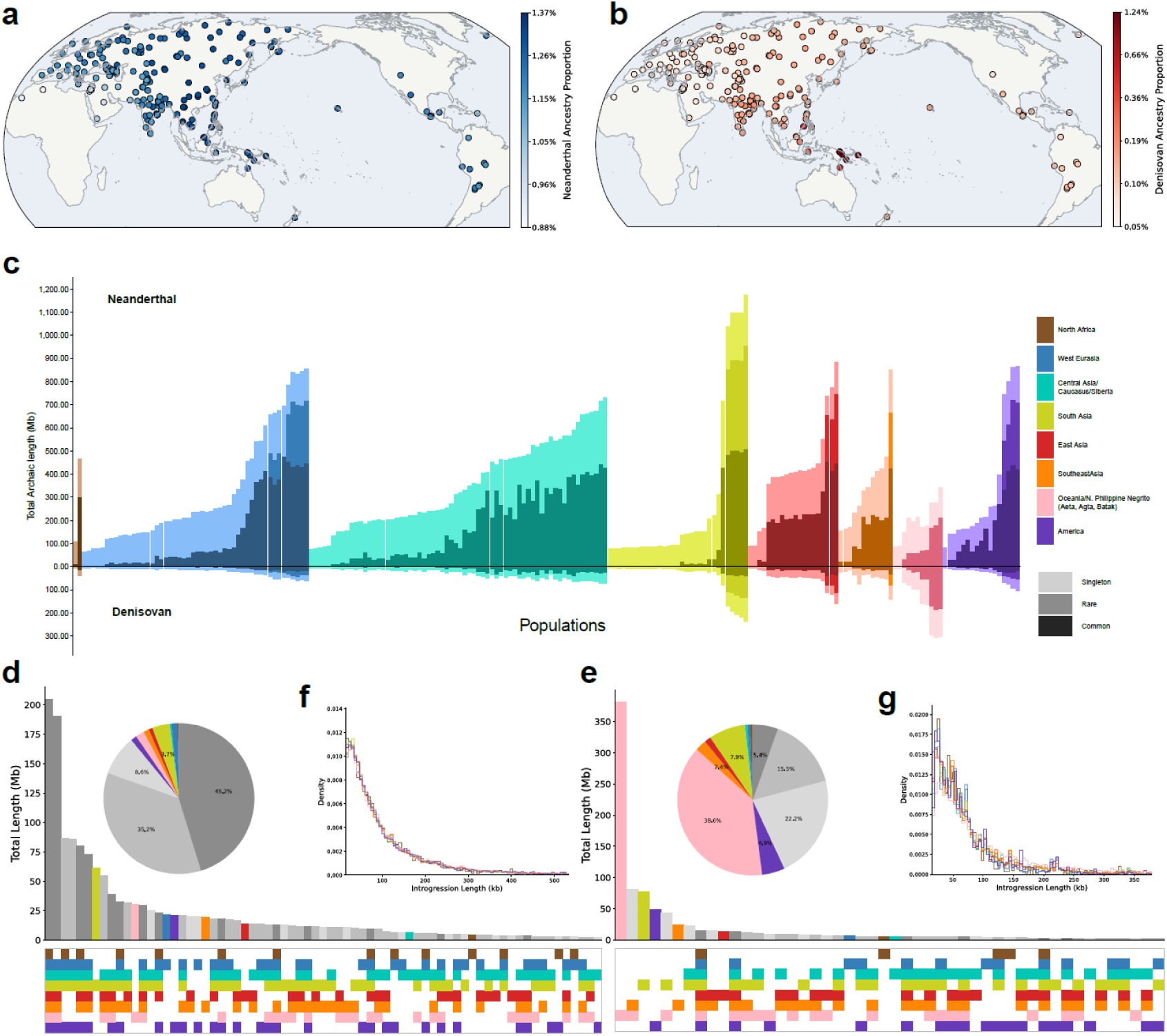
Global landscape of archaic introgression across present-day human populations. (a, b) Geographic distribution of inferred Neanderthal (a) and Denisovan (b) ancestry proportions across populations in the integrated modern dataset. (c) Population-level total inferred archaic sequence length (Mb), shown separately for Neanderthal and Denisovan and decomposed by frequency class (common, rare and singleton), with populations grouped by geographic region. (d, e) UpSet analysis of cross-region overlap structure for inferred Neanderthal (d) and Denisovan (e) introgressed segments. Vertical bars denote total sequence length (Mb) for each region-intersection pattern, and the connected matrix indicates region membership of each intersection. Insets summarize the relative contributions of major intersection classes. (f, g) Density distributions of inferred introgressed tract lengths for Neanderthal (f) and Denisovan (g) across regions.

At the population level, total inferred archaic sequence length and its frequency-stratified components further resolved this contrast (Fig. 5c). We then used UpSet analysis to quantify cross-region sharing architecture of inferred introgressed segments. Neanderthal and Denisovan signals showed different intersection structures across regions, including both broadly shared components and region-enriched subsets (Fig. 5d, e), consistent with differences in admixture history and subsequent demographic reshaping.

Finally, tract-length density profiles showed enrichment of short fragments with progressively decreasing density toward longer tracts for both sources (Fig. 5f, g). Region-specific differences in the long-tract tail support heterogeneous post-admixture histories superimposed on a shared recombination-driven fragmentation process.

### High-frequency AS3-specific introgressed segments show locus-level evidence of archaic introgression

Because cross-method comparisons indicated that AS3 captures a non-redundant component of archaic introgression signal, we next examined high-frequency AS3-specific introgressed segments at the locus level. Among all AS3-specific Neanderthal-introgressed segments, the most prevalent segment was located at chr20:63,697,389–63,769,790, spanning approximately 72.4 kb and reaching high frequency in East Asian populations, with frequencies of 61.1–68.0% across CDX, CHB, CHS, JPT and KHV; the same segment was also observed at 52.4% in PEL (Supplementary Table S16; Fig. 6a, b; Fig. S6.1, S6.2). Absolute nucleotide distances to the Neanderthal reference genome exhibited a bimodal distribution, with introgressed haplotypes forming a low-distance component separated from African haplotypes (Fig. 6a). Within this region, at least 26 informative alleles reached high frequency in East Asian populations while being absent or rare in African haplotypes, and approximately 73% of these alleles matched Neanderthal references (Fig. 6b). A maximum-likelihood haplotype tree further showed that a subset of present-day haplotypes clustered closely with Neanderthal haplotypes, supporting the Neanderthal-related origin of this AS3-specific introgressed segment (Fig. S6.2).

**Fig. 6.**
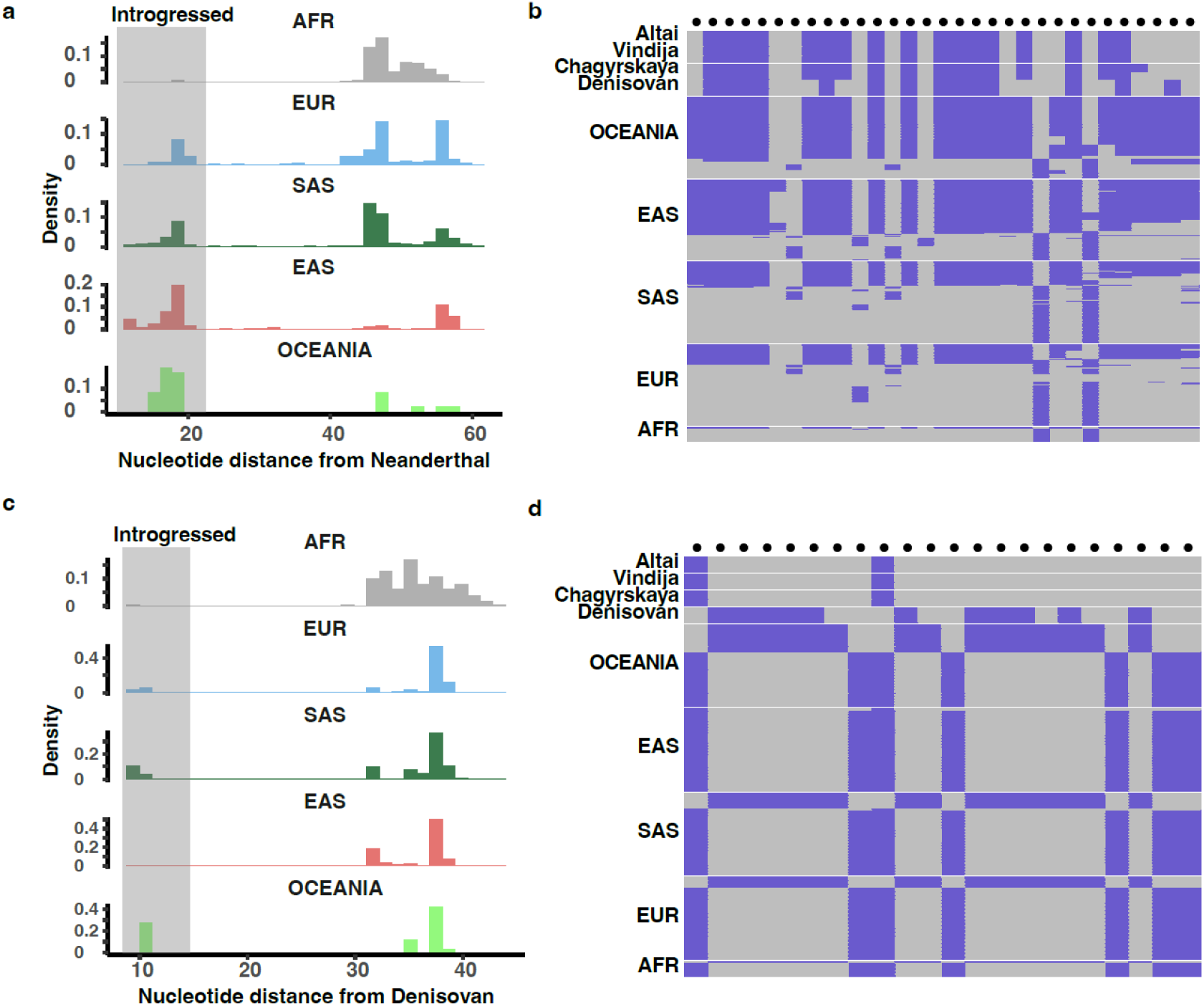
High-frequency AS3-specific introgressed segments supported by locus-level haplotype patterns. (a) Distribution of absolute nucleotide distances to the Neanderthal reference for haplotypes spanning the chr20:63,697,389–63,769,790 Neanderthal-introgressed segment. The grey shaded interval marks the distance range occupied by AS3-inferred introgressed haplotypes. Population groups are shown at right. (b) Haplotype-state matrix for informative SNPs at the chr20 Neanderthal-introgressed segment. Rows represent archaic references and modern haplotypes grouped by population region; columns represent informative SNPs absent or rare in African haplotypes. Purple indicates the derived allele and grey indicates the ancestral allele; when the ancestral state is unavailable, purple indicates the alternative allele and grey indicates the reference allele. Black dots mark informative sites. (c) Distribution of absolute nucleotide distances to the Denisovan reference for haplotypes across the chr11 window encompassing the QC-refined Denisovan-introgressed segment at chr11:30,668,800–30,706,896. (d) Haplotype-state matrix for informative SNPs at the chr11 Denisovan-introgressed segment, shown as in (b). AFR, African; EUR, European; SAS, South Asian; EAS, East Asian; OCEANIA, Oceanian/Papuan haplotypes.

We also identified a high-frequency AS3-specific Denisovan-introgressed segment in Papuans at chr11:30,668,800–30,706,896, spanning approximately 38.1 kb and reaching a frequency of 50.0% (Supplementary Table S16; Fig. 6c, d; Fig. S6.3, S6.4). Introgressed haplotypes at this locus showed substantially lower nucleotide distance to the Denisovan reference than African haplotypes, with approximately nine differences in the introgressed haplotypes compared with approximately 36 differences in African haplotypes (Fig. 6c). Within this region, at least 23 informative alleles reached high frequency in Papuan haplotypes but were absent or rare in African haplotypes, and approximately 78% matched the Denisovan reference (Fig. 6d). Phylogenetic reconstruction further showed that a subset of Papuan haplotypes clustered near the Denisovan lineage, supporting Denisovan introgression for this AS3-specific segment (Fig. S6.4).

Gene annotations further provided biological context for these representative AS3-specific introgressed segments. The chr20 Neanderthal-introgressed segment falls in a gene-rich 20q13.33 interval annotated with *RTEL1-TNFRSF6B*, *TNFRSF6B*, *ARFRP1*, *ZGPAT*, *LIME1*, *SLC2A4RG* and *ZBTB46* (Supplementary Table S16). Several genes in this interval have reported roles in immune signaling, transcriptional regulation or metabolic regulation: *TNFRSF6B*/*DcR3* was originally described as a soluble decoy receptor for Fas ligand^49^, *LIME1* has been implicated in T-cell activation^50^, *SLC2A4RG*/*GEF* has been linked to transcriptional regulation of the *GLUT4* promoter^51^, and *ZBTB46* has been used as a marker of classical dendritic-cell lineage commitment^52^. In contrast, the chr11 Denisovan-introgressed segment was not assigned an overlapping protein-coding gene in Supplementary Table S16, but the local gene track includes *ENSG00000309197*, a poorly characterized non-protein-coding transcript annotation^53^ that overlaps part of the refined interval (Fig. S6.3). Together, these examples show that high-frequency AS3-specific introgressed segments are supported by independent haplotype-level and phylogenetic evidence, and illustrate how AS3 can prioritize functionally annotated introgressed intervals for downstream investigation, highlighting its potential to uncover biologically informative introgression signals beyond those captured by existing callsets.

## Discussion

ArchaicSeeker 3.0 (AS3) is a haplotype-resolved framework for archaic introgression inference designed to remain accurate across demographic regimes while being practical to deploy at biobank scale. Across simulations spanning multiple demographic models and empirical concordance analyses, AS3 delivers strong tract detection and archaic-source classification while improving boundary localization relative to existing approaches (Fig. 3; Fig. 4). More broadly, AS3 illustrates a complementary inference strategy for a class of population-genetic problems in which the primary goal is local ancestry or segment-level state assignment under complex and only partially specified histories. For such problems, performance often depends less on precise estimation of a small number of global demographic parameters than on accurate discrimination of local sequence patterns and source-consistent haplotypic structure. A key advantage over cohort-calibrated LAI pipelines is AS3’s haplotype-wise inference paradigm: because inference is performed independently per haplotype using a fixed learned decision rule, the model does not require cohort-level parameter estimation (such as allele frequency recalibration) at test time. Consequently, AS3 remains stable even when the target cohort is small or structurally unbalanced (Fig. 3a), enabling robust analyses in under-sampled populations.

Mechanistically, AS3 achieves this combination of robustness and deployability through three complementary design choices. First, the base model compresses reference information into simple per-site match statistics and combines these with a distance channel, allowing the network to operate with a constant-width input representation independent of target cohort size. Second, we depart from standard Transformer architectures by adopting a selective state-space backbone (based on Mamba-2). This allows AS3 to model multi-locus haplotypic patterns across 4,096-SNP windows with linear computational complexity, avoiding the quadratic memory bottleneck of attention mechanisms while capturing richer local sequence structure than more strongly localized approaches. Third, our ablation analyses indicate that robustness is primarily driven by diversity in the training demographic distribution rather than sample count alone. Importantly, the training simulations do not need to exhaustively enumerate every evaluation-time demographic model. Instead, robustness appears to emerge when the model is exposed to a sufficiently broad range of tract-generating histories from which more transferable sequence-level decision rules can be learned. In this sense, AS3 does not simply fit a narrow family of demographic templates, but captures part of the relevant historical complexity through a learned inference rule.

The operating regime experiments clarify when and why performance degrades under practical constraints. Reducing marker density leads to a decline in archaic-seeking sensitivity, consistent with the loss of informative haplotypic structure needed to detect introgressed tracts; however, our benchmarks indicate the model retains practical utility even at densities characteristic of genotyping arrays (Fig. 3d). Variant ascertainment is a stronger bottleneck: filtering targets toward common variation substantially reduces tract detection, indicating that low-frequency and private-allele information contributes disproportionately to identifying introgressed segments (Fig. 3e). By contrast, recombination-map inputs are not a strict requirement: using a constant-rate approximation performs comparably to fine-scale maps (Fig. S3.9), suggesting that the positional signal encoded by our feature construction and smoothing pipeline is sufficient for accurate boundary placement in typical use cases. Importantly, boundary fidelity is not merely a cosmetic property of the final calls. Because tract length distributions carry information about the timing, multiplicity, and structure of introgression events, improved cross-window coherence and more accurate boundaries may also improve downstream historical interpretation, including analyses of whether admixture occurred in one or multiple pulses and on what timescale.

Beyond accuracy, AS3 is engineered to scale near-linearly with data volume under chunked, batched inference. Inference requires only phased target genotypes and a phased reference panel, enabling embarrassingly parallel execution across haplotypes and genomic regions. Profiling results show high throughput with a compact memory footprint (Table 1), supporting the feasibility of UK Biobank–scale analyses under standard cluster scheduling (Table 2). Together with the orthogonal concordance observed on phased 1000 Genomes samples (Fig. 4) and locus-level support for high-frequency AS3-specific introgressed segments (Fig. 6), these results support AS3 as a practical foundation for large-scale studies of human genetic diversity. Although archaic introgression is the primary use case examined here, the same general framework may also be relevant to related problems that can be formulated as local sequence- or tract-level labeling tasks under realistic simulations, including local ancestry inference in admixed populations and, in principle, tract detection involving unsampled donor lineages.

Several limitations motivate future work. First, AS3 currently assumes phased inputs and data quality broadly comparable to modern sequencing or genotyping datasets. Although training perturbations such as offset jitter and allelic flip noise improve tolerance to modest errors, substantial phasing switch errors, extensive missingness, and uncertain genotype calls—as are common in ancient DNA and very low-coverage data—would be expected to reduce accuracy. In addition, sensitivity depends on marker density and variant ascertainment, so heavily filtered datasets require cautious interpretation. More fundamentally, the genome is not a homogeneous substrate: mutation rates vary across loci, and local recombination patterns, sequence architecture, and functional constraint can all influence the visibility of introgressed haplotypes and the stability of tract boundaries. These sources of heterogeneity are only partially represented by the current training simulations and compact per-site encoding, and incorporating richer error models and more heterogeneous training regimes should further improve robustness.

A further limitation concerns the complexity of archaic population history itself. AS3 identifies archaic segments relative to the reference lineages currently available, but early human gene flow was likely more complex than a simple Neanderthal-versus-Denisovan labeling scheme, potentially involving unsampled ghost lineages as well as admixture among archaic groups. Under such scenarios, some difficult-to-interpret segments may reflect genuine archaic ancestry that is imperfectly captured by present-day source labels rather than spurious signal. Better resolution of these segments will require both methodological extensions—such as uncertainty-aware or open-set source assignment—and additional high-quality archaic genomes. Importantly, this limitation does not diminish the value of archaic-segment discovery itself; rather, it underscores that robust detection and source interpretation are related but distinct goals, and that more complete archaic references will be essential for reconstructing the full complexity of early human evolutionary history.

## Code availability statements

The code for ArchaicSeeker3.0 is available at the GitHub repository: https://github.com/Shuhua-Group/ArchaicSeeker3.0. The repository includes the core source code, installation files, example workflows, and scripts for preprocessing and running the main analysis pipeline. It also provides code for benchmarking and simulation-based evaluation, together with demographic-model configuration files, enabling users to reproduce the main computational experiments reported in this study. In addition, pretrained model weights are provided in the exp directory, allowing users to run inference directly without retraining the model.

## Acknowledgements

This study was supported by the National Key Research and Development Program of China (No. 2023YFC2605400), the Shenzhen Medical Research Fund (C2503001), the National Natural Science Foundation of China (NSFC) grants (32288101, 12271026), the Shanghai Science and Technology Commission Program (25JS2810100, 23JS1410100, QNKJ2024023), the Fundamental Research Funds for the Central Universities (2025JBZY012), Fundamental and Interdisciplinary Disciplines Breakthrough Plan of the Ministry of Education of China (JYB2025XDXM508), Fund of Fudan University and Cao’ejiang Basic Research (24FCB08), and the Office of Global Partnerships (Key Projects Development Fund). The CFFF Computing Platform of Fudan University supported the computational work in this study.

## Author Contributions

S.X. conceived and designed the study. S.X. supervised the project. B.W., C.L. and H.L. developed the algorithm and wrote the code for ArchaicSeeker 3.0. H.L. contributed to the evaluation pipeline and data processing for AS3. S.S. carried out part of the real-data evaluation and developed the front-end of the AS3 web server. X.M. and W.Z. carried out part of the real-data analyses using AS3. K.Y. and X.N. revised the manuscript and provided helpful suggestions. B.W. wrote the manuscript and prepared the supplementary materials. S.X. substantially revised the manuscript. All authors discussed the results and implications and commented on the manuscript.

## Declaration of interests

The authors declare no competing interests.

## Notes

https://sharehost.hms.harvard.edu/genetics/reich_lab/sgdp/phased_data2021/

https://web.archive.org/web/20170809212155/ https://evolbio.ut.ee/CGgenomes_VCF

https://reich.hms.harvard.edu/index.php/datasets

https://storage.googleapis.com/gcp-public-data--gnomad/release/3.1.2/vcf/genomes

https://ftp.1000genomes.ebi.ac.uk/vol1/ftp/data_collections/1000G_2504_high_coverage/working/20201028_3202_raw_GT_with_annot/

https://data.mendeley.com/datasets/y7hyt83vxr/1

https://doi.org/10.5281/zenodo.14136628

https://doi.org/10.5281/zenodo.14552025

https://github.com/Shuhua-Group/ArchaicSeeker2.0/tree/master/IntrogressedSeg/KGP

https://zenodo.org/records/13368126

### Competing Interest Statement

The authors have declared no competing interest.

