## Supplementary Information for "*ArchaicSeeker* 3.0: A deep-learning framework for scalable, haplotype-resolved inference of archaic introgression"

We generated simulated datasets using msprime[1-3] and adopted demographic models and parameterizations curated in stdpopsim[4,5].

### Benchmark

We benchmarked ArchaicSeeker 3.0 against other widely used, published methods for detecting archaic introgression under a unified evaluation protocol on an extensive suite of simulated datasets, demonstrating the superior performance of ArchaicSeeker 3.0.

#### Data simulation and ground truth

We simulated data under nine demographic models (Table S9), including seven models in human populations and two models in the chimpanzee–bonobo system. For the human settings, we organized the seven models by the number of introgressing sources, the temporal architecture of gene flow and the phylogenetic origin of the introgressing lineages, thereby covering a broad set of representative scenarios. Specifically, four models assume single-source introgression in which present-day humans experienced one episode of gene flow from a single archaic lineage (for example, Neanderthal or Denisovan), whereas three models represent dual-source introgression by introducing gene flow from distinct archaic lineages (for example, multiple Neanderthal lineages or joint Neanderthal–Denisovan introgression) to emulate more complex histories. These models additionally span different temporal modes, including continuous gene flow (CGF), in which exchange persists over an extended interval, and admixture-pulse models, in which introgression is concentrated at one or a few discrete time points; in selected configurations, we further introduced deep African lineages to assess the impact of complex African structure and deeper lineage divergence. By jointly considering these diverse human demographic models, we systematically evaluated the robustness and applicability of the proposed method under progressively more challenging conditions, with increasing introgression-ancestry complexity, temporal architectures and lineage depth.

For each demographic model, we simulated 50-Mb genomic sequences with 100 replicates and report performance averaged across replicates. We subsampled the reference panel to 10 or 50 diploid individuals and the target panel to 1 or 10 diploid individuals, emphasizing detection performance at the level of single individuals, which is often the most biologically informative resolution for introgressed-tract discovery. Simulated data were stored in both tree-sequence (ts) and VCF formats. For VCF-based analyses, we filtered SNPs to retain only biallelic sites, and all methods were evaluated on the same filtered VCFs.

Because the ancestral recombination graph (ARG) encoded by the tree sequence records the complete genealogical history, we extracted ancestry tracts from the ts as ground truth. For single-source introgression models, ground-truth tracts were defined as segments in the target population (tgt) inherited from the introgressing source population (src). For dual-source introgression models, we extracted tgt segments inherited from src1 and src2 separately (labeled src1 and src2) and additionally merged them into a combined archaic label (src), allowing us to evaluate archaic-seeking performance independently of any downstream source-matching procedure.

**Software execution and result processing**

We systematically compared Sprime, IBDmix, hmmix, DAIseg, ArchaicSeeker 2.0 and ArchaicSeeker 3.0. Although all of these methods were developed to detect archaic introgression, they differ in both inference targets and input requirements (Table S11). ArchaicSeeker 3.0 does not require additional prior information; however, to ensure that competing methods were evaluated under their best-performing settings, we supplied all required model- and data-generating parameters (for example, mutation rate, recombination rate and phylogenetic configuration) according to the simulation settings.

For Sprime, the reported score reflects confidence in introgression. We retained only calls with scores above 150,000, as recommended in the original publication. Under dual-source introgression models, we further retained only records with similarity > 0 to at least one source and assigned the source label according to the higher similarity.

IBDmix does not require an African reference panel but expects at least two target individuals as input. Therefore, when $n_{tgt} = 1$, we duplicated the target individual to meet the input requirement. IBDmix reports an sLOD score as a measure of confidence, and we retained only calls with sLOD > 4. Although IBDmix was originally developed for Neanderthal introgression and its parameters are not tailored to other archaic lineages, we nevertheless applied it in dual-source settings by running the method separately for each archaic source, retaining sLOD > 4 calls and assigning the corresponding source label, and finally merging the two call sets to obtain a combined archaic label (src).

For hmmix, we retained records whose inferred state was Archaic and for which the target shared at least one SNP with an archaic reference. In dual-source settings, we first treated all retained records as archaic (src) and then determined the source label by comparing whether the target shared more SNPs with src1 or with src2.

For DAIseg, we constructed the demographic history model file required for inference by following the model settings used in ArchaicSeeker 2.0 and leveraging stdpopsim. We retained only segments whose decoded state corresponded to an archaic ancestry. Under dual-source introgression settings, DAIseg can distinguish the two archaic sources (e.g., Denisovan and Neanderthal) in a single run; thus, segments decoded as src1 or src2 were labeled as the corresponding source.

For ArchaicSeeker 2.0, we used the BestMatchedPop field to assign ancestry. Specifically, BestMatchedPop reports the branch and temporal depth on the phylogenetic tree that best matches each segment. Segments mapping exclusively to src1 or to src2 were labeled as src1 or src2, respectively; segments mapping to src1, src2 or src1_src2 (representing the ancestral branch shared by src1 and src2) were additionally grouped under the combined archaic label (src).

For ArchaicSeeker 3.0, we retained records with Score ≥ 0.6. ArchaicSeeker 3.0 outputs an inferred ancestry label for each segment; in addition to src1 and src2, it may also return segments labeled as mosaic. For analyses focused on detecting archaic tracts irrespective of source, we merged src1, src2 and mosaic into the combined archaic label (src).

**Evaluation metrics**

Notably, only hmmix, DAIseg, ArchaicSeeker 2.0 and ArchaicSeeker 3.0 provide haplotype-resolved outputs, and only the ArchaicSeeker methods explicitly support discrimination among archaic sources. Accordingly, we unified all methods at the sample level for benchmarking, with an emphasis on their ability to recover archaic segments; because Sprime operates at the population level, it is denoted separately in figures.

**Ability to identify archaic segments**

For each method under each demographic model, we evaluated the ability to identify archaic segments by computing precision, recall and F1 from the overlap between simulated (ground-truth) tracts and inferred tracts.

$$Precision = \frac{TP}{TP + FP} = \frac{L_{overlapping}}{L_{infered}}$$

$$Recall = \frac{TP}{TP + FN} = \frac{L_{overlapping}}{L_{simulated}}$$

Here, $TP$, $TP$ and $FN$ denote true positives, false positives and false negatives, respectively. $L_{overlapping}$ is the total base-pair overlap between inferred and ground-truth archaic segments, $L_{infered}$ is the total inferred archaic length, and $L_{simulated}$ is the total ground-truth archaic length.

$$F1 = \frac{2 \times Precision \times Recall}{Precision + Recall}$$

We summarized performance using precision–recall scatter plots across demographic models. All methods benefited from increased numbers of target samples and reference individuals; therefore, for bar-plot comparisons we focused on the setting $n_{tgt}$=10, $n_{ref}$=50. Because Sprime yields population-level calls, its metrics can only be computed at the population level, which may confer a slight advantage; we thus indicated Sprime results with a dashed outline for clarity.

In addition, we reported the ratio

$$Ratio = \frac{L_{infered}}{L_{simulated}}$$

to quantify whether inferred archaic content tends to overestimate or underestimate the true amount of introgression.

Beyond length-based metrics, we also evaluated performance at the segment level. A simulated (ground-truth) segment was considered successfully recovered if the overlap between that segment and any predicted segment covered at least 80% of the simulated segment length. Because hmmix typically outputs shorter fragments and Sprime reports population-level introgressed regions rather than per-individual segments, these two methods were excluded from this segment-level analysis.

Several demographic models included introgression from Neanderthals (six models) and/or Denisovans (three models). For these two archaic lineages, we aggregated results and reported mean performance separately, using the same metrics as above. This analysis reflects overall capability in both detecting archaic segments and distinguishing their sources when relevant.

In two-source introgression models, DAIseg requires two distinct outgroup references to separate segments from different archaic sources; however, for human populations, identifying appropriate outgroups that satisfy this assumption is often impractical. We therefore excluded DAIseg from source-specific performance evaluation of single archaic ancestries under the two-source setting.

We further compared ArchaicSeeker 3.0, ArchaicSeeker 2.0, hmmix, IBDmix and Sprime with respect to their ability to distinguish archaic sources under two-source introgression. Notably, only ArchaicSeeker methods incorporate source discrimination in their core model; the remaining methods rely on post hoc matching to archaic reference genomes to assign ancestry.

Specifically, we first retained segments whose overlap with the corresponding ground-truth introgressed segments was at least 0.8, treating them as successfully detected archaic segments. Among these, a segment was counted as correctly classified if the inferred source label matched the true simulated source. We then defined source-classification accuracy (ACC) as the proportion of correctly classified segments among all successfully detected archaic segments.

**Reconstructing segment-length distributions and localizing segment boundaries**
Recombination is a key mechanism shaping the structure of introgressed ancestry tracts: repeated recombination events fragment introgressed material, and the resulting tract-length distribution is informative about the timing, intensity and dynamics of introgression. We therefore evaluated ArchaicSeeker 3.0, ArchaicSeeker 2.0, IBDmix and DAIseg in their ability to localize tract boundaries (that is, recombination breakpoints) and to reconstruct tract-length distributions. Because hmmix and Sprime do not directly output continuous tracts with explicit boundaries by design, they were not included in this analysis.

Unless otherwise noted, we used $n_{tgt} = 10$、$n_{ref} = 50$. For each human demographic model, we computed tract lengths from both simulations and inferred outputs, standardized lengths to kilobases, and compared the inferred and ground-truth length distributions for tracts of at least 10 kb.

To further quantify each method's ability to localize tract boundaries, we focused on segments that were successfully detected, defined as inferred tracts whose overlap with the simulated ground-truth introgressed tract was at least 0.8. For each such tract, we recorded the simulated (ground-truth) boundaries $[l_{\mathrm{sim}}, r_{\mathrm{sim}}]$ and the inferred boundaries $[l_{\mathrm{infer}}, r_{\mathrm{infer}}]$. We then defined the relative distance of recombination breakpoints as follows.

$$Relative\ Distance = \frac{|l_{\text{sim}} - l_{\text{infer}}| + |r_{\text{sim}} - r_{\text{infer}}|}{r_{\text{sim}} - l_{\text{sim}}}.$$

This metric summarizes the normalized boundary deviation by measuring the combined displacement of the left and right boundaries relative to the true tract length, thereby quantifying the precision with which each method localizes ancestral recombination breakpoints.

### Characterization of ArchaicSeeker 3.0

To systematically characterize the behaviour of ArchaicSeeker 3.0 across different input configurations and to quantify how performance changes with input scale, we performed controlled experiments in which we varied the size of each major input component. The three primary inputs to ArchaicSeeker 3.0 are the target samples (tgt), a modern reference panel (modern reference) and an archaic reference panel (archaic reference).

We assessed performance at two levels: detection of introgressed tracts and discrimination of archaic sources. Specifically, we report length-based precision, recall, F1 and ratio for archaic, Neanderthal (Nean) and Denisovan (Den) tracts, and additionally compute count-based precision at the segment level. To quantify source discrimination, we report the accuracy (ACC) for distinguishing Nean from Den, together with misclassification rates for Nean-to-Den and Den-to-Nean errors.

All analyses were conducted under the Papuan_NeanDen demographic model. This model was not among the primary training models for ArchaicSeeker 3.0, and the Papuan population typically harbours a relatively high proportion of Denisovan ancestry; it therefore provides a representative two-source human introgression setting for systematically evaluating source discrimination. Under this model, we simulated an original dataset comprising 1,000 target individuals, 100 modern reference individuals, 10 Neanderthal individuals and 10 Denisovan individuals, with a genome length of 10 Mb. Each experimental configuration was repeated 100 times to reduce stochastic variation.

To evaluate the effect of target sample size, we fixed a subset of 50 modern reference individuals, 3 Neanderthal individuals and 1 Denisovan individual from the original dataset, and constructed test sets with $n_{tgt}$ =1, 10, 25, 50, 100, 250, 500 and 1,000.

To evaluate the effect of modern reference-panel size, we fixed 10 target individuals together with 3 Neanderthal and 1 Denisovan individuals, and varied the modern reference-panel size by setting $n_{ref}$ =5, 10, 25, 50 and 100.

To evaluate the influence of archaic reference-panel composition, we considered both balanced and imbalanced settings. In the balanced setting, we fixed 50 modern reference individuals and 10 target individuals and set the total number of archaic reference individuals to 2, 6, 10 and 20, corresponding to equal numbers of Neanderthal and Denisovan individuals (1, 3, 5 and 10, respectively). Given the limited availability of high-coverage archaic genomes in practice (commonly 3 Neanderthal individuals and 1 Denisovan individual), we further evaluated imbalanced configurations. When varying the number of Neanderthal individuals, we fixed 50 modern reference individuals, 10 target individuals and 1 Denisovan individual and set the number of

Neanderthal individuals to 1, 2, 3, 5 and 10; conversely, when varying the number of Denisovan individuals, we fixed 50 modern reference individuals, 10 target individuals and 3 Neanderthal individuals and set the number of Denisovan individuals to 1, 2, 3, 5 and 10.

**Robustness of ArchaicSeeker 3.0**

To assess the robustness of ArchaicSeeker 3.0 (AS3) under practical conditions, we systematically examined how performance changes under data degradation and common pre-processing choices that may arise in real applications. Specifically, we considered three perturbation factors: variation in SNP density, filtering of low-frequency variants in the modern reference panel, and filtering of low-frequency variants in the target data. In all robustness analyses, we used the same underlying simulated datasets and evaluation metrics as described above, and introduced perturbations only by constructing modified test datasets under each condition.

Effective SNP density in empirical datasets can vary substantially due to sequencing depth and coverage, variant calling and quality-control procedures, array or capture design, and downstream filtering and missing-data handling, which in turn may affect methods that rely on marker density. To emulate different SNP densities, we randomly masked a fraction of variant sites in the original simulated VCF. We applied masking ratios of 0, 10, 20, …, 90% and re-computed SNP density for each setting. We then constructed test datasets with fixed parameters ($n_{ref}$=50, $n_{tgt}$=10, $n_{Nean}$=3, $n_{Den}$=1) to quantify AS3 performance as SNP density decreases.

Because AS3 relies substantially on low-frequency variants in the modern reference panel, and such variants can be difficult to obtain at sufficient quantity and quality in practice, we next evaluated the impact of progressively filtering low-frequency variants in the reference panel. In this analysis, we fixed $n_{ref}$=100, $n_{tgt}$=10, $n_{Nean}$=3, $n_{Den}$=1 and extracted the corresponding subset from the original dataset. We then applied a series of minor allele frequency (MAF) filters to the reference-panel VCF, retaining only SNPs that satisfied MAF > 0, 0.01, 0.02, 0.1 or 0.2 in the reference panel, thereby generating a set of test datasets for benchmarking.

In many applied settings, low-frequency variants in the target cohort are filtered to improve analytical stability under certain conditions. We therefore further examined the effect of filtering low-frequency variants in the target data on AS3 inference. We fixed $n_{ref}$=50, $n_{Nean}$=3 and $n_{Den}$=1 , first computed allele frequencies within the target data, and then constructed filtered target VCFs under multiple thresholds. These included filters based on minor allele count (MAC > 0, 1, 2 or 5) and minor allele frequency (MAF > 0.005, 0.01, 0.025, 0.05, 0.1, 0.25 or 0.5). From each filtered dataset, we sampled 10 target individuals to form test sets and systematically evaluated how target-side rare-variant filtering influences AS3 performance.

**Computational cost of ArchaicSeeker 3.0**

To characterize the computational cost of ArchaicSeeker 3.0 (AS3) under varying data scales and configurations, we profiled wall-clock runtime, peak resident set size (peak RSS) and peak GPU memory usage across a range of settings. Guided by AS3's algorithmic structure and typical use cases, we focused on four key factors expected to drive resource consumption and adopted a one-factor-at-a-time design by varying each factor while holding the others fixed. All experiments were conducted on a Linux server running CentOS 7, equipped with dual Intel Xeon Silver 4310 CPUs (2.10 GHz), 64 GB of RAM, and four NVIDIA GeForce RTX 4090 GPUs (24 GB VRAM each).

When assessing the effect of target cohort size, we fixed the modern reference-panel size at $n_{ref}$=50, used a SNP density of approximately 19,505 SNP/Mb and a genome length of 10 Mb, and constructed datasets with $n_{tgt}$=1, 10, 25, 50, 100, 250, 500 and 1,000. To evaluate the effect of modern reference-panel size, we fixed $n_{tgt}$=10 with the same SNP density (≈19,505 SNP/Mb) and genome length (10 Mb) and varied $n_{ref}$=5, 10, 25, 50 and 100. We further examined SNP-density effects by fixing $n_{tgt}$=10, $n_{ref}$=50 and a 10-Mb genome and adjusting the number of variant sites to span densities from approximately 1,950 to 19,505 SNP/Mb. Finally, to assess scaling with genome length, we fixed $n_{tgt}$=1, $n_{ref}$=50 and SNP density at approximately 13,344 SNP/Mb and varied the simulated genome length across 10, 20, 50, 75, 100, 150, 200 and 250 Mb; the 250-Mb setting exceeds the length of the longest human chromosome and was included to probe AS3's resource usage at very large genomic scales.

**Parameter settings for ArchaicSeeker 3.0**

To systematically assess how key parameters and configuration choices affect ArchaicSeeker 3.0 (AS3), we performed one-factor sensitivity analyses under a unified demographic model and simulation framework. All experiments used the Papuan_NeanDen model, with 1 target individual, 50 modern reference individuals, 3 Neanderthal individuals and 2 Denisovan individuals. Simulations were carried out on human chromosome 19 (58,585,793 bp), assuming genome-wide constant mutation and recombination rates (mutation_rate = $1.4 \times 10^{-8}$; r = $1.83848 \times 10^{-8}$), following the human reference settings in stdpopsim. Each parameter configuration was replicated 100 times to reduce stochastic variability.

We first examined the effect of filtering short, fragmented calls before the merge step. Predicted segments were grouped according to whether they overlapped the simulated ground-truth tracts, and we compared the resulting groups in terms of their length and score distributions. We then evaluated tract-detection performance (F1) under a grid of filtering thresholds, with Min Length set to 0.5, 1, 2, 3, 4 or 5 kb and Min Score set to 0.4, 0.5, 0.55, 0.65, 0.7 or 0.8. Although these filters can be tuned to specific study goals, under the settings considered here we recommend removing segments no longer than 5 kb prior to merging, while not imposing an additional score threshold at this stage.

Building on the pre-merge filtering strategy above, we next evaluated the Merge Distance parameter, using F1 for archaic-tract detection as the performance metric. We systematically tested Merge Distance values of 0, 1,000, 2,500, 5,000, 7,500, 10,000, 12,500, 15,000, 17,500, 20,000, 25,000, 30,000, 40,000 and 50,000 bp. While the optimal value may depend on dataset characteristics, we found Merge Distance = 10,000 bp to be a robust default under the conditions considered here.

After merging, we further assessed a joint filtering strategy based on segment score and segment length (Score–Length filtering) (Fig. S3.7; Fig. S3.8). In this analysis, we reported length-weighted precision, recall, F1 and ratio for archaic segments, along with segment-level precision and the accuracy (ACC) of discriminating Neanderthal and Denisovan sources. Although thresholds can be adapted to specific applications, we recommend retaining segments with Score ≥ 0.85 and length ≥ 15 kb in the setting studied here.

AS3 additionally supports supplying a recombination map as an auxiliary input. To evaluate whether providing recombination-rate variation yields measurable gains, we designed paired control experiments that followed the simulation setting above but differed in the recombination process: one dataset assumed a constant recombination rate, whereas the other used an empirical recombination map to model position-dependent recombination intensity along chromosome 19. Recombination parameters and maps were taken from the human reference settings in stdpopsim. We then ran AS3 with and without the recombination map provided as input for each dataset, enabling a direct assessment of the utility of recombination-map information when recombination-rate variation is known or unknown.

Unless otherwise specified, AS3 uses the default model, inference, decoding, and annotation parameters summarized in Supplementary Table S15. The pre-merge, merge, and post-merge defaults listed below were guided by the one-factor sensitivity analyses shown in Fig. S3.7 and Fig. S3.8.

### Cross-validation with published results

#### Downloading and processing KGP2504

Because most previously reported archaic-introgression callsets are derived from the 2,504 unrelated individuals released in 1000 Genomes Project Phase 3[6,7], we also ran ArchaicSeeker 3.0 on this dataset. We started from the original unphased data and performed statistical phasing using SHAPEIT5[8]. We filtered variants by retaining only sites that passed Variant Quality Score Recalibration (VQSR), restricting to biallelic SNPs, and removing sites with missing genotypes. After filtering, we obtained 99,084,738 SNPs (Table S10). In SHAPEIT5, variants with minor allele frequency (MAF) below 0.01 were treated as rare, and a two-step strategy was used to phase common and rare variants separately to improve phasing accuracy.

#### Downloading and harmonizing published introgression callsets on KGP

To minimize confounding effects introduced by heterogeneous filtering pipelines or parameter choices, we downloaded publicly released results from SPrime, hmmix, IBDmix and ArchaicSeeker 2.0 and performed lightweight harmonization[9-13]. For these callsets and for ArchaicSeeker 3.0, we restricted analyses to non-African populations in KGP for cross-validation.

For SPrime, the score represents site-level confidence in introgression, and all reported sites satisfy the recommended minimum threshold of 150,000. Following the SPrime protocol[14], we converted the raw output into a segment-level callset and then assigned putative sources by comparing segment similarity to Neanderthal and Denisovan references. Finally, we converted coordinates from hg19 to hg38 using CrossMap[15].

For hmmix, we retained introgressed segments with inferred introgression probability greater than 0.8.

We did not apply additional post-processing to the released IBDmix callset. IBDmix reports a LOD score that reflects the likelihood that a region shares identity-by-descent with a Neanderthal reference. In the released results used here, all records have LOD > 4 and segment length ≥ 50 kb. Because IBDmix was developed for detecting Neanderthal introgression and its default parameterization and filtering are not tailored to Denisovan introgression, we did not include IBDmix in Denisovan-focused comparisons[16].

For ArchaicSeeker 2.0, we converted coordinates from hg19 to hg38 using CrossMap and assigned Neanderthal versus Denisovan ancestry based on the BestMatchedPop field in the released output.

For ArchaicSeeker 3.0, post-processing used a two-stage decoding regime: pre-merge minimum length = 5 kb; merge distance $\delta$ = 10 kb; post-merge minimum score = 0.85; and post-merge minimum length = 15 kb. Only tracts passing all post-merge criteria were retained for downstream population-level summaries and cross-method comparisons.

**How we evaluated concordance**

To ensure comparability across methods, we first harmonized ancestry labels into three categories—Neanderthal, Denisovan, and Archaic—with non–single-source or ambiguous labels such as mosaic/both/none collapsed into Archaic.

For haplotype-resolved outputs from AS3, AS2, and HMMix, we performed hap-level post-processing within each sample and chromosome by splitting tracts into minimal non-overlapping intervals defined by segment boundaries and jointly considering the two haplotypes; intervals with conflicting labels were uniformly recoded as Archaic.

At the sample level, we constructed union segments using the combined breakpoint set from AS3, AS2, HMMix, and IBDmix, and annotated each union segment with whether each method had any overlapping call.

At the population level, to avoid incomparability due to unequal sample coverage across methods, we computed detection frequencies only within the intersection of samples shared by AS3, AS2, IBDmix, and HMMix. SPrime was not used for sample-level frequency estimation in this workflow; instead, we recorded a binary 1/0 indicator of whether Sprime had any overlapping call in each interval, without applying the intersection-sample filter or frequency calculation to Sprime.

**Identification of high-frequency AS3-specific introgressed segments**

We used a strategy similar to previous work[9] to summarize introgressed genomic regions carrying putative introgressed alleles at high frequency in present-day populations. Starting from post-processed AS3 calls, we extracted introgressed segments that were detected only by AS3 and did not overlap the corresponding comparison callsets used in the empirical concordance analyses. Within each population and introgression source class, adjacent AS3-specific introgressed segments were merged. We first retained introgressed alleles with frequency ≥0.30. Within each introgressed segment, we then retained alleles whose frequency was no more than 0.20 below the segment-specific maximum introgressed-allele frequency. For each initial AS3-specific segment, SNP-level quality control and frequency pruning were applied before final boundary definition; the reportable introgressed interval may therefore be narrower than the upstream AS3 segment, and final coordinates in Supplementary Table S16 were defined by the first and last retained informative introgressed alleles. Retained introgressed regions were further required to contain at least 10 alleles comparable to the corresponding archaic reference. Neanderthal-introgressed regions were retained if their match rate to Neanderthal references was >50% and higher than their match rate to the Denisovan reference. Denisovan-introgressed regions were retained if their match rate to the Denisovan reference was >40% and higher than their match rate to Neanderthal references. Retained high-frequency AS3-specific introgressed segments are summarized in Supplementary Table S16.

**Locus-level haplotype and phylogenetic analysis**

To evaluate representative AS3-specific introgressed segments, we extracted phased modern human haplotypes from 1000 Genomes populations, including African, European, South Asian and East Asian groups, together with Papuan haplotypes from HGDP. High-coverage archaic haplotypes from Altai, Vindija and Chagyrskaya Neanderthals and from the Denisovan genome were included as archaic references. Ancestral allele states were obtained from Ensembl Release 115; when ancestral states were unavailable, reference and alternate allele states were used for haplotype-matrix visualization. Haplotype sequences were multiple-sequence aligned and used for maximum-likelihood phylogenetic reconstruction with IQ-TREE[17]. ModelFinder Plus was used to select the best-fit nucleotide substitution model. Branch support was assessed using 1,000 ultrafast bootstrap replicates and 1,000 SH-aLRT replicates, with up to 2,000 tree-search iterations and automatic thread allocation. Final trees were visualized and annotated in R using ggtree[18].

### Application to real-world data

**Overview of modern datasets**

To systematically characterize the global landscape of archaic introgression in present-day populations, we assembled a broad collection of modern human genomic datasets spanning worldwide genetic diversity. Specifically, we integrated genomes from KGP[6,7], HGDP[19], SGDP[20,21], EGDP[22], and HO [23,24], covering 8 regions, 209 populations and 3453 samples in total (Table S14).

Because many existing introgression-detection methods rely heavily on population-level parameter estimation and therefore require substantial sample sizes, they often yield unstable results for small or sparsely sampled groups. ArchaicSeeker 3.0 does not depend on population-level parameter estimation, enabling us to include a substantially broader set of global populations and to construct a more complete landscape of archaic introgression.

**Processing of the HGDPTGP dataset**

For KGP and HGDP, we used a jointly called, harmonized dataset constructed from 1kGP and HGDP samples, hereafter referred to as HGDPTGP [25]. This resource combines 4,094 high-quality whole-genome samples from 80 populations across HGDP and 1kGP and incorporates allele-frequency information from gnomAD [26], yielding more than 153 million high-quality variants (SNPs, indels and structural variants). To obtain a reliable variant set for downstream inference, we performed a systematic and stringent quality-control pipeline, with most processing carried out using bcftools.

At the variant level, we first removed SNPs located within ±5 bp of indels to reduce errors introduced by local alignment instability. We then retained only biallelic SNPs that passed VQSR and were labeled PASS, while filtering out all indels and structural variants to better satisfy downstream modeling assumptions. To further mitigate sequencing noise and potential mapping artifacts, we excluded singletons, removed SNPs in low-complexity regions (LCRs) and segmental duplications, and applied the GRCh38 accessibility mask from KGP to further increase the reliability of the variant set.

At the sample level, we partitioned the dataset into seven geographic subsets (AFR, MID, EUR, EAS, CSA, OCE and AMR) and estimated pairwise relatedness within each subset using PLINK. For any samples related within the second degree, we retained only one individual and removed the remaining related samples to ensure approximate independence.

We computed site-level missingness and performed Hardy–Weinberg equilibrium (HWE) tests within each geographic subset. For each SNP, we integrated missingness and HWE results across subsets and retained only variants with a minimum HWE p value of at least $1\times10^{-30}$ across subsets, and with site missingness exceeding 0.1 in no more than three subsets.

After quality control and relatedness filtering, we merged all subsets into a unified dataset and performed phasing using SHAPEIT5 [8,27]. SHAPEIT5 applies a staged strategy that phases common and rare variants separately; we set the rare-variant frequency threshold to 0.01 to improve phasing accuracy for low-frequency and rare variants. The resulting dataset includes 3,498 samples from 80 populations across seven major geographic regions (Table S13) and 48,503,278 SNPs (Table S10).

**Harmonized processing of additional modern datasets (SGDP, EGDP and HO)**

In addition to HGDPTGP, we incorporated SGDP, EGDP and HO to further expand global population coverage.

For SGDP, we directly used the high-quality processed release from the David Reich laboratory. This dataset was constructed by Ali Akbari and imputed using 1000 Genomes Project Phase 3 as a reference. Initial variant calling was performed with bcftools (v1.10.2) [28], followed by staged phasing and imputation using GLIMPSE (v1.0.0)[29].

EGDP and HO were originally generated at lower marker density, which is typically insufficient for local ancestry inference and archaic-introgression analyses that benefit from dense genome-wide variation. We therefore applied a unified processing strategy to these datasets to enable consistent analyses alongside higher-density datasets. Briefly, we used Picard together with the UCSC LiftOver chain (hg19 to hg38) to convert variant coordinates to GRCh38. We then performed phasing and genotype imputation, using the processed HGDPTGP dataset as a reference panel and Minimac4 for imputation, thereby obtaining a high-density, genome-wide variant set.

The HO (Human Origins) dataset includes 2,345 modern human samples from 203 populations and was originally generated using the Human Origins array. EGDP was processed from its original variant data; because haplotypes had already been phased in EGDP, we skipped the phasing step and performed imputation directly. Through this harmonized phasing–imputation pipeline, EGDP and HO were standardized to the same coordinate system and broadly comparable variant density as HGDPTGP and SGDP for downstream archaic-introgression analyses.

**Archaic genomes and construction of the reference panel**

ArchaicSeeker 3.0 requires a reference panel that jointly includes modern reference samples and archaic reference genomes. Under the assumption that sub-Saharan African populations are minimally affected by known archaic introgression, we selected 146 sub-Saharan African individuals as the modern reference panel. For archaic references, we used four high-coverage archaic genomes in hg38 coordinates[30], including three Neanderthals (Altai, Vindija and Chagyrskaya)[31-33] and one Denisovan individual[34].

To construct the reference panel, we merged these 150 samples and retained only variants observed in both the modern and archaic datasets to ensure comparability across data sources. To further reduce potential impacts from sequencing errors, mapping bias and complex genomic architecture, we applied multiple masks and retained only regions with high-quality coverage and reliable mapping in KGP as well as in the Altai, Vindija, Chagyrskaya Neanderthal genomes and the Denisovan genome.

The final reference panel comprises 146 sub-Saharan African samples from 12 populations together with four archaic genomes (Table S12), and contains 34539791 SNPs. This panel provides a stable and consistent reference basis for downstream archaic-introgression inference with ArchaicSeeker 3.0.

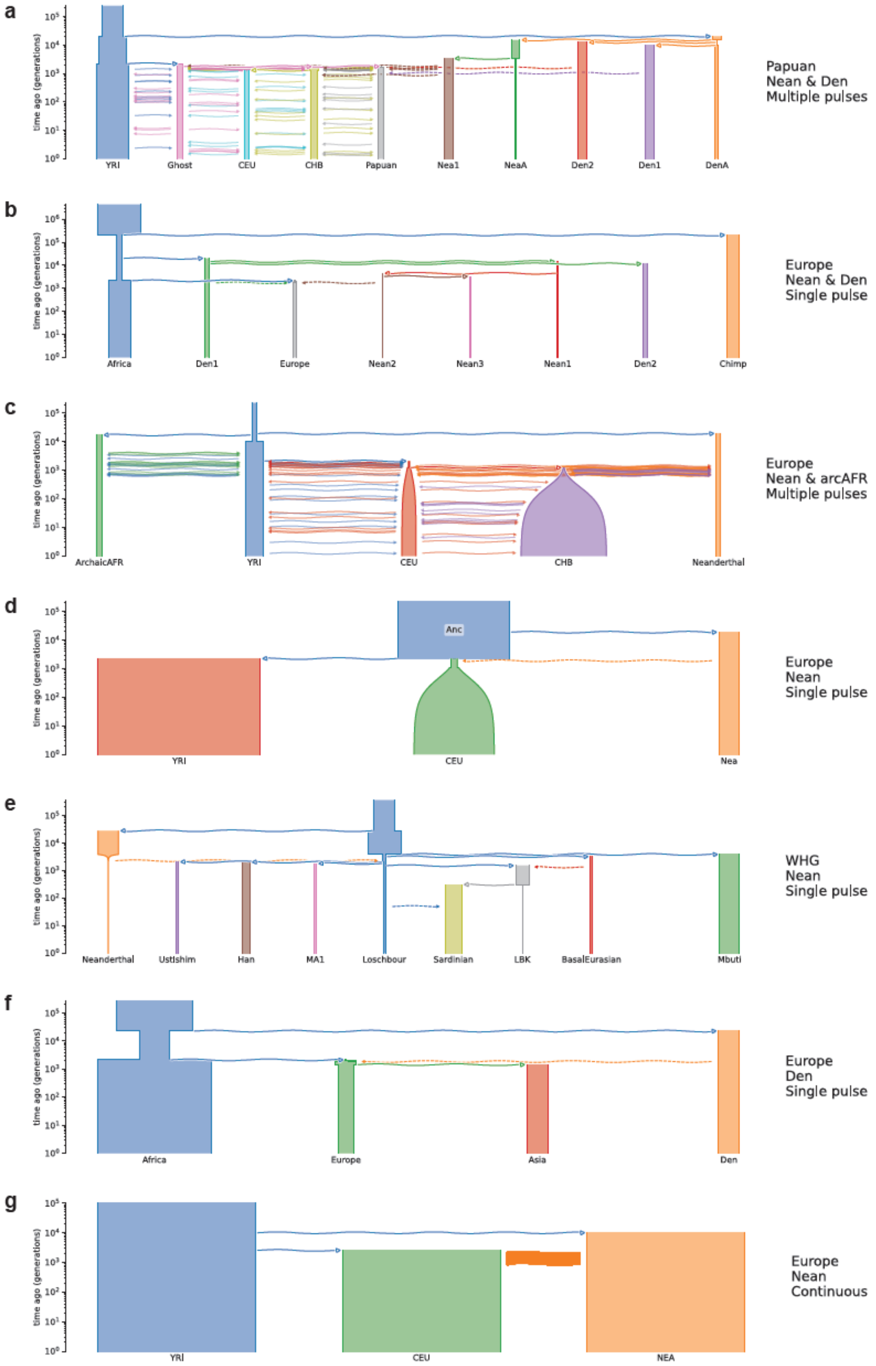


**Fig. S2.1 | Demographic models used for simulation-based validation.**
Panels (a–g) show the demographic histories used to generate reference and target haplotypes for simulation-based validation, including model training and downstream evaluation (log-scaled time axis; generations before present). Each panel is identified by its model name, admixture type, archaic source(s), reference population, and target population: (a) Papuan_NeanDen (multiple pulses; 2-source), with archaic sources Neanderthal (Nea1) and Denisovan (Den1), reference population YRI, and target population Papuans; (b) Europe_NeanDen (single pulse; 2-source), with archaic sources Neanderthal (Nean1/Nean3) and Denisovan (Den2), reference population Africa, and target population Europe; (c) Europe_NeanArcAFR (multiple pulses; 2-

source), with archaic sources Archaic African (arcAFR) and Neanderthal, reference population YRI, and target population CEU; (d) Europe_Nean_HI (single pulse; 1-source), with archaic source Neanderthal, reference population YRI, and target population CEU; (e) WHG_Nean (single pulse; 1-source), with archaic source Neanderthal, reference population Mbuti, and target population Loschbour (WHG); (f) Europe_Den (single pulse; 1-source), with archaic source Denisovan, reference population Africa, and target population Europe; and (g) Europe_Nean_CGF (continuous introgression; 1-source), with archaic source Neanderthal, reference population YRI, and target population CEU.

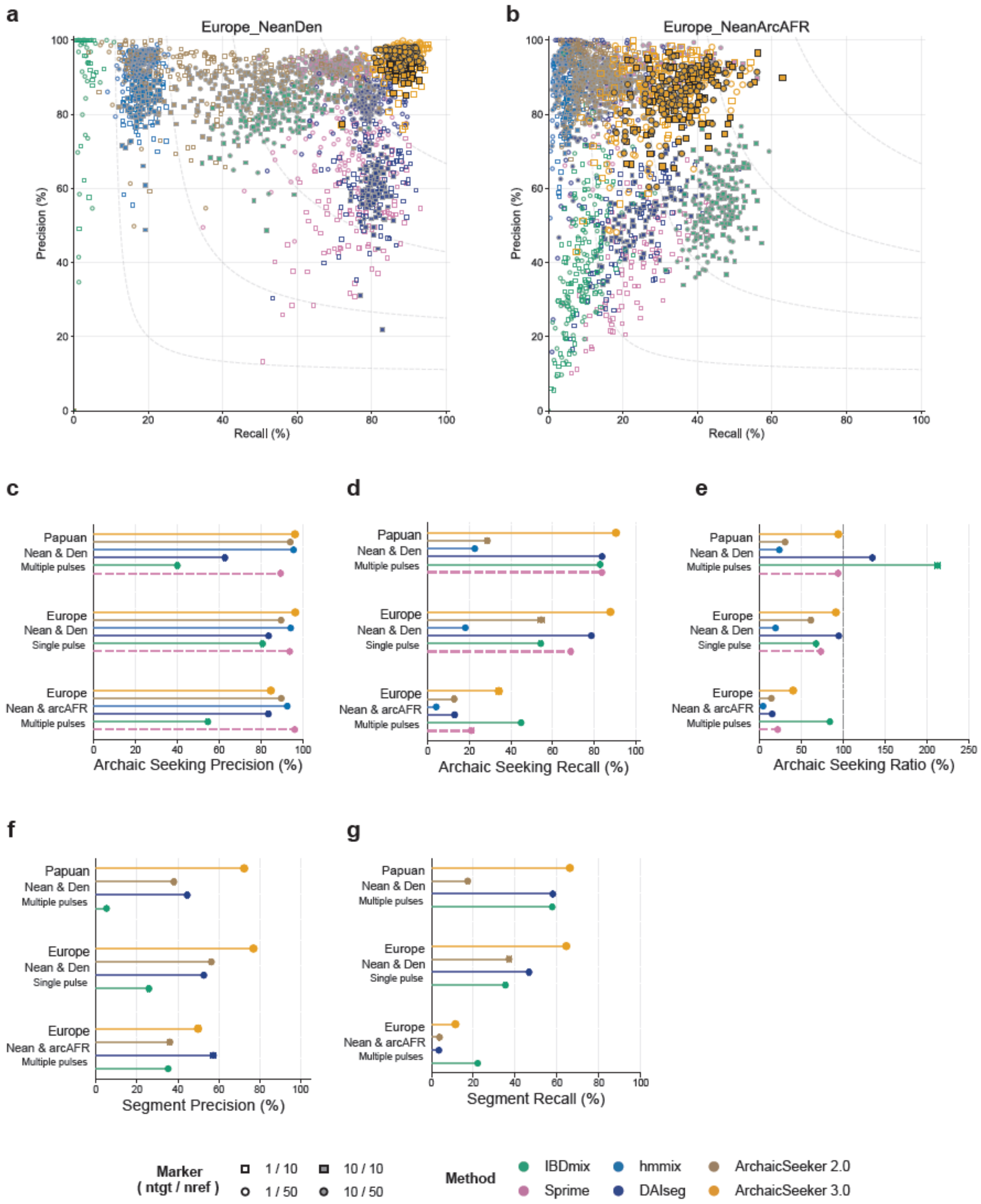


**Fig. S2.2 | Archaic-seeking performance comparison in selected scenarios.**
(a,b) Segment-level precision–recall distributions across replicates and sample-size settings for Europe_NeanDen and Europe_NeanArcAFR, respectively. Marker shape encodes ($n_{tgt}/n_{ref}$): open square, 1/10; open circle, 1/50; filled square, 10/10; filled circle, 10/50. Grey dashed curves indicate iso-F1 contours.
(c–e) Archaic-seeking precision, recall, and precision/recall ratio (%) across Papuan Nean & Den (multiple pulses), Europe Nean & Den (single pulse), and Europe Nean & arcAFR (multiple pulses).
(f,g) Segment-level precision and recall (%) for methods that output haplotype-resolved tracts.
Methods compared: IBDmix, Sprime, hmmix, DAIseg, ArchaicSeeker 2.0, and ArchaicSeeker 3.0.

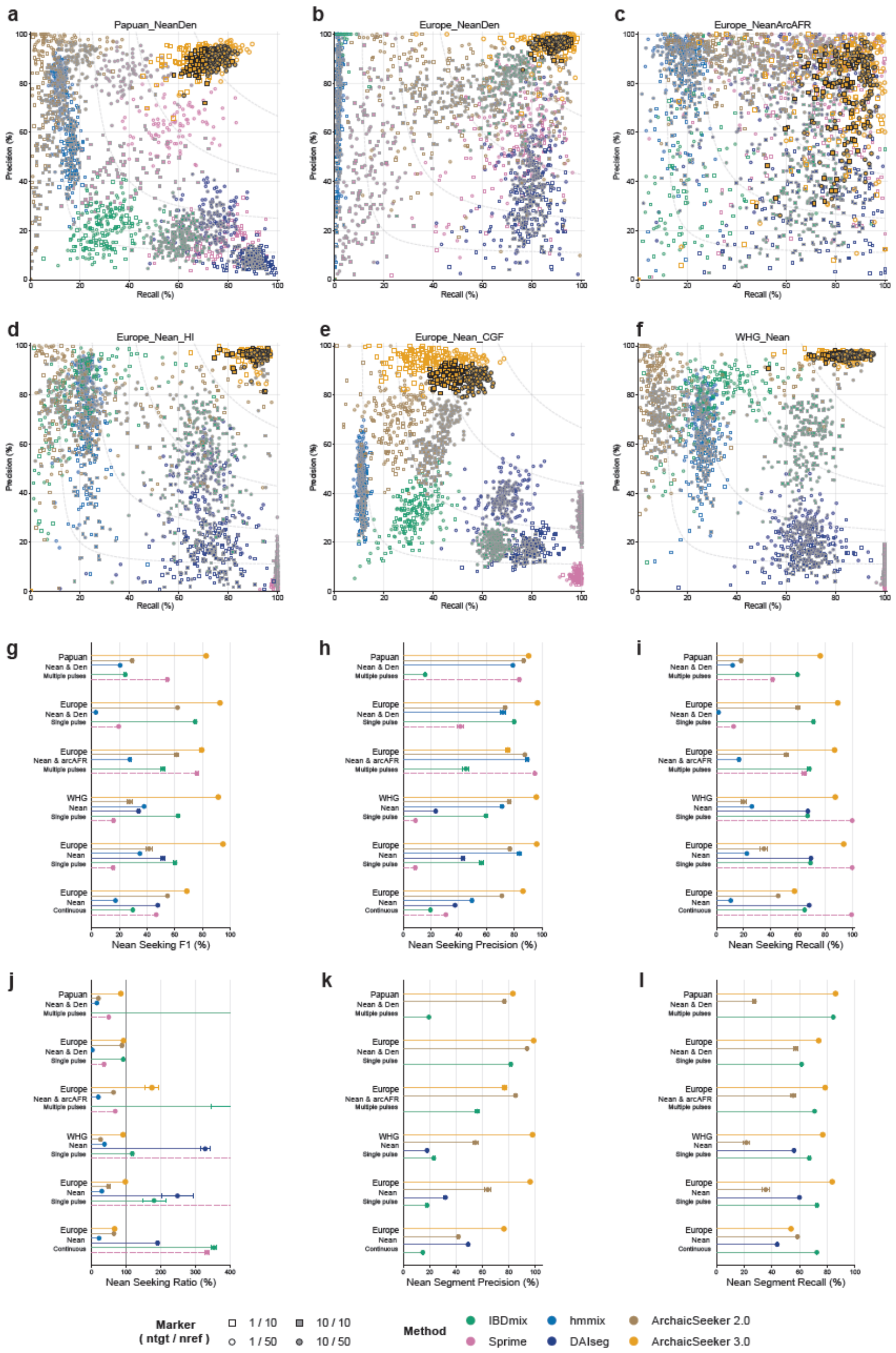


**Fig. S2.3 | Neanderthal-seeking benchmark across six demographic regimes.** (a–f) Segment-level precision–recall distributions for Neanderthal-seeking in Papuan_NeanDen, Europe_NeanDen, Europe_NeanArcAFR, Europe_Nean_HI, Europe_Nean_CGF, and WHG_Nean. Marker shapes encode ($n_{tgt}/n_{ref}$) as in Fig. S2.2; grey dashed curves show iso-F1 contours.

(g–i) Nean-seeking F1, precision, and recall (%).

(j) Nean-seeking precision/recall ratio (%).

(k,l) Segment-level precision and recall (%) for methods with directly comparable haplotype-level tract outputs.

Methods compared: IBDmix, Sprime, hmmix, DAIseg, ArchaicSeeker 2.0, and ArchaicSeeker 3.0.

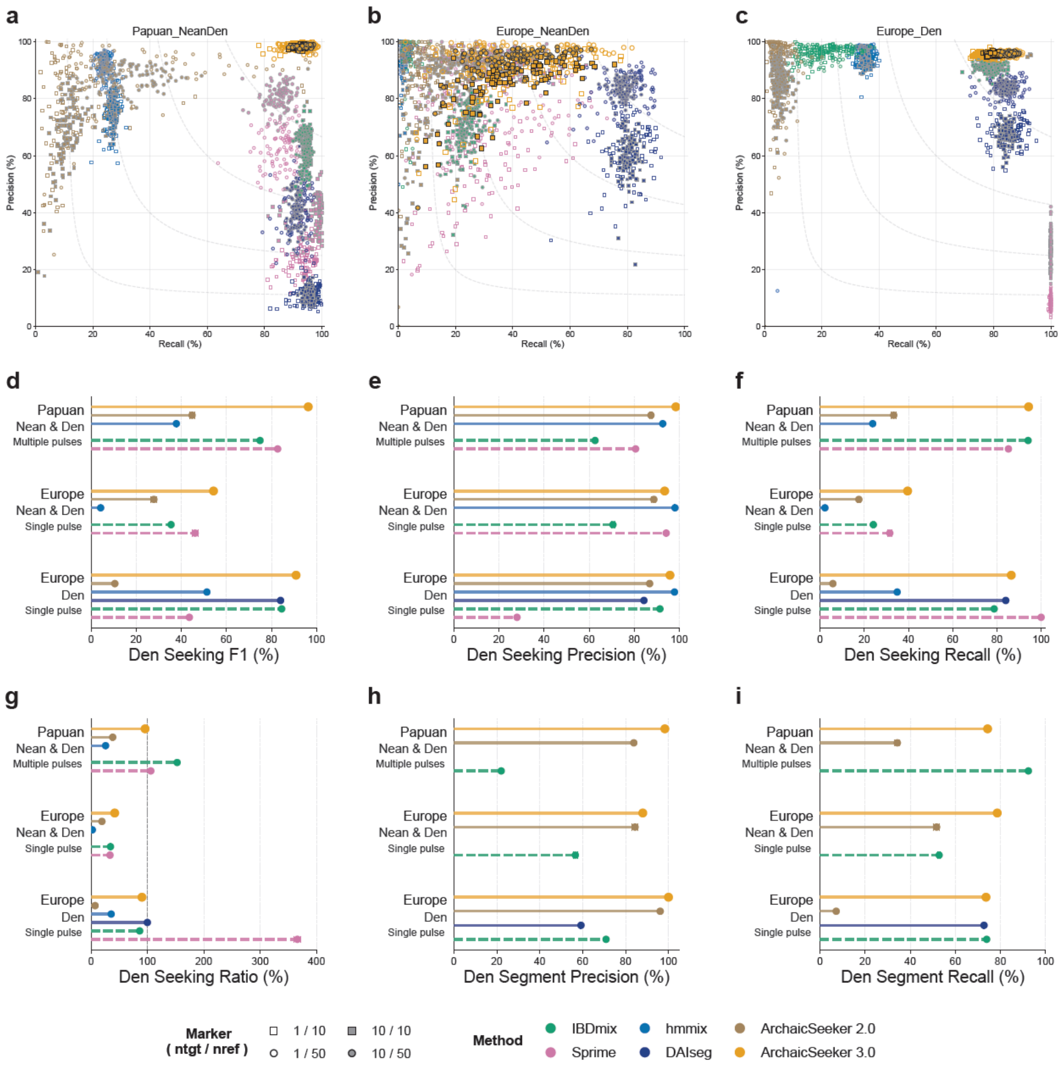


**Fig. S2.4 | Denisovan-seeking benchmark.**

(a–c) Segment-level precision–recall distributions for Den-seeking in Papuan_NeanDen, Europe_NeanDen, and Europe_Den. Marker shapes encode ($n_{tgt}/n_{ref}$) as in Fig. S2.2; grey dashed curves show iso-F1 contours.

(d–f) Den-seeking F1, precision, and recall (%).

(g) Den-seeking precision/recall ratio (%).

(h,i) Segment-level precision and recall (%) for methods with directly comparable tract outputs.

Methods compared: IBDmix, Sprime, hmmix, DAIseg, ArchaicSeeker 2.0, and ArchaicSeeker 3.0.

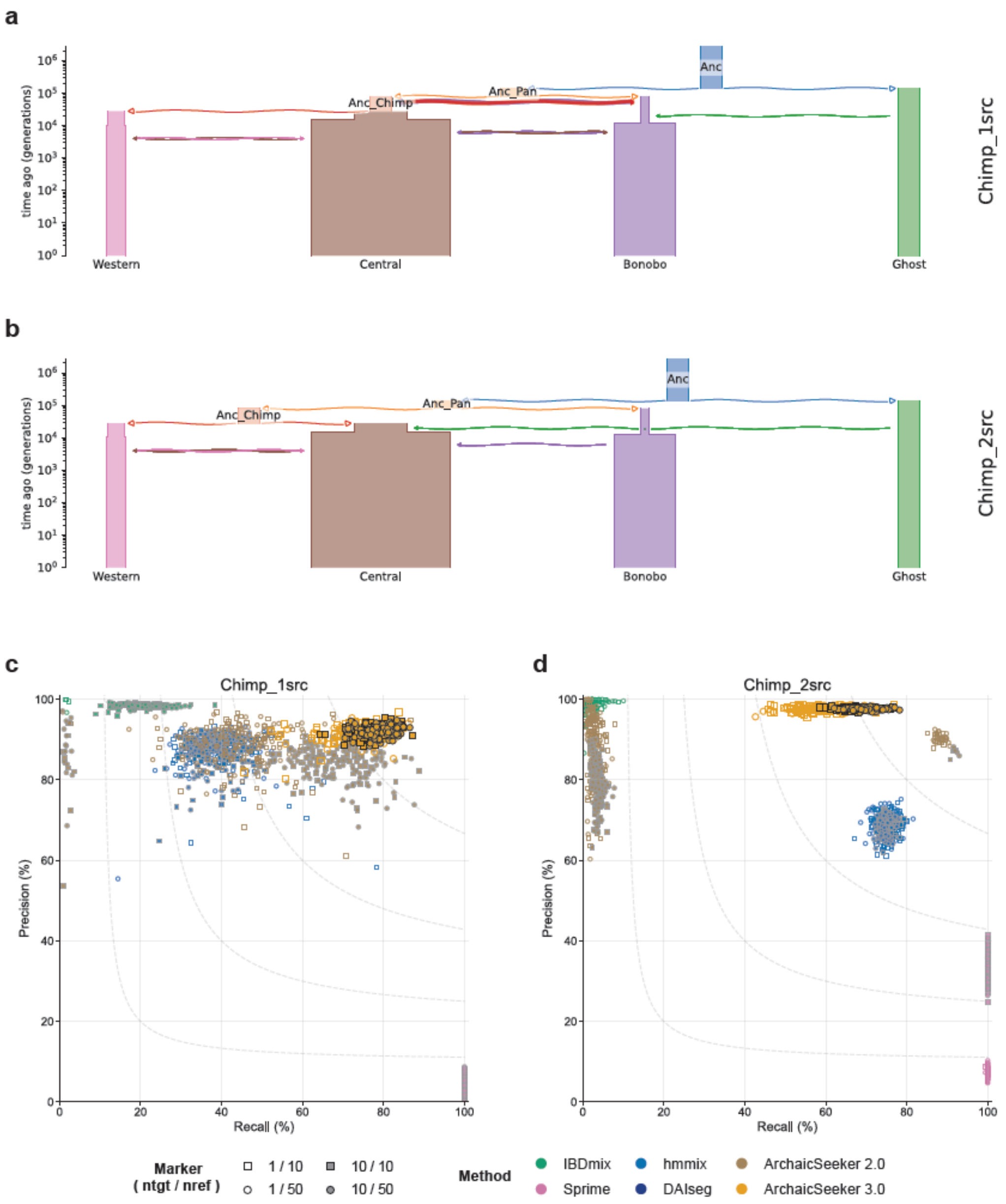


**Fig. S2.5 | Chimpanzee-lineage simulation benchmark.**
(a,b) Two chimpanzee demographic/source configurations used for simulation (Chimp_1src and Chimp_2src).
(c,d) Segment-level precision–recall distributions for archaic-seeking under Chimp_1src and Chimp_2src, respectively. Marker shapes encode ($n_{tgt}/n_{ref}$) as in Fig. S2.2; grey dashed curves indicate iso-F1 contours.
Methods compared: IBDmix, Sprime, hmmix, DAIseg, ArchaicSeeker 2.0, and ArchaicSeeker 3.0.

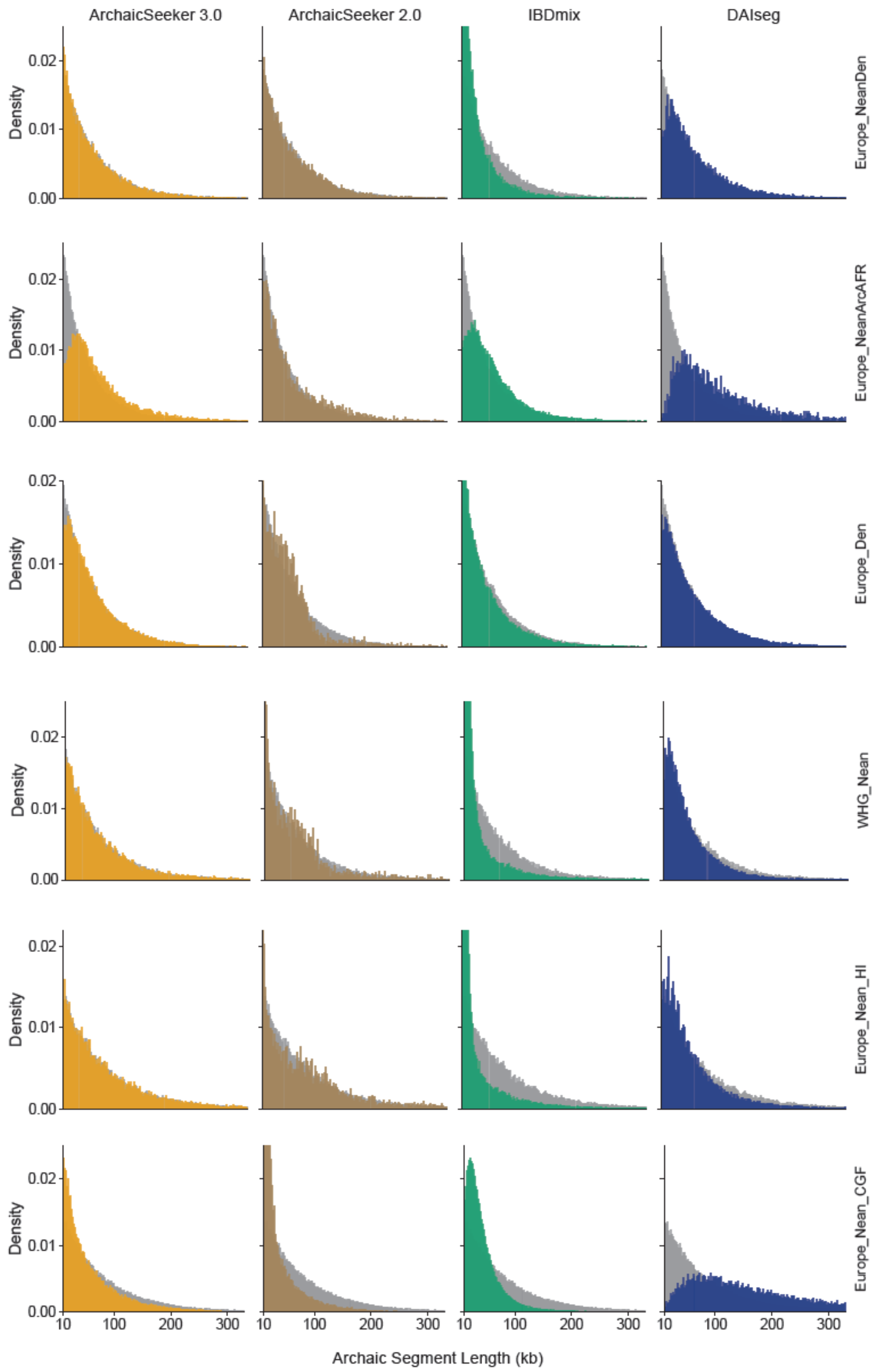


**Fig. S2.6 | Distribution of inferred archaic tract lengths across methods and scenarios.**

Histogram-based density distributions of inferred archaic segment length (kb) are shown by method (columns: ArchaicSeeker 3.0, ArchaicSeeker 2.0, IBDmix, DAIseg) and by scenario (rows: Europe_NeanDen, Europe_NeanArcAFR, Europe_Den, WHG_Nean, Europe_Nean_HI, Europe_Nean_CGF). Grey histograms indicate the simulated truth distribution in each scenario.
Only methods producing directly comparable haplotype-level tract calls are included.

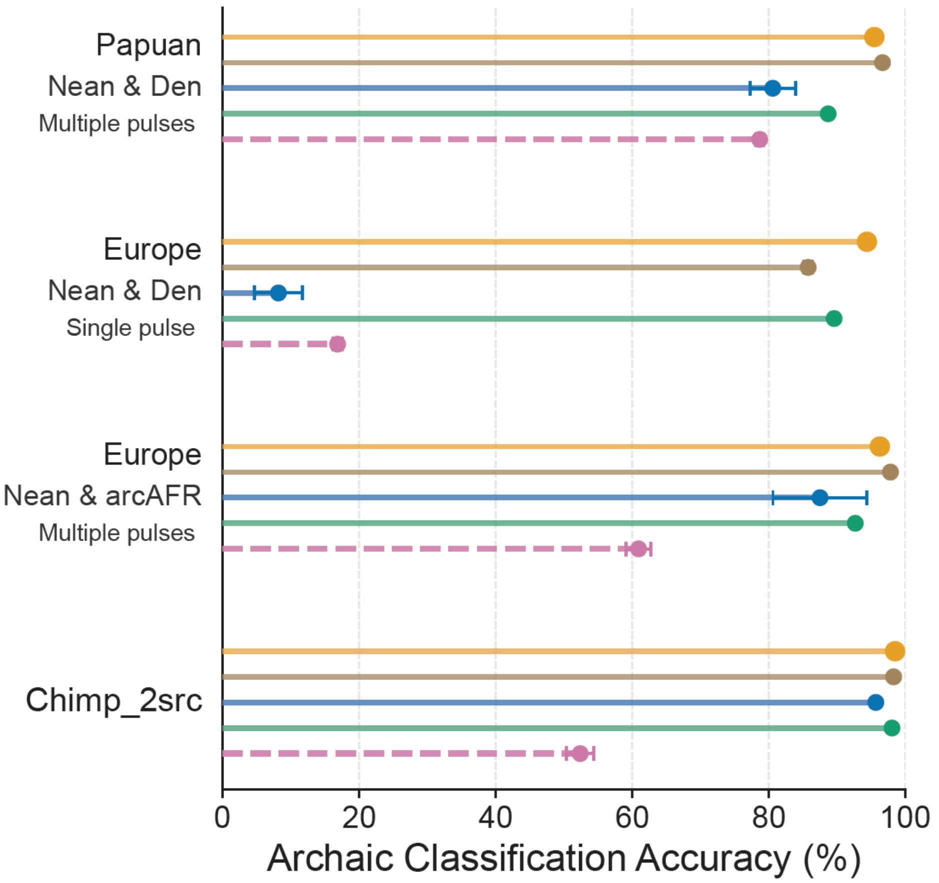


**Fig. S2.7 | Archaic-source classification accuracy across simulation settings.** Archaic classification accuracy (%) across Papuan Nean & Den (multiple pulses), Europe Nean & Den (single pulse), Europe Nean & arcAFR (multiple pulses), and Chimp_2src.

Points indicate method-level performance; horizontal error bars (where shown) denote variability across replicates.

Methods compared correspond to the benchmark set used in Fig. S2.

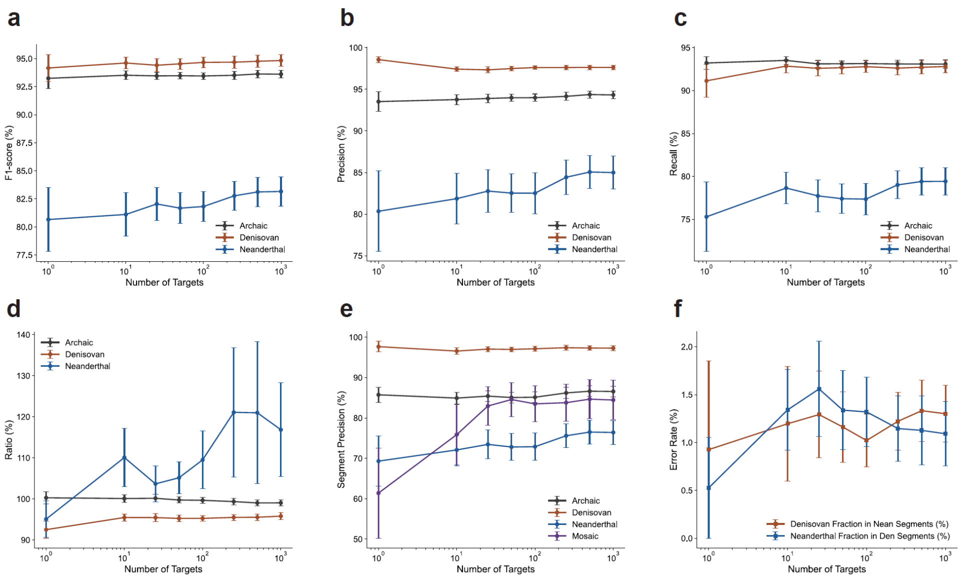


**Fig. S3.1 | Stability across target sample size ($n_{tgt}$).**

Performance of the ArchaicSeeker 3.0 across varying numbers of target samples in the evaluation cohort, with all other settings fixed as described in Methods. Panels (a–f) summarize robustness to target cohort size: (a) archaic-seeking F1 score; (b) precision; (c) recall; (d) archaic ratio (as defined in Methods); (e) segment-based precision; and (f) cross-archaic misclassification rates, quantifying errors where Denisovan segments are misclassified as Neanderthal and Neanderthal segments are misclassified as Denisovan.

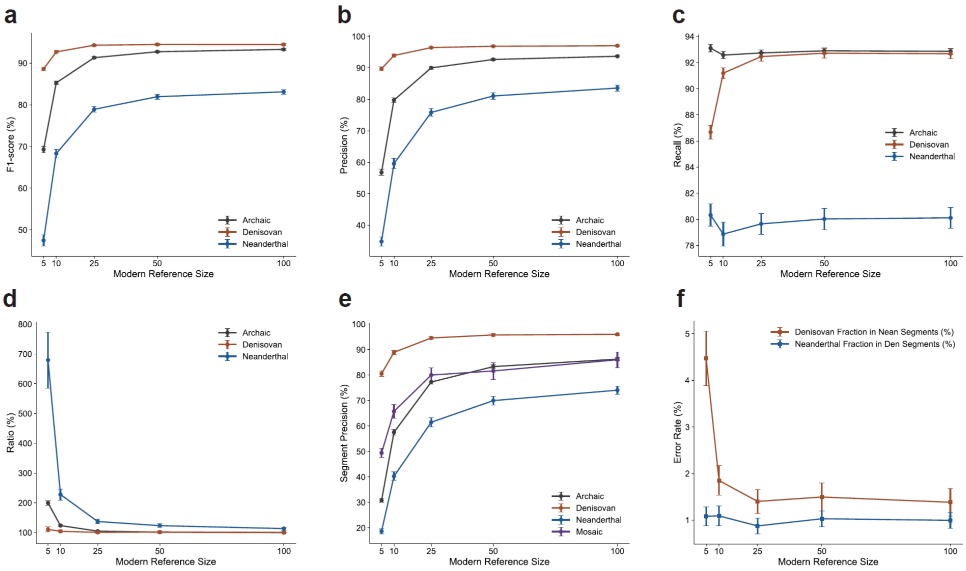


**Fig. S3.2 | Dependence on modern reference panel size ($n_{ref}$).**
Performance of the ArchaicSeeker 3.0 across varying numbers of modern reference haplotypes used for feature construction, with all other settings fixed as described in Methods. Panels (a–f) report (a) archaic-seeking F1 score, (b) precision, (c) recall, (d) archaic ratio, (e) segment-based precision, and (f) cross-archaic misclassification rates (Denisovan→Neanderthal and Neanderthal→Denisovan), illustrating how performance changes with modern reference panel size.

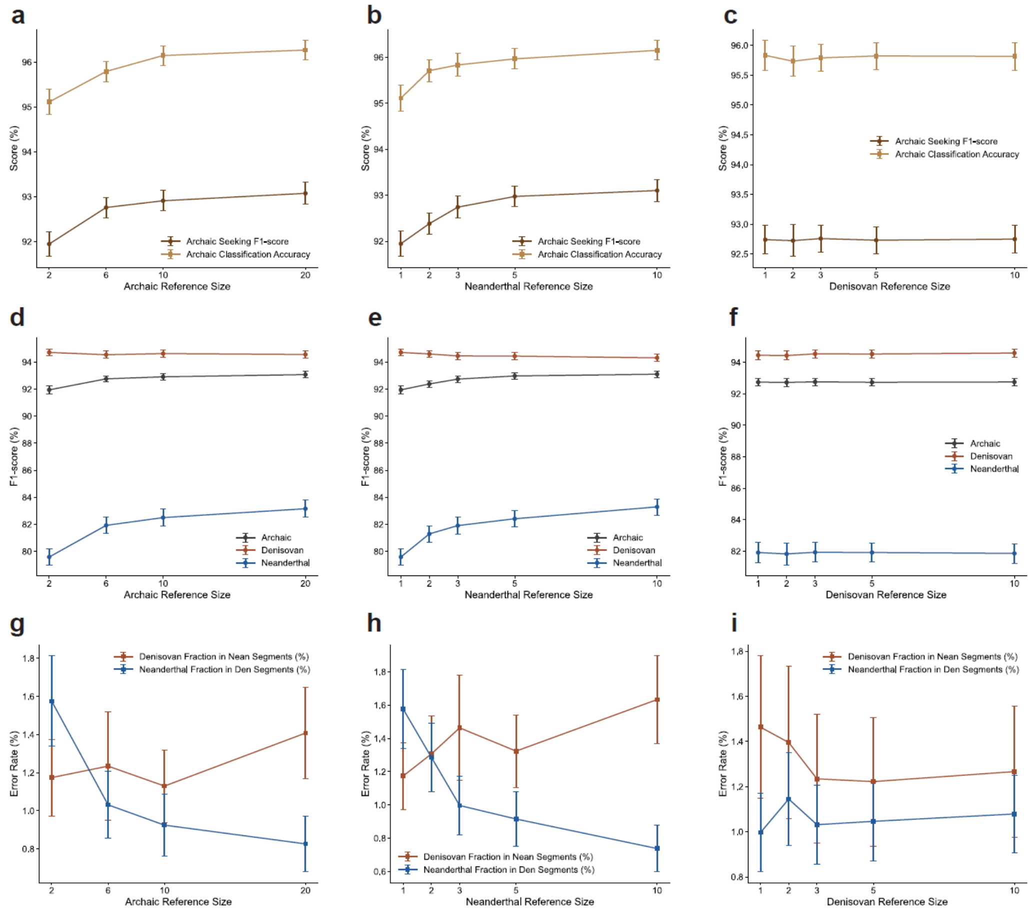


**Fig. S3.3 |** Weak dependence on archaic reference panel size.
Performance of the ArchaicSeeker 3.0 across varying numbers of archaic reference haplotypes, with all other settings fixed as described in Methods. Panels (a–f) report (a) archaic-seeking F1 score, (b) precision, (c) recall, (d) archaic ratio, (e) segment-based precision, and (f) cross-archaic misclassification rates (Denisovan→Neanderthal and Neanderthal→Denisovan). Overall, performance shows only a minor change with archaic reference depth, indicating limited impact on tract detection and source assignment.

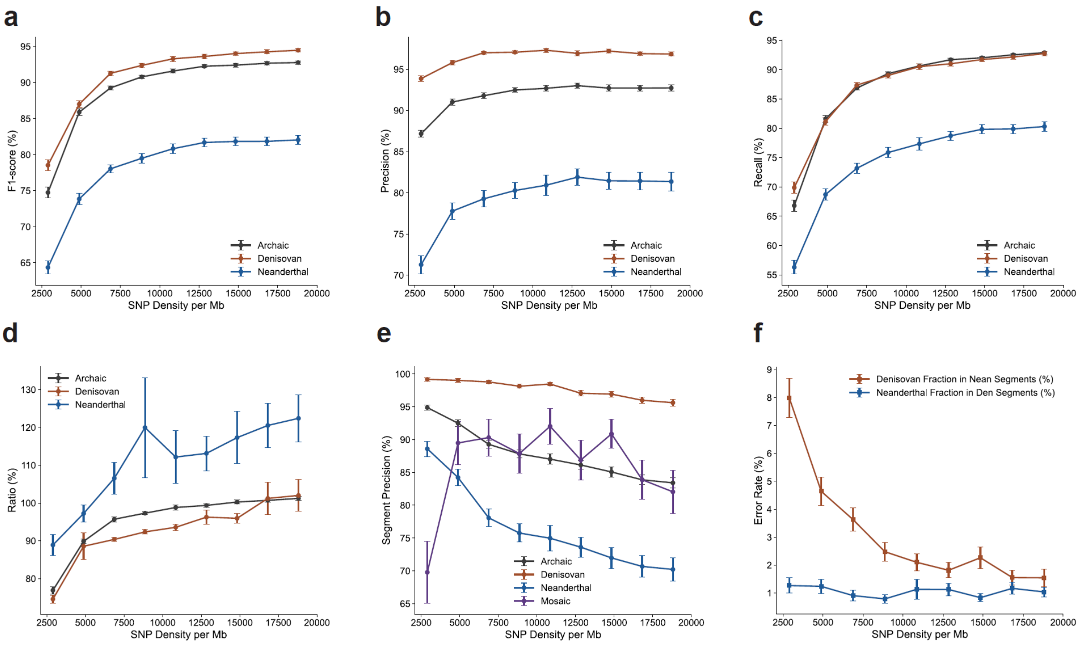


**Fig. S3.4 | Robustness to SNP marker density.**
Performance of the ArchaicSeeker 3.0 under progressively reduced SNP density in the target data, with all other settings fixed as described in Methods. Panels (a–f) report (a) archaic-seeking F1 score, (b) precision, (c) recall, (d) archaic ratio, (e) segment-based precision, and (f) cross-archaic misclassification rates (Denisovan→Neanderthal and Neanderthal→Denisovan) as marker density decreases. These analyses delineate the practical operating range of ArchaicSeeker 3.0 under sparse-marker settings.

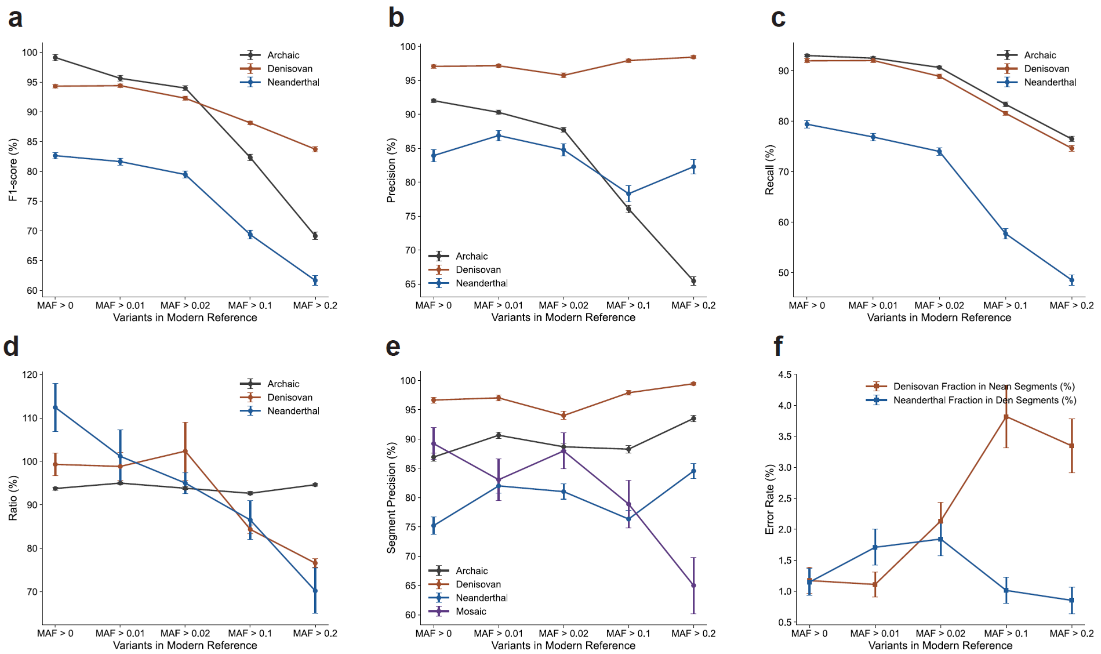


**Fig. S3.5 | Robustness to reference-frequency filtering.**

Performance of the ArchaicSeeker 3.0 after filtering variants in the modern reference data using allele-frequency criteria, with all other settings fixed as described in Methods. Panels (a–f) report (a) archaic-seeking F1 score, (b) precision, (c) recall, (d) archaic ratio , (e) segment-based precision, and (f) cross-archaic misclassification rates (Denisovan→Neanderthal and Neanderthal→Denisovan) across filtering regimes. Results assess how reference ascertainment via frequency filtering impacts tract detection and source classification.

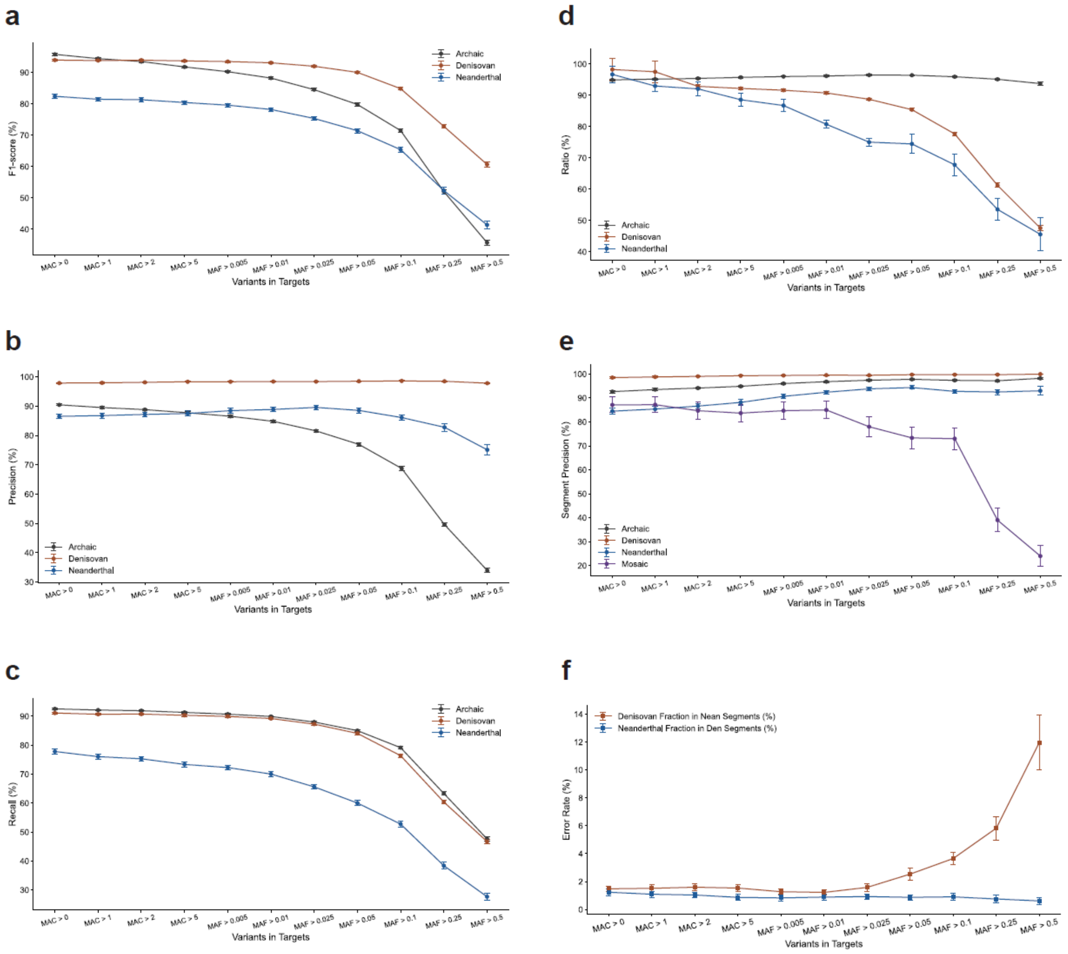


**Fig. S3.6 | Robustness to target-frequency filtering.**

Performance of the ArchaicSeeker 3.0 after filtering variants in the target data using allele-frequency criteria, with all other settings fixed as described in Methods. Panels (a–f) report (a) archaic-seeking F1 score, (b) precision, (c) recall, (d) archaic ratio , (e) segment-based precision, and (f) cross-archaic misclassification rates (Denisovan→Neanderthal and Neanderthal→Denisovan) across target-frequency thresholds. Results quantify how target ascertainment via frequency filtering impacts archaic signal recovery and source classification.

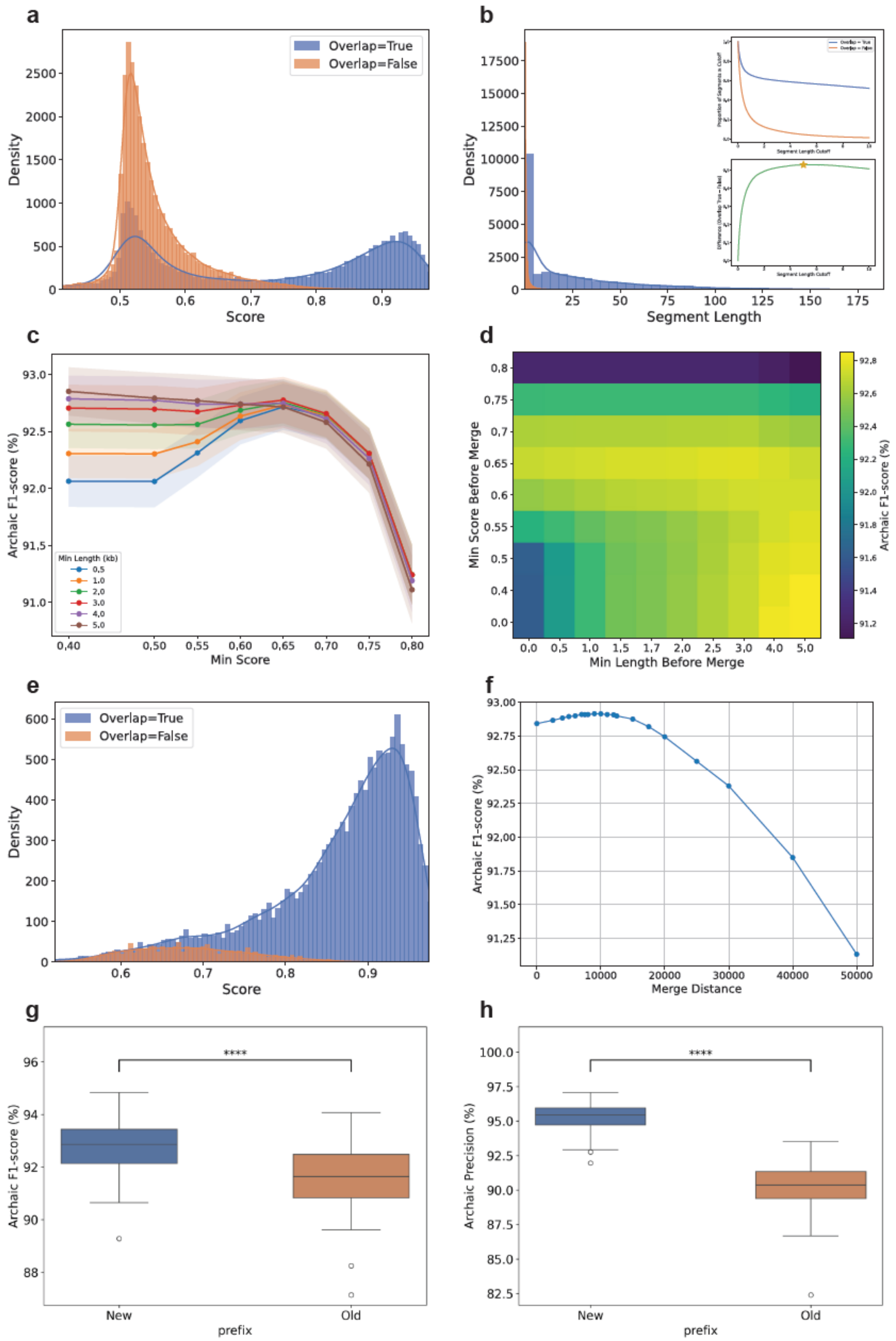


**Fig. S3.7 | Optimization of pre-merge filtering and merge parameters for ArchaicSeeker 3.0.**

Overlap = True denotes ArchaicSeeker 3.0-inferred archaic segments that overlap simulated ground-truth archaic tracts. Panels (a–b) characterize candidate segments prior to merging, showing the score distribution (a) and segment-length distribution (b) for overlapping versus non-overlapping segments. The inset in (b) summarizes, across a range of length cutoffs, the fraction of segments assigned to each group; the green curve shows the difference between groups, which peaks at ~5 kb, indicating an empirically optimal length cutoff for separating true from spurious calls. Panels (c–d) evaluate how pre-merge filtering thresholds affect overall accuracy, reporting archaic-seeking F1 as a function of minimum score and minimum length before merge (c), and as a joint grid over score and length thresholds (d). Panel (e) shows the score distributions after applying the recommended pre-merge length filter (≥5 kb), illustrating improved separation between overlapping and non-overlapping segments. Panel (f) assesses sensitivity to the merge distance parameter, reporting archaic-seeking F1 across merge distances. Panels (g–h) compare two workflows: New, applying pre-merge filtering before merging, and Old, merging first without pre-filtering. Pre-merge filtering improves both overall archaic-seeking F1 (g) and precision (h), supporting the adopted filtering-and-merge strategy in the ArchaicSeeker 3.0 pipeline.

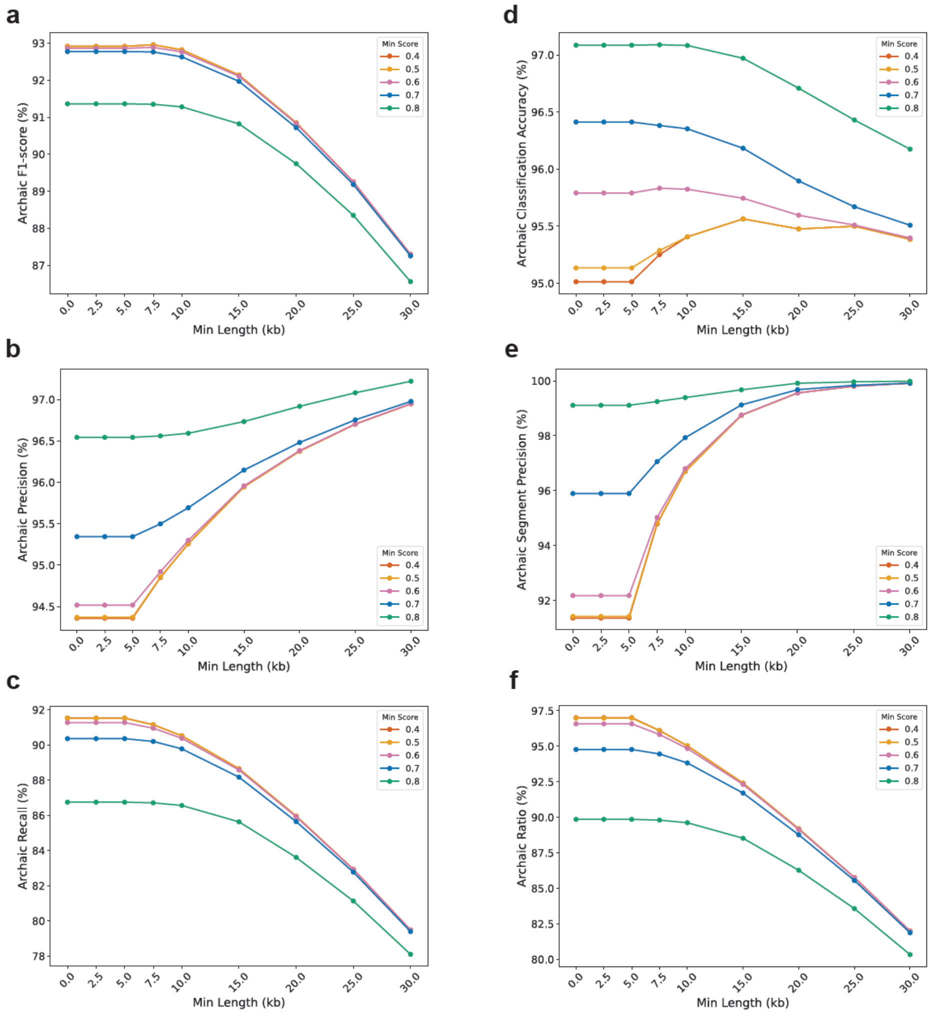


**Fig. S3.8 | post-merge filtering and its impact on performance.**
Performance of the ArchaicSeeker 3.0 pipeline under alternative post-merge filtering settings, which we provide as user-selectable options to accommodate different study designs and operating points. Using simulated data, panels (a–f) report (a) archaic-seeking F1 score, (b) precision, (c) recall, (d) archaic source-classification accuracy, (e) segment-based precision, and (f) inferred archaic ratio (defined in Methods), illustrating the trade-offs induced by different post-merge filtering parameter choices.

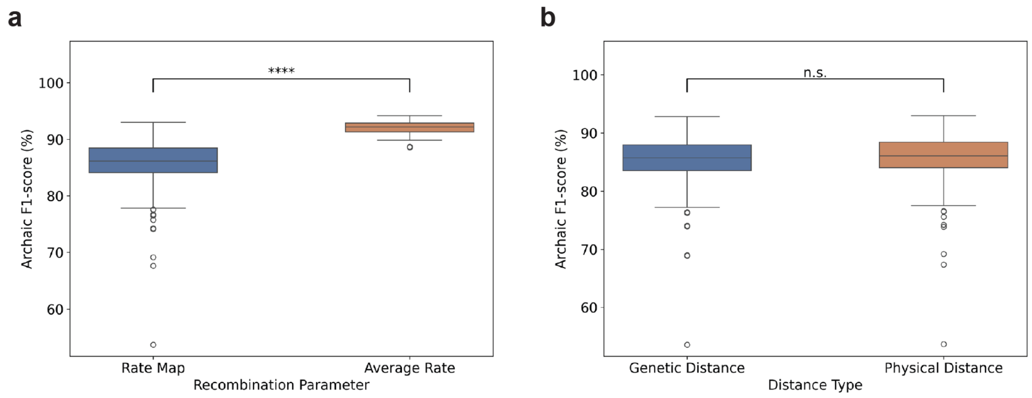


**Fig. S3.9 | Effect of recombination map specification.**

(a) Comparison of performance when simulated data are generated under a fine-scale recombination map versus a constant (genome-wide average) recombination rate.

(b) Effect of providing a fine-scale recombination map as an input to ArchaicSeeker 3.0 versus using a constant-rate approximation during preprocessing/inference.

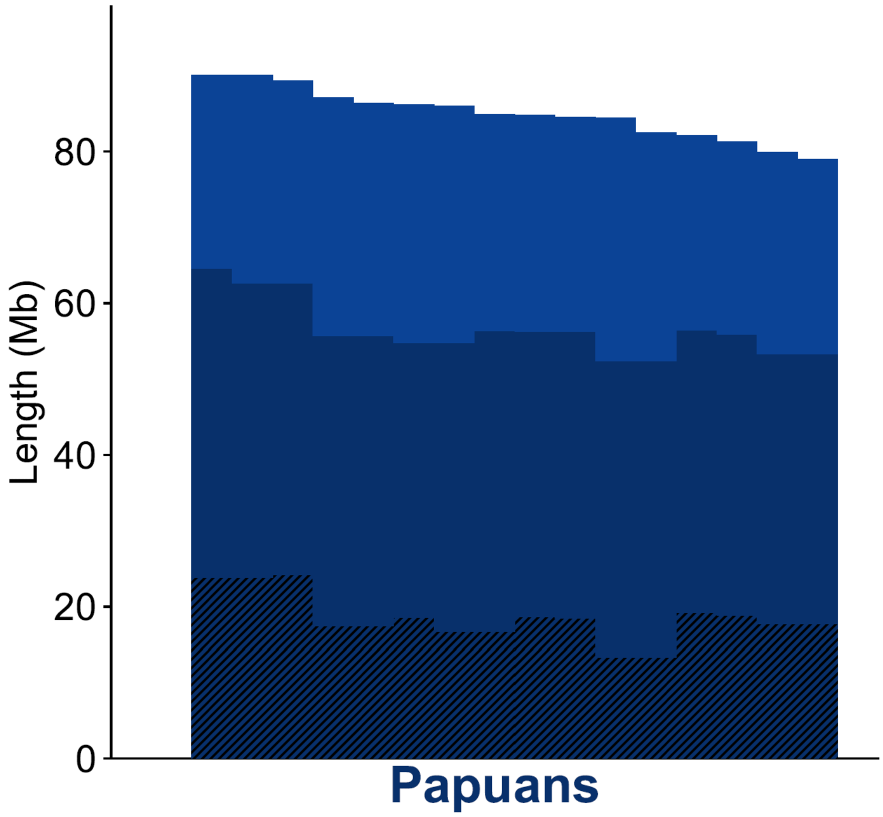


**Fig. S4.1 | Distribution of Denisovan introgressed segment lengths in Papuans inferred by ArchaicSeeker 3.0.**

Distribution of inferred Denisovan segment lengths in Papuans detected by ArchaicSeeker 3.0. The x axis shows segment length (Mb), and the y axis shows the distribution across inferred segments.

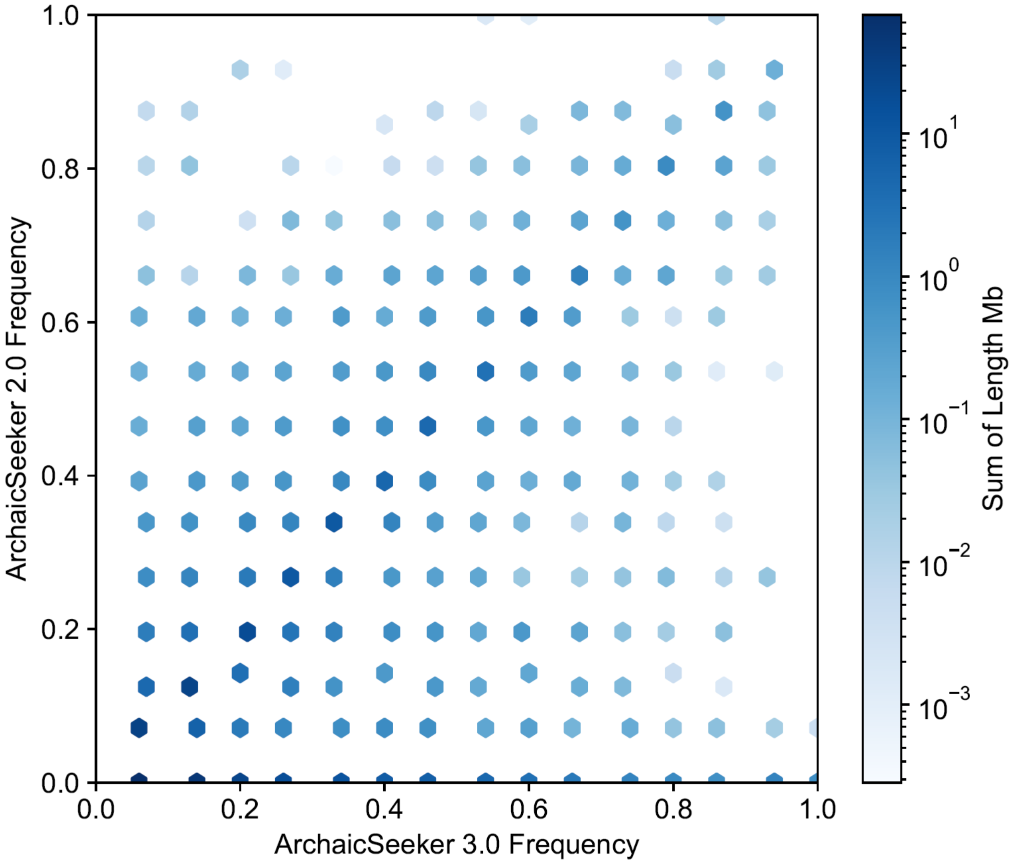


**Fig. S4.2 | Concordance of Denisovan frequency estimates between ArchaicSeeker 3.0 and ArchaicSeeker 2.0 in Papuans.**

Hexbin comparison of per-bin Denisovan introgression frequency inferred by ArchaicSeeker 3.0 (x axis) versus ArchaicSeeker 2.0 (y axis). Each hexagon summarizes bins with the same frequency pair, and color encodes the summed genomic length (Mb; log scale).

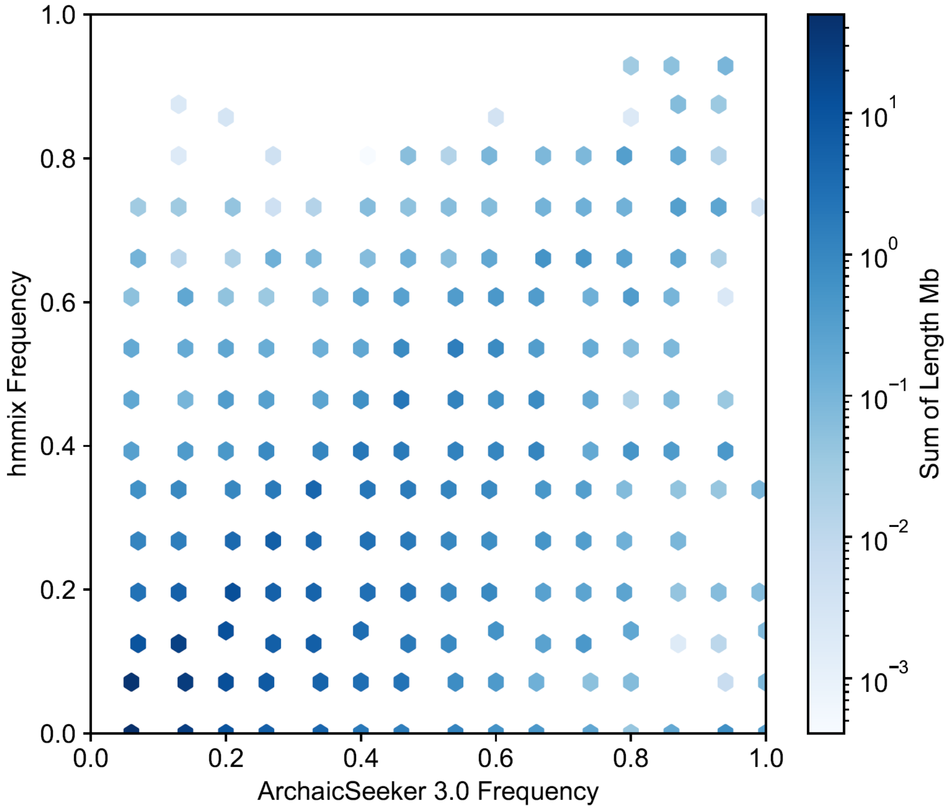


**Fig. S4.3 | Concordance of Denisovan frequency estimates between ArchaicSeeker 3.0 and hmmix in Papuans.**

Hexbin comparison of per-bin Denisovan introgression frequency inferred by ArchaicSeeker 3.0 (x axis) versus hmmix (y axis). Hexagon color represents summed genomic length (Mb; log scale), highlighting agreement and disagreement regions across the frequency space.

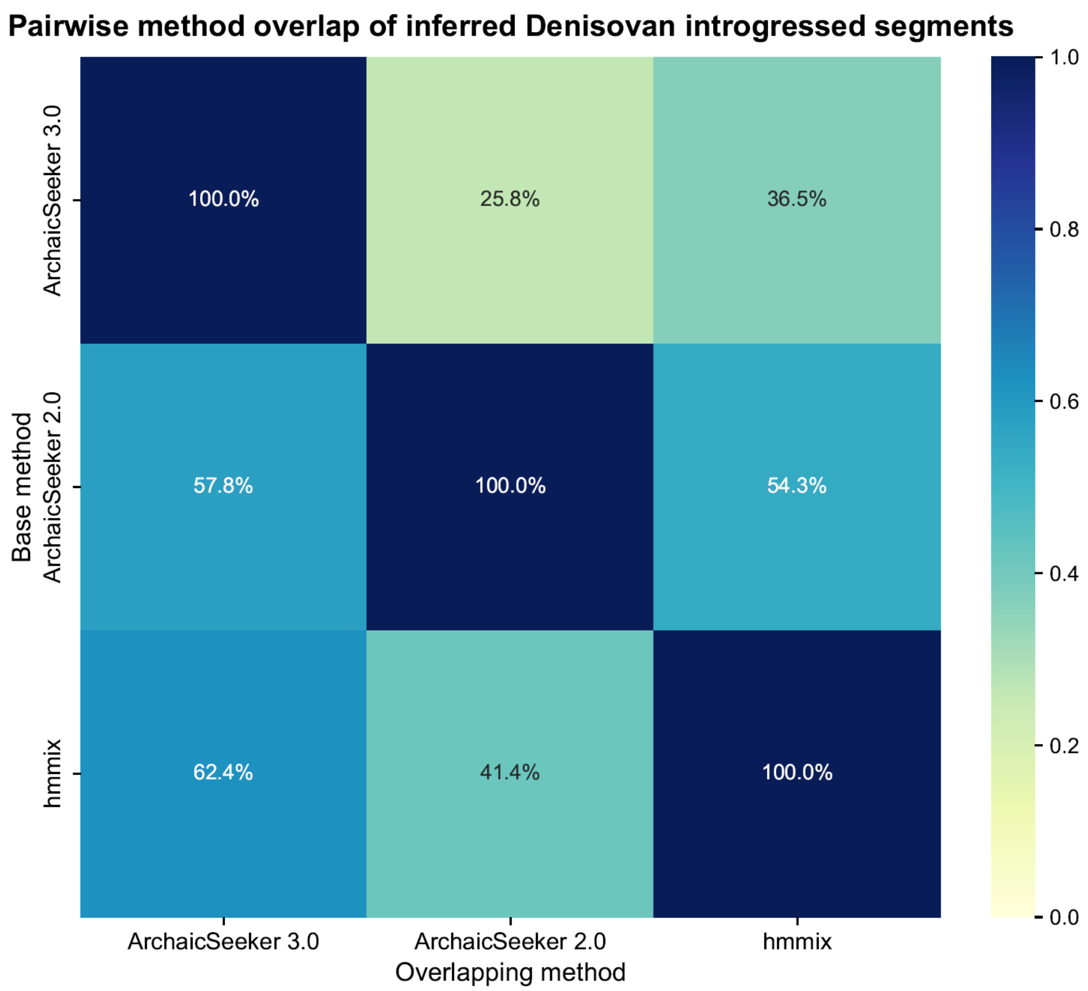


**Fig. S4.4 | Pairwise overlap heat map of inferred Denisovan introgressed segments across methods in Papuans.**

Directed pairwise overlap matrix among ArchaicSeeker 3.0, ArchaicSeeker 2.0, and hmmix. Rows denote the base method and columns denote the overlapping method; diagonal entries are 100%. Off-diagonal values are asymmetric (ArchaicSeeker 3.0→ArchaicSeeker 2.0: 25.8%, ArchaicSeeker 2.0→ArchaicSeeker 3.0: 57.8%; ArchaicSeeker 3.0→hmmix: 36.5%, hmmix→ArchaicSeeker 3.0: 62.4%; ArchaicSeeker 2.0→hmmix: 54.3%, hmmix→ArchaicSeeker 2.0: 41.4%), reflecting denominator dependence on the base method.

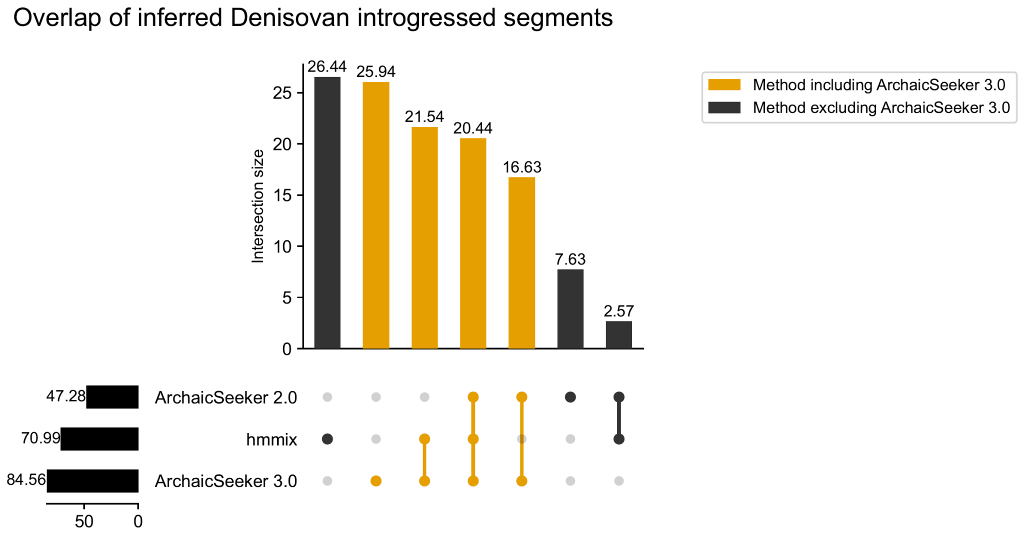


**Fig. S4.5 | UpSet analysis of overlap structure of inferred Denisovan introgressed segments in Papuans.**

UpSet plot summarizing intersection sizes among ArchaicSeeker 3.0, ArchaicSeeker 2.0, and hmmix. Vertical bars indicate intersection sizes for each method combination, and the connected-dot matrix indicates set membership. The plot explicitly contrasts intersections that include ArchaicSeeker 3.0 versus those that exclude it, and also reports method-level set sizes.

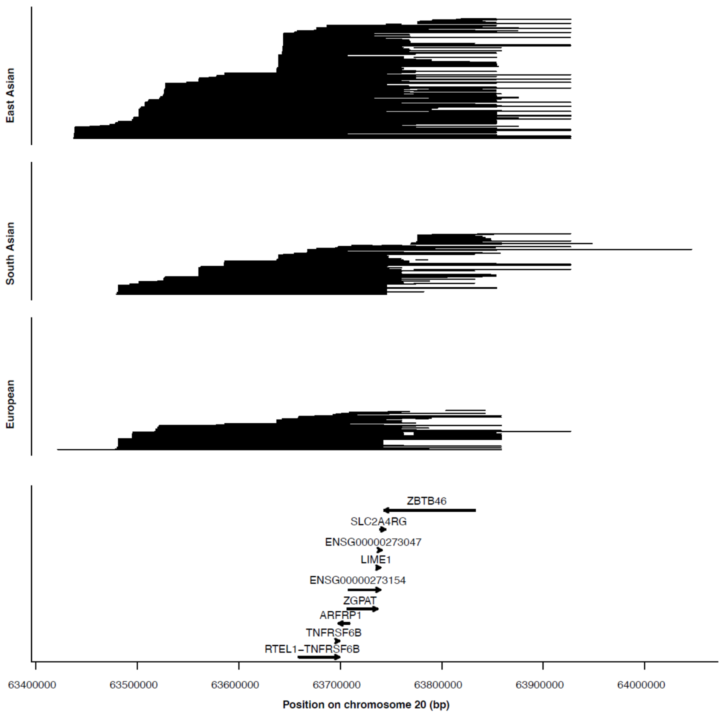


**Fig. S6.1 | AS3-specific Neanderthal-introgressed segments at the chr20 locus.**

Horizontal lines represent AS3-inferred Neanderthal-introgressed haplotypes in representative East Asian, South Asian and European populations across the plotted chr20 window surrounding chr20:63,697,389–63,769,790. The lower track shows local gene annotations, and genomic positions are shown in bp on GRCh38.

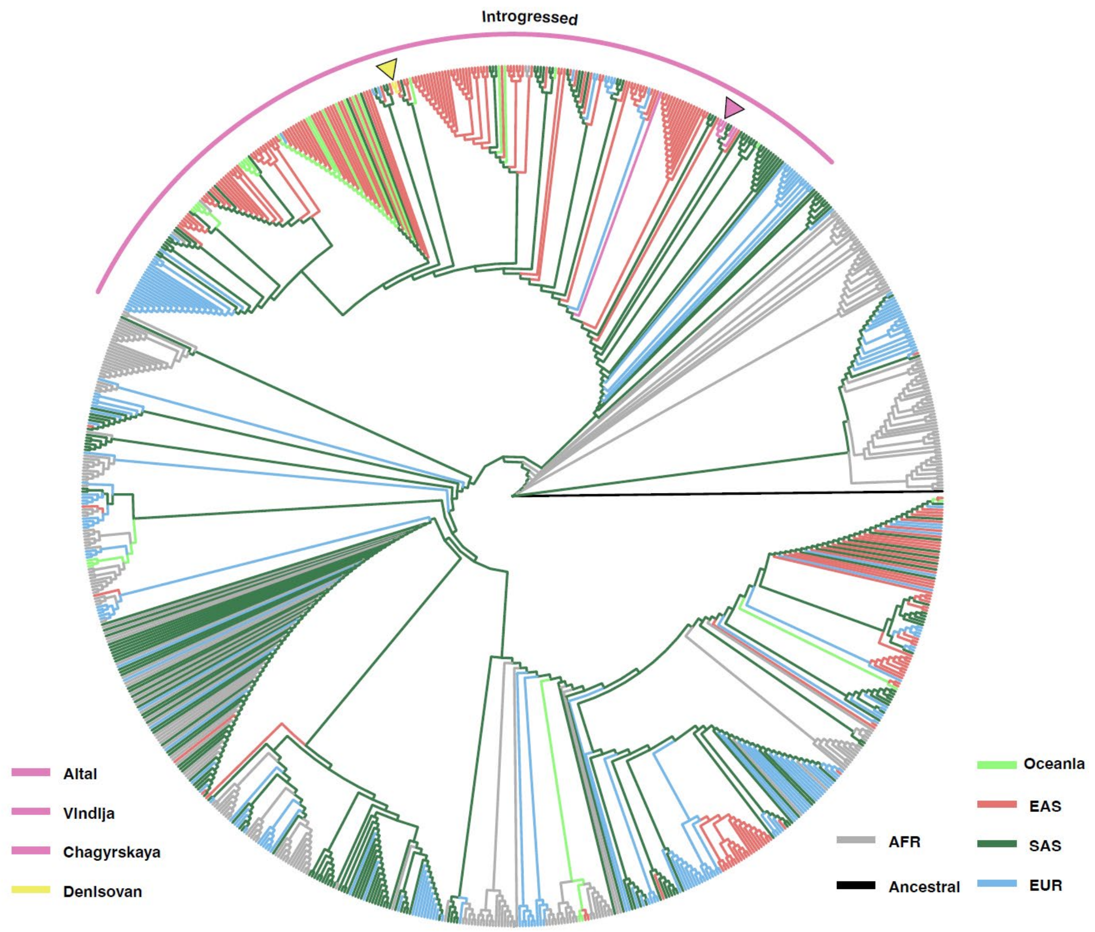


**Fig. S6.2 | Maximum-likelihood phylogeny of haplotypes across the chr20 Neanderthal-introgressed segment.**

The tree includes haplotypes from present-day populations, Neanderthal references, the Denisovan reference and the reconstructed ancestral sequence across the extended chr20 analysis window. Branch colors denote population or source groups, and the outer arc highlights the clade enriched for AS3-inferred introgressed haplotypes.

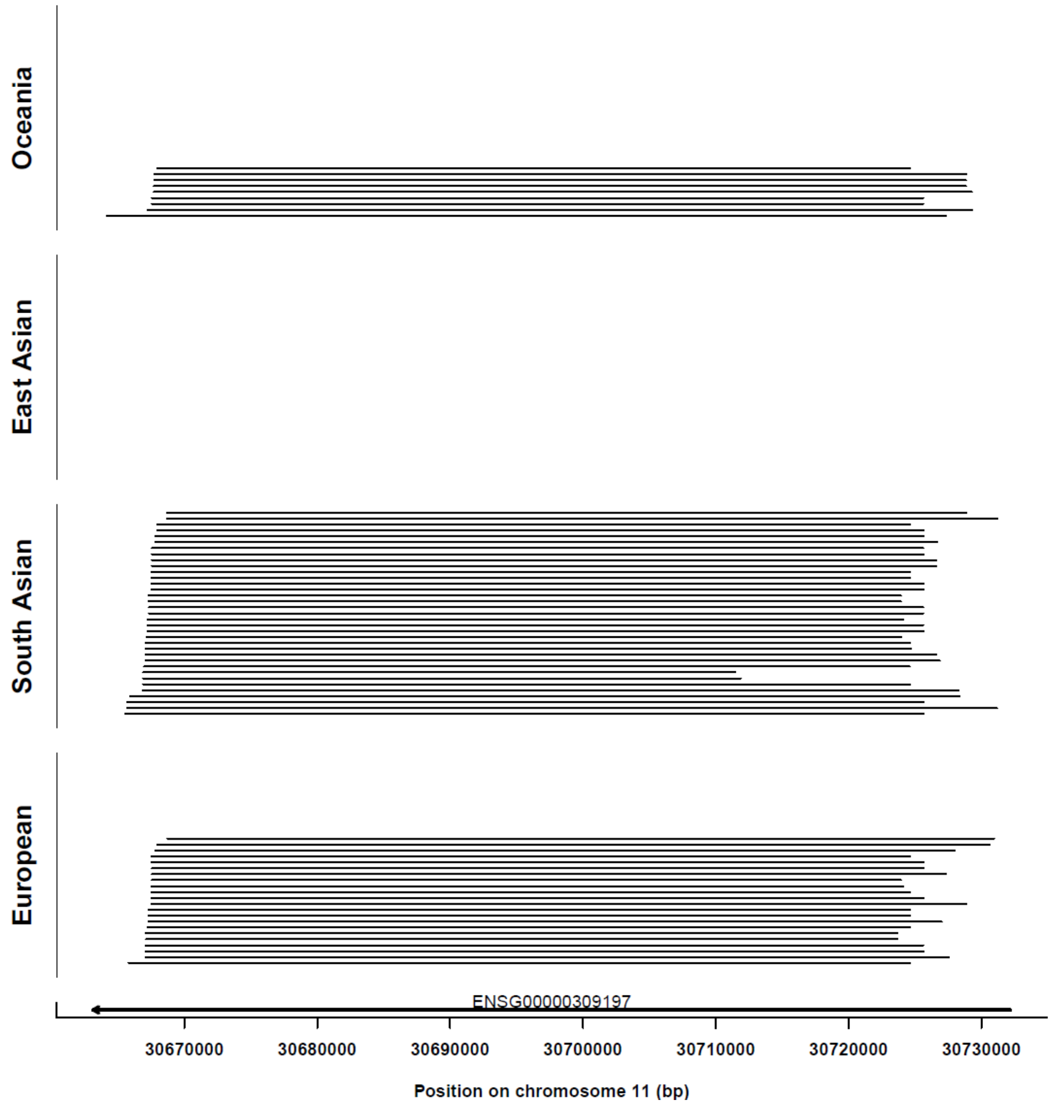


**Fig. S6.3 | AS3-specific Denisovan-introgressed segment at chr11:30,668,800–30,706,896.**

Horizontal lines represent AS3-inferred Denisovan-introgressed haplotypes across the plotted chr11 window encompassing the QC-refined introgressed segment at chr11:30,668,800–30,706,896. The lower track shows local gene annotation, including ENSG00000309197, and genomic positions are shown in bp on GRCh38.

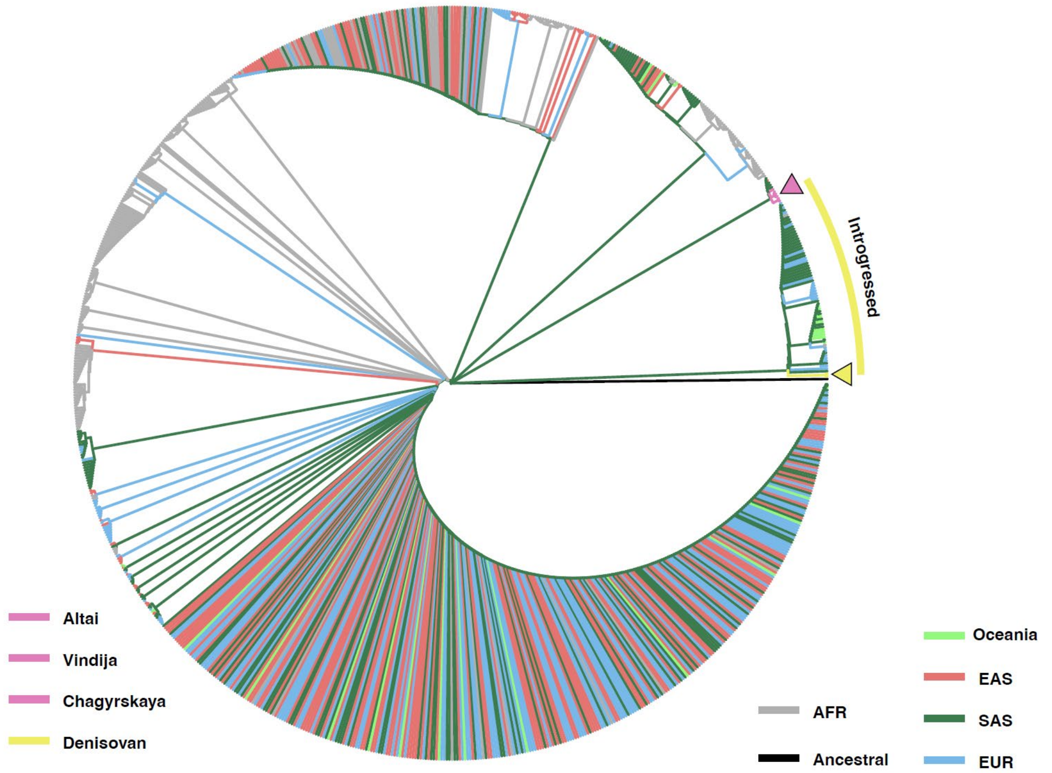


**Fig. S6.4 | Maximum-likelihood phylogeny of haplotypes across the chr11 Denisovan-introgressed segment.**

The tree includes haplotypes from present-day populations, Neanderthal references, the Denisovan reference and the reconstructed ancestral sequence across the plotted chr11 window encompassing the QC-refined introgressed segment at chr11:30,668,800–30,706,896. Branch colors denote population or source groups, and the outer arc highlights the clade enriched for AS3-inferred introgressed haplotypes.

### Data access link：

- SGDP：https://sharehost.hms.harvard.edu/genetics/reich_lab/sgdp/phased_data2021/
- EGDP
  https://web.archive.org/web/20170809212155/https://evolbio.ut.ee/CGgenomes_VCF
- HO：
  https://reich.hms.harvard.edu/index.php/datasets
- HGDPTGP：
  https://storage.googleapis.com/gcp-public-data--gnomad/release/3.1.2/vcf/genomes
- KGP2504:
  https://ftp.1000genomes.ebi.ac.uk/vol1/ftp/data_collections/1000G_2504_high_coverage/working/20201028_3202_raw_GT_with_annot/
- Sprime：
  https://data.mendeley.com/datasets/y7hyt83vxr/1
- hmmix：
  https://doi.org/10.5281/zenodo.14136628
- IBDmix：
  https://doi.org/10.5281/zenodo.14552025
- ArchaicSeeker 2.0：
  https://github.com/Shuhua-Group/ArchaicSeeker2.0/tree/master/IntrogressedSeg/KGP
- Archaic：
  https://zenodo.org/records/13368126
- Altai FilterBed:
  http://cdna.eva.mpg.de/neandertal/Vindija/FilterBed/Altai
- Chagyrskaya FilterBed
  http://cdna.eva.mpg.de/neandertal/Chagyrskaya/FilterBed/
- Vindija33.19 FilterBed
  http://cdna.eva.mpg.de/neandertal/Vindija/FilterBed/Vindija33.19
- Denisova FilterBed
  http://cdna.eva.mpg.de/neandertal/Vindija/FilterBed/Denisova
- KGP Mask：
  http://ftp.1000genomes.ebi.ac.uk/vol1/ftp/data_collections/1000_genomes_project/working/20160622_genome_mask_GRCh38/StrictMask/20160622.allChr.mask.bed

1. Kelleher, J., Etheridge, A. M. & McVean, G. Efficient Coalescent Simulation and Genealogical Analysis for Large Sa mple Sizes. *PLOS Computational Biology* **12**, e1004842 (2016). https://doi.org/10.1371/journal.pcbi.1004842
2. Kelleher, J., Thornton, K. R., Ashander, J. & Ralph, P. L. Efficient pedigree recording for fast population genetics simulation. *PLOS Computational Biology* **14**, e1006581 (2021). https://doi.org/10.1371/journal.pcbi.1006581
3. Franz, B. *et al.* Efficient ancestry and mutation simulation with msprime 1.0. *Genetics* **220**, iyab229 (2022). https://doi.org/10.1093/genetics/iyab229
4. Adrion, J. R. *et al.* A community-maintained standard library of population genetic models. *eLife* **9**, e54967 (2020). https://doi.org/10.7554/eLife.54967
5. Lauterbur, M. E. *et al.* Expanding the stdpopsim species catalog, and lessons learned for reali stic genome simulations. *eLife* **12**, RP84874 (2023). https://doi.org/10.7554/eLife.84874
6. Auton, A. *et al.* A global reference for human genetic variation. *Nature* **526**, 68-74 (2015). https://doi.org/10.1038/nature15393
7. Byrska-Bishop, M. *et al.* High-coverage whole-genome sequencing of the expanded 1000 Genomes Pro ject cohort including 602 trios. *Cell* **185**, 3426-3440.e3419 (2022). https://doi.org/10.1016/j.cell.2022.08.004
8. Hofmeister, R. J., Ribeiro, D. M., Rubinacci, S. & Delaneau, O. Accurate rare variant phasing of whole-genome and whole-exome sequenci ng data in the UK Biobank. *Nature Genetics* **55**, 1243-1249 (2023). https://doi.org/10.1038/s41588-023-01415-w
9. Browning, S. R., Browning, B. L., Zhou, Y., Tucci, S. & Akey, J. M. Analysis of Human Sequence Data Reveals Two Pulses of Archaic Denisovan Admixture. *Cell* **173**, 53-61.e59 (2018). https://doi.org/10.1016/j.cell.2018.02.031
10. skov, I. (Zenodo, 2024).
11. Liang, S.-A. *et al.* (Zenodo, 2024).
12. Yuan, K. *et al.* Refining models of archaic admixture in Eurasia with ArchaicSeeker 2.0. *Nature Communications* **12**, 6232 (2021). https://doi.org/10.1038/s41467-021-26503-5
13. Zhang, R., Yuan, K. & Xu, S. Detecting archaic introgression and modeling multiple-wave admixture w ith ArchaicSeeker 2.0. *STAR Protocols* **3**, 101314 (2022). https://doi.org/10.1016/j.xpro.2022.101314
14. Zhou, Y. & Browning, S. R. Protocol for detecting introgressed archaic variants with SPrime. *STAR Protocols* **2**, 100550 (2021). https://doi.org/10.1016/j.xpro.2021.100550
15. Zhao, H. *et al.* CrossMap: a versatile tool for coordinate conversion between genome assemblies. *Bioinformatics* **30**, 1006-1007 (2014). https://doi.org/10.1093/bioinformatics/btt730
16. Chen, L., Wolf, A. B., Fu, W., Li, L. & Akey, J. M. Identifying and Interpreting Apparent Neanderthal Ancestry in African Individuals. *Cell* **180**, 677-687.e616 (2020). https://doi.org/10.1016/j.cell.2020.01.012
17. Minh, B. Q. *et al.* IQ-TREE 2: New Models and Efficient Methods for Phylogenetic Inference in the Genomic Era. *Molecular Biology and Evolution* **37**, 1530-1534 (2020). https://doi.org/10.1093/molbev/msaa015
18. Xu, S. *et al.* Ggtree: A serialized data object for visualization of a phylogenetic tree and annotation data. *iMeta* **1**, e56 (2022). https://doi.org/https://doi.org/10.1002/imt2.56

19 Bergström, A. *et al.* Insights into human genetic variation and population history from 929 diverse genomes. *Science* **367** (2020). https://doi.org/10.1126/science.aay5012

20 Fan, S. *et al.* African evolutionary history inferred from whole genome sequence data of 44 indigenous African populations. *Genome Biology* **20**, 82 (2019). https://doi.org/10.1186/s13059-019-1679-2

21 Mallick, S. *et al.* The Simons Genome Diversity Project: 300 genomes from 142 diverse popu lations. *Nature* **538**, 201-206 (2016). https://doi.org/10.1038/nature18964

22 Pagani, L. *et al.* Genomic analyses inform on migration events during the peopling of Eurasia. *Nature* **538**, 238-242 (2016). https://doi.org/10.1038/nature19792

23 Lazaridis, I. *et al.* Ancient human genomes suggest three ancestral populations for present- day Europeans. *Nature* **513**, 409-413 (2014). https://doi.org/10.1038/nature13673

24 Patterson, N. *et al.* Ancient Admixture in Human History. *Genetics* **192**, 1065-1093 (2012). https://doi.org/10.1534/genetics.112.145037

25 Koenig, Z. *et al.* A harmonized public resource of deeply sequenced diverse human genomes. *Genome Research* **34**, 796-809 (2024). https://doi.org/10.1101/gr.278378.123

26 Chen, S. *et al.* A genomic mutational constraint map using variation in 76,156 human genomes. *Nature* **625**, 92-100 (2024). https://doi.org/10.1038/s41586-023-06045-0

27 Rubinacci, S., Hofmeister, R. J., Sousa da Mota, B. & Delaneau, O. Imputation of low-coverage sequencing data from 150,119 UK Biobank gen omes. *Nature Genetics* **55**, 1088-1090 (2023). https://doi.org/10.1038/s41588-023-01438-3

28 Danecek, P. *et al.* Twelve years of SAMtools and BCFtools. *GigaScience* **10** (2021). https://doi.org/10.1093/gigascience/giab008

29 Rubinacci, S., Ribeiro, D. M., Hofmeister, R. J. & Delaneau, O. Efficient phasing and imputation of low-coverage sequencing data using large reference panels. *Nature Genetics* **53**, 120-126 (2021). https://doi.org/10.1038/s41588-020-00756-0

30 Skov, L. (Zenodo, 2024).

31 Prüfer, K. *et al.* The complete genome sequence of a Neanderthal from the Altai Mountains. *Nature* **505**, 43-49 (2014). https://doi.org/10.1038/nature12886

32 Prüfer, K. *et al.* A high-coverage Neandertal genome from Vindija Cave in Croatia. *Science* **358**, 655-658 (2017). https://doi.org/10.1126/science.aao1887

33 Mafessoni, F. *et al.* A high-coverage Neandertal genome from Chagyrskaya Cave. *Proc Natl Acad Sci U S A* **117**, 15132-15136 (2020). https://doi.org/10.1073/pnas.2004944117

34 Meyer, M. *et al.* A High-Coverage Genome Sequence from an Archaic Denisovan Individual. *Science* **338**, 222-226 (2012). https://doi.org/doi:10.1126/science.1224344
