## Supplementary Tables for "*ArchaicSeeker* 3.0: A deep-learning framework for scalable, haplotype-resolved inference of archaic introgression"

**Table S1. Parameters for the Bonobo-Ghost model**

| Description | Value | Unit |
|---|---|---|
| reference population | Western chimpanzees | |
| target population | Bonobo | |
| source population | Ghost | |
| ancestral pop. size | 10000 | |
| ghost pop. Size | 10000 | |
| ancestral bonobo-chimpanzee pop. size | 11600 | |
| ancestral common chimpanzee pop. size | 10200 | |
| ancestral bonobo pop. size | 3700 | |
| ancestral central chimpanzee pop. size | 24900 | |
| ancestral western chimpanzee pop. size | 8000 | |
| current bonobo pop. size | 29100 | |
| current central chimpanzee pop. size | 65900 | |
| current western chimpanzee pop. size | 9200 | |
| ghost/other split time | 3500 | kya |
| bonobo-common chimpanzee start migration | 1500 | kya |
| bonobo-common chimpanzee stop migration | 1200 | kya |
| bonobo/chimpanzee split time | 1990 | kya |
| central/western chimpanzee split time | 700 | kya |
| time of ghost → bonobo gene flow | 500 | kya |
| time of central chimpanzee resize | 379 | kya |
| time of bonobo resize | 308 | kya |
| time of western chimpanzee resize | 261 | kya |
| time of bonobo <-> central chimpanzee gene flow | 155.05 | kya |
| time of western <-> central chimpanzee gene flow | 100.1 | kya |
| generation time | 25 | years |
| ghost → bonobo admixture proportion | 2 | % |
| bonobo → central chimpanzee admixture proportion | 0.125 | % |
| central chimpanzee → bonobo admixture proportion | 0.1 | % |
| central chimpanzee → western chimpanzee admixture | 1.5 | % |
| western chimpanzee → central chimpanzee admixture | 0.5 | % |
| migration rate between bonobo and ancestral | $1\times10^{-7}$ | per generation |
| sampling time of source genome (Ghost) | 0 | kya |
| mutation rate | $1.2\times10^{-8}$ | per base per generation |
| recombination rate | $0.7\times10^{-8}$ | per base per generation |

**Table S2. Parameters for the Human-Neanderthal model**

| Description | Value | Unit |
|---|---|---|
| reference population | YRI | |
| target population | CEU | |
| source population | Neanderthal | |
| ancestral pop. size | 18500 | |
| Neanderthal pop. size | 3400 | |
| YRI pop. size | 27000 | |
| CEU bottleneck pop. size | 1080 | |
| CEU growth-start pop. size | 1450 | |
| CEU current pop. size | 13377 | |
| CEU growth rate | 0.00202 | |
| CEU time at growth start | 31.9 | kya |
| Nea/other split time | 550 | kya |
| CEU/YRI split time | 65.7 | kya |
| time of Nea → CEU gene flow | 55 | kya |
| generation time | 29 | years |
| Nea → CEU admixture proportion | 2.25 | % |
| sampling time of source genome (Neanderthal) | 47650 | years |
| mutation rate | $1.29\times10^{-8}$ | per base per generation |
| recombination rate | $1\times10^{-8}$ | per base per generation |

**Table S3. Parameters for the Human-Denisovan model**

| Description | Value | Unit |
|---|---|---|
| reference population | Africa | |
| target population | Europe | |
| source population | Den | |
| Africa start size | 18296 | |
| Africa bottleneck size | 7000 | |
| Africa current size | 27122 | |
| Den start size | 5000 | |
| Europe start size | 250 | |
| Europe grow up size 1 | 5000 | |
| Europe bottleneck size | 1305 | |
| Europe grow up size 2 | 5000 | |
| Europe current size | 3899 | |
| Asia start size | 5054 | |
| Africa bottleneck start time | 657 | kya |
| Africa grow up start time | 62.04 | kya |
| Den/Africa split time | 657 | kya |
| Europe/Africa split time | 62.04 | kya |
| Europe grow up start time 1 | 59.14 | kya |
| Europe bottleneck start time | 57.1 | kya |
| Europe grow up start time 2 | 54.2 | kya |
| Europe migration to Asia start time | 42 | kya |
| Europe/Asia split time | 42 | kya |
| time of Den → Europe gene flow | 54 | kya |
| generation time | 29 | years |
| Den → Europe admixture proportion | 5 | % |
| sampling time of source genome (Denisovan) | 57913 | years |
| mutation rate | $1.2\times10^{-8}$ | per base per generation |
| recombination rate | $1.2\times10^{-8}$ | per base per generation |

**Table S4. Parameters for the Chimpanzee-Ghost-Bonobo model**

| Description | Value | Unit |
|---|---|---|
| reference population | Western chimpanzees | |
| target population | Central chimpanzees | |
| source population 1 | Ghost | |
| source population 2 | Bonobo | |
| ancestral pop. size | 10000 | |
| ghost pop. Size | 10000 | |
| ancestral bonobo-chimpanzee pop. size | 11600 | |
| ancestral common chimpanzee pop. size | 10200 | |
| ancestral bonobo pop. size | 3700 | |
| ancestral central chimpanzee pop. size | 24900 | |
| ancestral western chimpanzee pop. size | 8000 | |
| current bonobo pop. size | 29100 | |
| current central chimpanzee pop. size | 65900 | |
| current western chimpanzee pop. size | 9200 | |
| ghost/other split time | 3500 | kya |
| bonobo-common chimpanzee start migration | 1500 | kya |
| bonobo-common chimpanzee stop migration | 1200 | kya |
| bonobo/chimpanzee split time | 1990 | kya |
| central/western chimpanzee split time | 700 | kya |
| time of ghost → central chimpanzee gene flow | 500 | kya |
| time of bonobo → central chimpanzee gene flow | 155.05 | kya |
| time of central chimpanzee resize | 379 | kya |
| time of bonobo resize | 308 | kya |
| time of western chimpanzee resize | 261 | kya |
| time of western <-> central chimpanzee gene flow | 100.1 | kya |
| generation time | 25 | years |
| ghost → central chimpanzee admixture proportion | 2 | % |
| bonobo → central chimpanzee admixture proportion | 2 | % |
| central chimpanzee → western chimpanzee admixture proportion | 1.5 | % |
| western chimpanzee → central chimpanzee admixture proportion | 0.5 | % |
| migration rate between bonobo and ancestral common chimpanzee | $1\times10^{-7}$ | per generation |
| sampling time of source genome 1 (Ghost) | 0 | kya |
| sampling time of source genome 2 (Bonobo) | 0 | kya |
| mutation rate | $1.2\times10^{-8}$ | per base per generation |
| recombination rate | $0.7\times10^{-8}$ | per base per generation |

**Table S5. Parameters for the Papuan-Neanderthal-Denisovan model**

| Description | Value | Unit |
|---|---|---|
| reference population | YRI | |
| target population | Papuan | |
| source population 1 | Nea1 | |
| source population 2 | Den1 & Den2 | |
| African pop. size | 48433 | |
| European pop. size | 6962 | |
| East Asian pop. size | 9025 | |
| Papuan pop. size | 8834 | |
| Altai Denisovan (DenA) pop. size | 5083 | |
| Altai Neandertal (NeaA) pop. size | 826 | |
| Introgressing Denisovan D1 pop. size | 13249 | |
| Introgressing Denisovan D2 pop. size | 13249 | |
| Introgressing Neandertal pop. size | 13249 | |
| Ghost (out-of-Africa lineage) pop. size | 8516 | |
| (European + East Asian) pop. size before divergence | 12971 | |
| (Ghost + African) pop. size before divergence | 41563 | |
| (Denisovan + Neandertal) pop. size before divergence | 13249 | |
| (Human + Archaic) pop. size before divergence | 32671 | |
| (Altai Denisovan + Introgressing Denisovan D1) pop. size before divergence | 100 | |
| (Altai Denisovan + Introgressing Denisovan D2) pop. size before divergence | 100 | |
| (Altai Neandertal + Introgressing Neandertal) pop. size before divergence | 13249 | |
| (European + East Asian) bottleneck pop. size | 2231 | |
| Papuan bottleneck pop. size | 243 | |
| Ghost (out-of-Africa) bottleneck pop. size | 1394 | |
| Time of European and East Asian split | 37497 | years |
| Time of Ghost and (European + East Asian) split | 50982 | years |
| Time of Ghost and Papuan split | 51736 | years |
| Time of Ghost and African split | 64322 | years |
| Time of Altai Neandertal and Introgressing Neandertal split | 97875 | years |
| Time of Altai Denisovan and Introgressing Denisovan D1 split | 282750 | years |
| Time of Altai Denisovan and Introgressing Denisovan D2 split | 362500 | years |
| Time of Denisovan and Neandertal split | 437610 | years |
| Time of Human and Archaic split | 586525 | years |
| Time of the (European + East Asian) bottleneck | 48111 | years |
| Time of the Papuan bottleneck | 48865 | years |
| Time of the Ghost (out-of-Africa) bottleneck | 61451 | years |
| Time of Denisovan D1 to Papuan admixture | 29800 | years |
| Time of Denisovan D2 to Papuan admixture | 45700 | years |
| Time of Neandertal to Ghost admixture | 53737 | years |
| Time of Neandertal to (European + East Asian) admixture | 45414 | years |
| Time of Neandertal to Papuan admixture | 40948 | years |
| Time of Neandertal to East Asian admixture | 25607 | years |
| (European + East Asian)–Papuan migration begins | 37497 | years |
| (European + East Asian)–Ghost migration begins | 37497 | years |
| generation time | 29 | years |
| Amount of Denisovan D1 admixture in Papuan pop. | 2.2 | % |
| Amount of Denisovan D2 admixture in Papuan pop. | 1.8 | % |
| Amount of Neandertal admixture in Ghost pop. | 2.4 | % |
| Amount of Neandertal admixture in (European + East Asian) pop. | 1.1 | % |
| Amount of Neandertal admixture in Papuan pop. | 0.2 | % |
| Amount of Neandertal admixture in East Asian pop. | 0.2 | % |
| Ghost–African migration rate | $1.79\times10^{-4}$ | per generation |
| Ghost–European migration rate | $4.42\times10^{-4}$ | per generation |
| European–East Asian migration rate | $31.4\times10^{-4}$ | per generation |
| East Asian–Papuan migration rate | $57.2\times10^{-4}$ | per generation |
| (European + East Asian)–Papuan migration rate | $5.72\times10^{-4}$ | per generation |
| (European + East Asian)–Ghost migration rate | $4.42\times10^{-4}$ | per generation |
| sampling time of source genome 1 (NeaA) | 75748 | years |
| sampling time of source genome 2 (DenA) | 59682 | years |
| mutation rate | $1.4\times10^{-8}$ | per base per generation |
| recombination rate | $1\times10^{-8}$ | per base per generation |

**Table S6. Parameters for the Europe-Neanderthal-Denisovan model**

| Description | Value | Unit |
|---|---|---|
| reference population | Africa | |
| target population | Europe | |
| source population 1 | Den1 | |
| source population 2 | Nean2 | |
| Africa start size | 52000 | |
| Africa size (after Africa/Chimp split) | 7000 | |
| Africa size (after Africa/Den1 split) | 7000 | |
| Africa current size (after Africa/Europe split) | 27122 | |
| Chimp start size | 15000 | |
| Den1 start size | 5000 | |
| Den1 size (after Den1/Nean1 split) | 5000 | |
| Nean1 start size | 2000 | |
| Nean1 size (After Nean1/Nean2 split) | 1000 | |
| Den2 start size | 5000 | |
| Nean2 start size | 1000 | |
| Nean3 start size | 1000 | |
| Europe start size | 250 | |
| Europe grow up size | 5000 | |
| Europe bottleneck size | 1305 | |
| Europe current size | 5000 | |
| Africa/Chimp split time | 6000 | kya |
| Africa/Den1 split time | 580 | kya |
| Africa/Europe split time | 62 | kya |
| Den1/Nean1 split time | 420 | kya |
| Den1/Den2 split time | 335 | kya |
| Nean1/Nean2 split time | 130 | kya |
| Nean2/Nean3 split time | 95 | kya |
| Europe grow up start time | 59.1 | kya |
| Europe bottleneck start time | 56.2 | kya |
| Europe bottleneck end time | 53.3 | kya |
| Nean2 →Europe admixture proportion | 1.6 | % |
| Den1 →Europe admixture proportion | 0.4 | % |
| Nean2 →Europe admixture time | 50 | kya |
| Den1 →Europe admixture time | 50 | kya |
| generation time | 29 | years |
| sampling time of source genome 1 (Den2) | 80 | kya |
| sampling time of source genome 2 (Nea1) | 120 | kya |
| sampling time of source genome 3 (Nea3) | 60 | kya |
| mutation rate | $1.2\times10^{-8}$ | per base per generation |
| recombination rate | $1.2\times10^{-8}$ | per base per generation |

Table S7. Summary of Simulation Data Used for ArchaicSeeker 3.0 Training

Simulating data under the Bonobo-Ghost model

| reference samples | target samples | source samples | reapted times | usage |
|---|---|---|---|---|
| 10 | 100 | 1 | 10 | training |
| 50 | 100 | 1 | 10 | training |
| 100 | 100 | 1 | 10 | training |
| 10 | 1 | 1 | 100 | test |
| 50 | 1 | 1 | 100 | test |

Simulating data under the Human-Neanderthal model

| reference samples | target samples | source samples | reapted times | usage |
|---|---|---|---|---|
| 100 | 100 | 1 | 10 | training |
| 10 | 1 | 1 | 100 | test |
| 50 | 1 | 1 | 100 | test |

Simulating data under the Human-Denisovan model

| reference samples | target samples | source samples | reapted times | usage |
|---|---|---|---|---|
| 10 | 1 | 1 | 100 | test |
| 50 | 1 | 1 | 100 | test |

Simulating data under the Chimp-Bonobo-Ghost model

| reference samples | target samples | source1 samples | source1 samples | reapted times | usage |
|---|---|---|---|---|---|
| 100 | 100 | 1 | 1 | 10 | training |
| 10 | 1 | 1 | 1 | 100 | test |
| 50 | 1 | 1 | 1 | 100 | test |

Simulating data under the Papuan-Neanderthal-Denisovan model

| reference samples | target samples | source1 samples | source2 samples | reapted times | usage |
|---|---|---|---|---|---|
| 10 | 100 | 1 | 1 | 10 | training |
| 50 | 100 | 1 | 1 | 10 | training |
| 100 | 100 | 1 | 1 | 10 | training |
| 10 | 1 | 1 | 1 | 100 | test |
| 50 | 1 | 1 | 1 | 100 | test |

Simulating data under the Europe-Neanderthal-Denisovan model

| reference samples | target samples | source1 samples | source2 & 3 samples | Den1/Den2 split time(kya) | Nean2/Nean3 split time(kya) | Den1 introgression proportion(%) | Nean2 intogression time(%) | Den1 intogression time(kya) | Nean2 intogression time(kya) | mutation rate | recombination rate | reapted times | filter the sites | genetic map | usage |
|---|---|---|---|---|---|---|---|---|---|---|---|---|---|---|---|
| 100 | 100 | 1 | 2 | 335 | 95 | 2 | 0 | 34 | 50 | $1.2\times10^{-8}$ | $1.2\times10^{-8}$ | 10 | yes | none | training |
| 100 | 100 | 1 | 2 | 335 | 95 | 2 | 0 | 42 | 50 | $1.2\times10^{-8}$ | $1.2\times10^{-8}$ | 10 | yes | none | training |
| 100 | 100 | 1 | 2 | 335 | 95 | 2 | 0 | 50 | 50 | $1.2\times10^{-8}$ | $1.2\times10^{-8}$ | 10 | yes | none | training |
| 100 | 100 | 1 | 2 | 335 | 95 | 2 | 0 | 58 | 50 | $1.2\times10^{-8}$ | $1.2\times10^{-8}$ | 10 | yes | none | training |
| 100 | 100 | 1 | 2 | 301 | 95 | 2 | 0 | 50 | 50 | $1.2\times10^{-8}$ | $1.2\times10^{-8}$ | 10 | yes | none | training |
| 100 | 100 | 1 | 2 | 318 | 95 | 2 | 0 | 50 | 50 | $1.2\times10^{-8}$ | $1.2\times10^{-8}$ | 10 | yes | none | training |
| 100 | 100 | 1 | 2 | 352 | 95 | 2 | 0 | 50 | 50 | $1.2\times10^{-8}$ | $1.2\times10^{-8}$ | 10 | yes | none | training |
| 100 | 100 | 1 | 2 | 369 | 95 | 2 | 0 | 50 | 50 | $1.2\times10^{-8}$ | $1.2\times10^{-8}$ | 10 | yes | none | training |
| 100 | 100 | 1 | 2 | 335 | 95 | 0.4 | 0 | 50 | 50 | $1.2\times10^{-8}$ | $1.2\times10^{-8}$ | 10 | yes | none | training |
| 100 | 100 | 1 | 2 | 335 | 95 | 0.8 | 0 | 50 | 50 | $1.2\times10^{-8}$ | $1.2\times10^{-8}$ | 10 | yes | none | training |
| 100 | 100 | 1 | 2 | 335 | 95 | 1.2 | 0 | 50 | 50 | $1.2\times10^{-8}$ | $1.2\times10^{-8}$ | 10 | yes | none | training |
| 100 | 100 | 1 | 2 | 335 | 95 | 1.6 | 0 | 50 | 50 | $1.2\times10^{-8}$ | $1.2\times10^{-8}$ | 10 | yes | none | training |
| 100 | 100 | 1 | 2 | 335 | 95 | 0 | 2 | 50 | 34 | $1.2\times10^{-8}$ | $1.2\times10^{-8}$ | 10 | yes | none | training |
| 100 | 100 | 1 | 2 | 335 | 95 | 0 | 2 | 50 | 42 | $1.2\times10^{-8}$ | $1.2\times10^{-8}$ | 10 | yes | none | training |
| 100 | 100 | 1 | 2 | 335 | 95 | 0 | 2 | 50 | 50 | $1.2\times10^{-8}$ | $1.2\times10^{-8}$ | 10 | yes | none | training |
| 100 | 100 | 1 | 2 | 335 | 95 | 0 | 2 | 50 | 58 | $1.2\times10^{-8}$ | $1.2\times10^{-8}$ | 10 | yes | none | training |
| 100 | 100 | 1 | 2 | 335 | 85 | 0 | 2 | 50 | 50 | $1.2\times10^{-8}$ | $1.2\times10^{-8}$ | 10 | yes | none | training |
| 100 | 100 | 1 | 2 | 335 | 90 | 0 | 2 | 50 | 50 | $1.2\times10^{-8}$ | $1.2\times10^{-8}$ | 10 | yes | none | training |
| 100 | 100 | 1 | 2 | 335 | 100 | 0 | 2 | 50 | 50 | $1.2\times10^{-8}$ | $1.2\times10^{-8}$ | 10 | yes | none | training |
| 100 | 100 | 1 | 2 | 335 | 105 | 0 | 2 | 50 | 50 | $1.2\times10^{-8}$ | $1.2\times10^{-8}$ | 10 | yes | none | training |
| 100 | 100 | 1 | 2 | 335 | 95 | 0 | 0.4 | 50 | 50 | $1.2\times10^{-8}$ | $1.2\times10^{-8}$ | 10 | yes | none | training |
| 100 | 100 | 1 | 2 | 335 | 95 | 0 | 0.8 | 50 | 50 | $1.2\times10^{-8}$ | $1.2\times10^{-8}$ | 10 | yes | none | training |
| 100 | 100 | 1 | 2 | 335 | 95 | 0 | 1.2 | 50 | 50 | $1.2\times10^{-8}$ | $1.2\times10^{-8}$ | 10 | yes | none | training |
| 100 | 100 | 1 | 2 | 335 | 95 | 0 | 1.6 | 50 | 50 | $1.2\times10^{-8}$ | $1.2\times10^{-8}$ | 10 | yes | none | training |
| 100 | 100 | 1 | 2 | 335 | 95 | 0.4 | 1.6 | 34 | 34 | $1.2\times10^{-8}$ | $1.2\times10^{-8}$ | 10 | yes | none | training |
| 100 | 100 | 1 | 2 | 335 | 95 | 0.8 | 1.2 | 34 | 34 | $1.2\times10^{-8}$ | $1.2\times10^{-8}$ | 10 | yes | none | training |
| 100 | 100 | 1 | 2 | 335 | 95 | 1.2 | 0.8 | 34 | 34 | $1.2\times10^{-8}$ | $1.2\times10^{-8}$ | 10 | yes | none | training |
| 100 | 100 | 1 | 2 | 335 | 95 | 1.6 | 0.4 | 34 | 34 | $1.2\times10^{-8}$ | $1.2\times10^{-8}$ | 10 | yes | none | training |
| 100 | 100 | 1 | 2 | 335 | 95 | 0.4 | 1.6 | 34 | 50 | $1.2\times10^{-8}$ | $1.2\times10^{-8}$ | 10 | yes | none | training |
| 100 | 100 | 1 | 2 | 335 | 95 | 0.8 | 1.2 | 34 | 50 | $1.2\times10^{-8}$ | $1.2\times10^{-8}$ | 10 | yes | none | training |
| 100 | 100 | 1 | 2 | 335 | 95 | 1.2 | 0.8 | 34 | 50 | $1.2\times10^{-8}$ | $1.2\times10^{-8}$ | 10 | yes | none | training |
| 100 | 100 | 1 | 2 | 335 | 95 | 1.6 | 0.4 | 34 | 50 | $1.2\times10^{-8}$ | $1.2\times10^{-8}$ | 10 | yes | none | training |
| 100 | 100 | 1 | 2 | 335 | 95 | 0.4 | 1.6 | 34 | 58 | $1.2\times10^{-8}$ | $1.2\times10^{-8}$ | 10 | yes | none | training |
| 100 | 100 | 1 | 2 | 335 | 95 | 0.8 | 1.2 | 34 | 58 | $1.2\times10^{-8}$ | $1.2\times10^{-8}$ | 10 | yes | none | training |
| 100 | 100 | 1 | 2 | 335 | 95 | 1.2 | 0.8 | 34 | 58 | $1.2\times10^{-8}$ | $1.2\times10^{-8}$ | 10 | yes | none | training |
| 100 | 100 | 1 | 2 | 335 | 95 | 1.6 | 0.4 | 34 | 58 | $1.2\times10^{-8}$ | $1.2\times10^{-8}$ | 10 | yes | none | training |
| 100 | 100 | 1 | 2 | 335 | 95 | 0.4 | 1.6 | 50 | 34 | $1.2\times10^{-8}$ | $1.2\times10^{-8}$ | 10 | yes | none | training |
| 100 | 100 | 1 | 2 | 335 | 95 | 0.8 | 1.2 | 50 | 34 | $1.2\times10^{-8}$ | $1.2\times10^{-8}$ | 10 | yes | none | training |
| 100 | 100 | 1 | 2 | 335 | 95 | 1.2 | 0.8 | 50 | 34 | $1.2\times10^{-8}$ | $1.2\times10^{-8}$ | 10 | yes | none | training |
| 100 | 100 | 1 | 2 | 335 | 95 | 1.6 | 0.4 | 50 | 34 | $1.2\times10^{-8}$ | $1.2\times10^{-8}$ | 10 | yes | none | training |
| 100 | 100 | 1 | 2 | 335 | 95 | 0.4 | 1.6 | 50 | 50 | $1.2\times10^{-8}$ | $1.2\times10^{-8}$ | 10 | yes | none | training |
| 100 | 100 | 1 | 2 | 335 | 95 | 0.8 | 1.2 | 50 | 50 | $1.2\times10^{-8}$ | $1.2\times10^{-8}$ | 10 | yes | none | training |
| 100 | 100 | 1 | 2 | 335 | 95 | 1.2 | 0.8 | 50 | 50 | $1.2\times10^{-8}$ | $1.2\times10^{-8}$ | 10 | yes | none | training |
| 100 | 100 | 1 | 2 | 335 | 95 | 1.6 | 0.4 | 50 | 50 | $1.2\times10^{-8}$ | $1.2\times10^{-8}$ | 10 | yes | none | training |
| 100 | 100 | 1 | 2 | 335 | 95 | 0.4 | 1.6 | 50 | 58 | $1.2\times10^{-8}$ | $1.2\times10^{-8}$ | 10 | yes | none | training |
| 100 | 100 | 1 | 2 | 335 | 95 | 0.8 | 1.2 | 50 | 58 | $1.2\times10^{-8}$ | $1.2\times10^{-8}$ | 10 | yes | none | training |
| 100 | 100 | 1 | 2 | 335 | 95 | 1.2 | 0.8 | 50 | 58 | $1.2\times10^{-8}$ | $1.2\times10^{-8}$ | 10 | yes | none | training |
| 100 | 100 | 1 | 2 | 335 | 95 | 1.6 | 0.4 | 50 | 58 | $1.2\times10^{-8}$ | $1.2\times10^{-8}$ | 10 | yes | none | training |
| 100 | 100 | 1 | 2 | 335 | 95 | 0.4 | 1.6 | 58 | 34 | $1.2\times10^{-8}$ | $1.2\times10^{-8}$ | 10 | yes | none | training |
| 100 | 100 | 1 | 2 | 335 | 95 | 0.8 | 1.2 | 58 | 34 | $1.2\times10^{-8}$ | $1.2\times10^{-8}$ | 10 | yes | none | training |
| 100 | 100 | 1 | 2 | 335 | 95 | 1.2 | 0.8 | 58 | 34 | $1.2\times10^{-8}$ | $1.2\times10^{-8}$ | 10 | yes | none | training |
| 100 | 100 | 1 | 2 | 335 | 95 | 1.6 | 0.4 | 58 | 34 | $1.2\times10^{-8}$ | $1.2\times10^{-8}$ | 10 | yes | none | training |
| 100 | 100 | 1 | 2 | 335 | 95 | 0.4 | 1.6 | 58 | 50 | $1.2\times10^{-8}$ | $1.2\times10^{-8}$ | 10 | yes | none | training |
| 100 | 100 | 1 | 2 | 335 | 95 | 0.8 | 1.2 | 58 | 50 | $1.2\times10^{-8}$ | $1.2\times10^{-8}$ | 10 | yes | none | training |
| 100 | 100 | 1 | 2 | 335 | 95 | 1.2 | 0.8 | 58 | 50 | $1.2\times10^{-8}$ | $1.2\times10^{-8}$ | 10 | yes | none | training |
| 100 | 100 | 1 | 2 | 335 | 95 | 1.6 | 0.4 | 58 | 50 | $1.2\times10^{-8}$ | $1.2\times10^{-8}$ | 10 | yes | none | training |
| 100 | 100 | 1 | 2 | 335 | 95 | 0.4 | 1.6 | 58 | 58 | $1.2\times10^{-8}$ | $1.2\times10^{-8}$ | 10 | yes | none | training |
| 100 | 100 | 1 | 2 | 335 | 95 | 0.8 | 1.2 | 58 | 58 | $1.2\times10^{-8}$ | $1.2\times10^{-8}$ | 10 | yes | none | training |
| 100 | 100 | 1 | 2 | 335 | 95 | 1.2 | 0.8 | 58 | 58 | $1.2\times10^{-8}$ | $1.2\times10^{-8}$ | 10 | yes | none | training |
| 100 | 100 | 1 | 2 | 335 | 95 | 1.6 | 0.4 | 58 | 58 | $1.2\times10^{-8}$ | $1.2\times10^{-8}$ | 10 | yes | none | training |
| 10 | 100 | 1 | 2 | 335 | 95 | 1 | 1 | 50 | 50 | $1.2\times10^{-8}$ | $1.2\times10^{-8}$ | 10 | yes | none | training |
| 50 | 100 | 1 | 2 | 335 | 95 | 1 | 1 | 50 | 50 | $1.2\times10^{-8}$ | $1.2\times10^{-8}$ | 10 | yes | none | training |
| 100 | 100 | 1 | 2 | 335 | 95 | 1 | 1 | 50 | 50 | $1.2\times10^{-8}$ | $1.2\times10^{-8}$ | 10 | yes | none | training |
| 100 | 20 | 1 | 2 | 335 | 95 | 1 | 1 | 50 | 50 | $0.8\times10^{-8}$ | $0.8\times10^{-8}$ | 10 | yes | none | training |
| 100 | 20 | 1 | 2 | 335 | 95 | 1 | 1 | 50 | 50 | $0.8\times10^{-8}$ | $1.0\times10^{-8}$ | 10 | yes | none | training |
| 100 | 20 | 1 | 2 | 335 | 95 | 1 | 1 | 50 | 50 | $0.8\times10^{-8}$ | $1.2\times10^{-8}$ | 10 | yes | none | training |
| 100 | 20 | 1 | 2 | 335 | 95 | 1 | 1 | 50 | 50 | $0.8\times10^{-8}$ | $1.4\times10^{-8}$ | 10 | yes | none | training |
| 100 | 20 | 1 | 2 | 335 | 95 | 1 | 1 | 50 | 50 | $0.8\times10^{-8}$ | $1.6\times10^{-8}$ | 10 | yes | none | training |
| 100 | 20 | 1 | 2 | 335 | 95 | 1 | 1 | 50 | 50 | $1.0\times10^{-8}$ | $0.8\times10^{-8}$ | 10 | yes | none | training |
| 100 | 20 | 1 | 2 | 335 | 95 | 1 | 1 | 50 | 50 | $1.0\times10^{-8}$ | $1.0\times10^{-8}$ | 10 | yes | none | training |
| 100 | 20 | 1 | 2 | 335 | 95 | 1 | 1 | 50 | 50 | $1.0\times10^{-8}$ | $1.2\times10^{-8}$ | 10 | yes | none | training |
| 100 | 20 | 1 | 2 | 335 | 95 | 1 | 1 | 50 | 50 | $1.0\times10^{-8}$ | $1.4\times10^{-8}$ | 10 | yes | none | training |
| 100 | 20 | 1 | 2 | 335 | 95 | 1 | 1 | 50 | 50 | $1.0\times10^{-8}$ | $1.6\times10^{-8}$ | 10 | yes | none | training |
| 100 | 20 | 1 | 2 | 335 | 95 | 1 | 1 | 50 | 50 | $1.2\times10^{-8}$ | $0.8\times10^{-8}$ | 10 | yes | none | training |
| 100 | 20 | 1 | 2 | 335 | 95 | 1 | 1 | 50 | 50 | $1.2\times10^{-8}$ | $1.0\times10^{-8}$ | 10 | yes | none | training |
| 100 | 20 | 1 | 2 | 335 | 95 | 1 | 1 | 50 | 50 | $1.2\times10^{-8}$ | $1.2\times10^{-8}$ | 10 | yes | none | training |
| 100 | 20 | 1 | 2 | 335 | 95 | 1 | 1 | 50 | 50 | $1.2\times10^{-8}$ | $1.4\times10^{-8}$ | 10 | yes | none | training |
| 100 | 20 | 1 | 2 | 335 | 95 | 1 | 1 | 50 | 50 | $1.2\times10^{-8}$ | $1.6\times10^{-8}$ | 10 | yes | none | training |
| 100 | 20 | 1 | 2 | 335 | 95 | 1 | 1 | 50 | 50 | $1.4\times10^{-8}$ | $0.8\times10^{-8}$ | 10 | yes | none | training |
| 100 | 20 | 1 | 2 | 335 | 95 | 1 | 1 | 50 | 50 | $1.4\times10^{-8}$ | $1.0\times10^{-8}$ | 10 | yes | none | training |
| 100 | 20 | 1 | 2 | 335 | 95 | 1 | 1 | 50 | 50 | $1.4\times10^{-8}$ | $1.2\times10^{-8}$ | 10 | yes | none | training |
| 100 | 20 | 1 | 2 | 335 | 95 | 1 | 1 | 50 | 50 | $1.4\times10^{-8}$ | $1.4\times10^{-8}$ | 10 | yes | none | training |
| 100 | 20 | 1 | 2 | 335 | 95 | 1 | 1 | 50 | 50 | $1.4\times10^{-8}$ | $1.6\times10^{-8}$ | 10 | yes | none | training |
| 100 | 20 | 1 | 2 | 335 | 95 | 1 | 1 | 50 | 50 | $1.6\times10^{-8}$ | $0.8\times10^{-8}$ | 10 | yes | none | training |
| 100 | 20 | 1 | 2 | 335 | 95 | 1 | 1 | 50 | 50 | $1.6\times10^{-8}$ | $1.0\times10^{-8}$ | 10 | yes | none | training |
| 100 | 20 | 1 | 2 | 335 | 95 | 1 | 1 | 50 | 50 | $1.6\times10^{-8}$ | $1.2\times10^{-8}$ | 10 | yes | none | training |
| 100 | 20 | 1 | 2 | 335 | 95 | 1 | 1 | 50 | 50 | $1.6\times10^{-8}$ | $1.4\times10^{-8}$ | 10 | yes | none | training |
| 100 | 20 | 1 | 2 | 335 | 95 | 1 | 1 | 50 | 50 | $1.6\times10^{-8}$ | $1.6\times10^{-8}$ | 10 | yes | none | training |
| 100 | 100 | 1 | 2 | 335 | 95 | 1 | 1 | 50 | 50 | $1.2\times10^{-8}$ | $1.2\times10^{-8}$ | 10 | yes | none | test |
| 100 | 100 | 1 | 2 | 335 | 95 | 1 | 1 | 50 | 50 | $1.2\times10^{-8}$ | $1.2\times10^{-8}$ | 10 | no | HapMap | test |
| 10 | 1 | 1 | 2 | 335 | 95 | 1 | 1 | 50 | 50 | $1.2\times10^{-8}$ | $1.2\times10^{-8}$ | 100 | no | none | test |
| 50 | 1 | 1 | 2 | 335 | 95 | 1 | 1 | 50 | 50 | $1.2\times10^{-8}$ | $1.2\times10^{-8}$ | 100 | no | none | test |

**Table S8. The relationship between validation F1 score and number and variaty of datasets**

| Europe-Neanderthal-Denisovan Model | Papuan-Neanderthal-Denisovan Model | Chimp-Bonobo-Ghost Model | Human-Neanderthal Model | Bonobo-Ghost Model | # Datasets | # Samples | Validation f1 | # datasets that can be generalized |
|---|---|---|---|---|---|---|---|---|
| 1 | 0 | 0 | 0 | 0 | 1 | 80 | 0.653 | 24 |
| 1 | 0 | 0 | 0 | 0 | 1 | 800 | 0.688 | 24 |
| 1 | 0 | 0 | 0 | 0 | 10 | 800 | 0.723 | 24 |
| 1 | 0 | 0 | 0 | 0 | 24 | 19200 | 0.749 | 24 |
| 1 | 0 | 0 | 0 | 0 | 60 | 48000 | 0.783 | 89 |
| 1 | 0 | 0 | 0 | 0 | 88 | 5480 | 0.798 | 89 |
| 1 | 0 | 0 | 0 | 0 | 88 | 52400 | 0.814 | 89 |
| 1 | 1 | 0 | 0 | 0 | 91 | 54800 | 0.841 | 92 |
| 1 | 1 | 1 | 0 | 0 | 92 | 55600 | 0.874 | 93 |
| 1 | 1 | 1 | 1 | 0 | 93 | 56400 | 0.883 | 93 |
| 1 | 1 | 1 | 1 | 1 | 96 | 58800 | 0.902 | 96 |

**Table S9. Demographic Models Used for the Simulation Benchmark**

| Demographic Model | Species | Admixture Type | Source Type | Source_1 (Sampled time) | Source_2 (Sampled time) | Reference_population | Target_population | mu | r | Citation |
|---|---|---|---|---|---|---|---|---|---|---|
| WHG_Nean | Human | Single pulse | 1src | Neanderthal (2272) | NA | Mbuti | Loschbour | 1.29E-08 | 1.00E-08 | AncientEurasia_9K19[3] |
| Europe_Nean_HI | Human | Single pulse | 1src | Neanderthal (1906) | NA | YRI | CEU | 1.29E-08 | 1.00E-08 | HumanNeanderthal_4G21[4] |
| Europe_Nean_CGF | Human | Continuous gene flow | 1src | Neanderthal (NA) | NA | YRI | CEU | 1.29E-08 | 1.00E-08 | OutOfAfricaArchaicAdmixture_5R19[5] |
| Europe_Den | Human | Single pulse | 1src | Denisovan (1997) | NA | Africa | Europe | 1.20E-08 | 1.20E-08 | skov_HumanDenisovan_demography[2] |
| Europe_NeanDen | Human | Single pulse | 2src | Nean1 (4138); Nean3 (2069) | Den2 (2759) | Africa | Europe | 1.20E-08 | 1.20E-08 | AS2_demography[1] |
| Papuan_NeanDen | Human | Multiple pulse | 2src | Nea1 (2612) | Den1 (2058) | YRI | Papuans | 1.40E-08 | 1.00E-08 | HumanNeanderthalDenisovan_PapuansOutOfAfrica_10J19[6,7] |
| Europe_NeanArcAFR | Human | Multiple pulse | 2src | ArchaicAFR (NA) | Neanderthal (NA) | YRI | CEU | 1.40E-08 | 1.00E-08 | OutOfAfricaExtendedNeandertalAdmixturePulse_3I21[8] |
| Chimp_1src | Chimpanzee | Single pulse | 1src | Ghost (NA) | NA | Western | Bonobo | 1.20E-08 | 7.00E-09 | BonoboGhost_4K19[9] |
| Chimp_2src | Chimpanzee | Single pulse | 2src | Ghost (NA) | Bonobo (NA) | Western | Central | 1.20E-08 | 7.00E-09 | ChimpBonoboGhost_4K19[10,11] |


1 Yuan, K. *et al.* Refining models of archaic admixture in Eurasia with ArchaicSeeker 2.0. *Nat. Commun.* **12**, 6232 (2021).
2 Skov, L. *et al.* Detecting archaic introgression using an unadmixed outgroup. *PLOS Genet.* **14**, e1007641 (2018).
3 Kamm, J., Terhorst, J., Durbin, R. & Song, Y. S. Efficiently Inferring the Demographic History of Many Populations With Allele Count Data. *Journal of the American Statistical Association* (2020).
4 Gower, G., Picazo, P. I., Fumagalli, M. & Racimo, F. Detecting adaptive introgression in human evolution using convolutional neural networks. *eLife* **10**, e64669 (2021).
5 Ragsdale, A. P. & Gravel, S. Models of archaic admixture and recent history from two-locus statistics. *PLOS Genet.* **15**, e1008204 (2019).
6 Malaspinas, A.-S. *et al.* A genomic history of Aboriginal Australia. *Nature* **538**, 207–214 (2016).
7 Jacobs, G. S. *et al.* Multiple Deeply Divergent Denisovan Ancestries in Papuans. *Cell* **177**, 1010-1021.e32 (2019).
8 Iasi, L. N. M., Ringbauer, H. & Peter, B. M. An Extended Admixture Pulse Model Reveals the Limitations to Human–Neandertal Introgression Dating. *Mol. Biol. Evol.* **38**, 5156–5174 (2021).
9 Kuhlwilm, M., Han, S., Sousa, V. C., Excoffier, L. & Marques-Bonet, T. Ancient admixture from an extinct ape lineage into bonobos. *Nat Ecol Evol* **3**, 957–965 (2019).
10 Fontsere, C., de Manuel, M., Marques-Bonet, T. & Kuhlwilm, M. Admixture in Mammals and How to Understand Its Functional Implications. *Bioessays* **41**, 1900123 (2019).
11 Huang, X., Kruisz, P. & Kuhlwilm, M. sstar: A Python Package for Detecting Archaic Introgression from Population Genetic Data with *S**. *Mol. Biol. Evol.* **39**, msac212 (2022).

**Table S10. SNP Counts Used in Real-world Data Analysis**

| Chromosome | HGDPTGP | KGP2504 | ReferencePanel |
|---|---|---|---|
| 1 | 3821040 | 7981662 | 2734101 |
| 2 | 4317151 | 8624926 | 3108727 |
| 3 | 3578700 | 7081951 | 2549071 |
| 4 | 3493641 | 6984254 | 2443346 |
| 5 | 3252469 | 6452190 | 2302964 |
| 6 | 3048923 | 6026848 | 2152836 |
| 7 | 2753948 | 5764861 | 1922240 |
| 8 | 2840500 | 5530643 | 2031802 |
| 9 | 2120701 | 4465880 | 1524355 |
| 10 | 2407186 | 4855350 | 1716044 |
| 11 | 2407206 | 4849278 | 1701627 |
| 12 | 2278814 | 4638552 | 1598773 |
| 13 | 1785965 | 3536056 | 1269267 |
| 14 | 1580687 | 3213795 | 1129015 |
| 15 | 1398265 | 2937608 | 1011574 |
| 16 | 1516146 | 3299984 | 1109806 |
| 17 | 1247275 | 2813152 | 895845 |
| 18 | 1415995 | 2772464 | 1035694 |
| 19 | 879172 | 2188922 | 571074 |
| 20 | 1111282 | 2270455 | 822729 |
| 21 | 644297 | 1367700 | 464391 |
| 22 | 603915 | 1428207 | 444510 |
| **Total** | **48503278** | **99084738** | **34539791** |

**Table S11. Comparison of Archaic Introgression Detection Methods**

| Method | Analysis level | Modern human reference required | Archaic reference required | Enough homogeneous targets required | Archaic source distinguishable |
|---|---|---|---|---|---|
| **ArchaicSeeker 3.0** | Haplotype | Yes | Yes | No | Yes |
| **ArchaicSeeker 2.0**[1,2] | Haplotype | Yes | Yes | Yes | Yes |
| **IBDmix**[3] | Individual | No | Yes | Yes | No† |
| **hmmix**[4] | Haplotype | Yes | Yes | Yes | Conditional ‡ |
| **DAIseg**[5] | Haplotype | Yes | No | Yes | Conditional § |
| **Sprime**[6,7] | Population | Yes | Yes | Yes | Conditional ‡ |

**† IBDmix does not explicitly distinguish archaic sources and typically requires separate runs for different archaic references.**
**‡ hmmix and Sprime do not directly label archaic sources during inference; source assignment generally relies on post hoc analyses.**
**§ DAIseg does not require an explicit archaic reference but distinguishes archaic ancestry only when two distinct outgroup populations are specified, making source attribution conditional on study design.**


1 Yuan, K. *et al.* Refining models of archaic admixture in Eurasia with ArchaicSeeker 2.0. *Nat. Commun.* **12**, 6232 (2021).
2 Zhang, R., Yuan, K. & Xu, S. Detecting archaic introgression and modeling multiple-wave admixture with ArchaicSeeker 2.0. *STAR Protocols* **3**, 101314 (2022).
3 Chen, L., Wolf, A. B., Fu, W., Li, L. & Akey, J. M. Identifying and Interpreting Apparent Neanderthal Ancestry in African Individuals. *Cell* **180**, 677-687.e16 (2020).
4 Skov, L. *et al.* Detecting archaic introgression using an unadmixed outgroup. *PLOS Genet.* **14**, e1007641 (2018).
5 Planche, L. *et al.* An archaic reference-free method to jointly infer Neanderthal and Denisovan introgressed segments in modern human genomes. 2025.03.17.643330 Preprint at https://doi.org/10.1101/2025.03.17.643330 (2025).
6 Browning, S. R., Browning, B. L., Zhou, Y., Tucci, S. & Akey, J. M. Analysis of Human Sequence Data Reveals Two Pulses of Archaic Denisovan Admixture. *Cell* **173**, 53-61.e9 (2018).
7 Zhou, Y. & Browning, S. R. Protocol for detecting introgressed archaic variants with SPrime. *STAR Protocols* **2**, 100550 (2021).

**Table S12. Composition of the Reference Panel Used in ArchaicSeeker3**

| Reference | Population | Brief description | Project | Sample size |
|---|---|---|---|---|
| Modern – West Africa | YRI | Yoruba population from Ibadan, Nigeria | 1000 Genomes Project | 20 |
| | Yoruba | Yoruba population (additional sampling) | HGDP | 10 |
| | MSL | Mende population from Sierra Leone | 1000 Genomes Project | 14 |
| | GWD | Gambian population from Western Divisions | 1000 Genomes Project | 14 |
| | ESN | Esan population from Nigeria | 1000 Genomes Project | 14 |
| | Mandenka | West African Mandenka population | HGDP | 10 |
| Modern – East Africa | LWK | Luhya population from Kenya | 1000 Genomes Project | 24 |
| | BantuKenya | East African Bantu-speaking population | HGDP | 6 |
| Modern – Central Africa | Mbuti | Central African rainforest hunter-gatherers (Mbuti Pygmi | HGDP | 10 |
| | Biaka | Central African hunter-gatherers (Biaka Pygmies) | HGDP | 10 |
| Modern – Southern Africa | BantuSouthAfrica | Southern African Bantu-speaking population | HGDP | 8 |
| | San | Southern African Khoisan-speaking hunter-gatherers | HGDP | 6 |
| Archaic – Neanderthal | Altai Neandertal | High-coverage Neanderthal genome (Altai Mountains) | [1] | 1 |
| | Vindija33.19 | High-coverage Neanderthal genome (Croatia) | [2] | 1 |
| | Chagyrskaya-Phalan | Neanderthal genome from Chagyrskaya Cave | [3] | 1 |
| Archaic – Denisovan | Denisovan | Denisovan genome from Denisova Cave | [4] | 1 |

The reference panel consists of 146 modern African individuals spanning West, East, Central, and Southern Africa, and four archaic hominin genomes (three Neanderthals and one Denisovan).


1 Prüfer, K. *et al.* The complete genome sequence of a Neanderthal from the Altai Mountains. *Nature* **505**, 43–49 (2014).
2 Prüfer, K. *et al.* A high-coverage Neandertal genome from Vindija Cave in Croatia. *Science* **358**, 655–658 (2017).
3 Mafessoni, F. *et al.* A high-coverage Neandertal genome from Chagyrskaya Cave. *Proc. Natl. Acad. Sci. U.S.A.* **117**, 15132–15136 (2020).
4 Meyer, M. *et al.* A High-Coverage Genome Sequence from an Archaic Denisovan Individual. *Science* **338**, 222–226 (2012).

**Table S13. Samples in HGDPTGP**

| Region | Population | Sample_Count |
|---|---|---|
| AFR | ACB | 95 |
| AFR | ASW | 55 |
| AFR | BantuKenya | 10 |
| AFR | BantuSouthAfrica | 8 |
| AFR | Biaka | 23 |
| AFR | ESN | 106 |
| AFR | GWD | 119 |
| AFR | LWK | 97 |
| AFR | MSL | 88 |
| AFR | Mandenka | 21 |
| AFR | Mbuti | 12 |
| AFR | San | 6 |
| AFR | YRI | 121 |
| AFR | Yoruba | 21 |
| AMR | CLM | 97 |
| AMR | Colombian | 7 |
| AMR | Karitiana | 10 |
| AMR | MXL | 64 |
| AMR | Maya | 21 |
| AMR | PEL | 86 |
| AMR | PUR | 104 |
| AMR | Pima | 12 |
| AMR | Surui | 5 |
| CSA | BEB | 101 |
| CSA | Balochi | 24 |
| CSA | Brahui | 25 |
| CSA | Burusho | 24 |
| CSA | GIH | 101 |
| CSA | Hazara | 20 |
| CSA | ITU | 104 |
| CSA | Kalash | 22 |
| CSA | Makrani | 25 |
| CSA | PJL | 104 |
| CSA | Pathan | 24 |
| CSA | STU | 101 |
| CSA | Sindhi | 24 |
| EAS | CDX | 92 |
| EAS | CHB | 103 |
| EAS | CHS | 106 |
| EAS | Cambodian | 9 |
| EAS | Dai | 9 |
| EAS | Daur | 10 |
| EAS | Han | 33 |
| EAS | Hezhen | 9 |
| EAS | JPT | 104 |
| EAS | Japanese | 29 |
| EAS | KHV | 101 |
| EAS | Lahu | 8 |
| EAS | Miao | 10 |
| EAS | Mongolian | 10 |
| EAS | Naxi | 8 |
| EAS | NorthernHan | 10 |
| EAS | Oroqen | 9 |
| EAS | She | 10 |
| EAS | Tu | 10 |
| EAS | Tujia | 10 |
| EAS | Uygur | 10 |
| EAS | Xibo | 9 |
| EAS | Yakut | 25 |
| EAS | Yi | 10 |
| EUR | Adygei | 17 |
| EUR | Basque | 24 |
| EUR | BergamoItalian | 12 |
| EUR | CEU | 121 |
| EUR | FIN | 99 |
| EUR | French | 27 |
| EUR | GBR | 91 |
| EUR | IBS | 107 |
| EUR | Orcadian | 15 |
| EUR | Russian | 25 |
| EUR | Sardinian | 27 |
| EUR | TSI | 107 |
| EUR | Tuscan | 8 |
| MID | Bedouin | 46 |
| MID | Druze | 42 |
| MID | Mozabite | 27 |
| MID | Palestinian | 45 |
| OCE | Bougainville | 10 |
| OCE | PapuanHighlands | 9 |
| OCE | PapuanSepik | 8 |

**Table S14. Samples used in the global archaic introgression landscape analysis**

| Region | Population | Sample_Count |
|---|---|---|
| America | Cachi | 5 |
| America | Chane | 1 |
| America | Colla | 4 |
| America | Colombian | 104 |
| America | Karitiana | 11 |
| America | Maya | 21 |
| America | Mexican | 64 |
| America | Mixe | 3 |
| America | Mixtec | 2 |
| America | Peruvian | 86 |
| America | Pima | 13 |
| America | Puerto_Rican | 104 |
| America | Quechua | 3 |
| America | Surui | 6 |
| America | Tlingit | 4 |
| America | Wichi | 4 |
| America | Zapotec | 2 |
| Central Asia/Caucasus/Siberia | Abkhasian | 2 |
| Central Asia/Caucasus/Siberia | Abkhazian | 3 |
| Central Asia/Caucasus/Siberia | Adygei | 17 |
| Central Asia/Caucasus/Siberia | Aleut | 7 |
| Central Asia/Caucasus/Siberia | Altaian | 13 |
| Central Asia/Caucasus/Siberia | Armenian | 8 |
| Central Asia/Caucasus/Siberia | Avars | 3 |
| Central Asia/Caucasus/Siberia | Azerbaijanis | 3 |
| Central Asia/Caucasus/Siberia | Balkar | 3 |
| Central Asia/Caucasus/Siberia | Balochi | 24 |
| Central Asia/Caucasus/Siberia | Bashkirs | 5 |
| Central Asia/Caucasus/Siberia | Brahui | 25 |
| Central Asia/Caucasus/Siberia | Burusho | 24 |
| Central Asia/Caucasus/Siberia | Buryats | 17 |
| Central Asia/Caucasus/Siberia | Chechen | 1 |
| Central Asia/Caucasus/Siberia | Chukchi | 29 |
| Central Asia/Caucasus/Siberia | Chuvash | 3 |
| Central Asia/Caucasus/Siberia | Circassian | 3 |
| Central Asia/Caucasus/Siberia | Dolgan | 3 |
| Central Asia/Caucasus/Siberia | Eskimo | 26 |
| Central Asia/Caucasus/Siberia | Even | 18 |
| Central Asia/Caucasus/Siberia | Evenks | 13 |
| Central Asia/Caucasus/Siberia | Forest_Nenets | 3 |
| Central Asia/Caucasus/Siberia | Georgian | 4 |
| Central Asia/Caucasus/Siberia | Hazara | 20 |
| Central Asia/Caucasus/Siberia | Ishkasim | 2 |
| Central Asia/Caucasus/Siberia | Itelman | 1 |
| Central Asia/Caucasus/Siberia | Itelmen | 6 |
| Central Asia/Caucasus/Siberia | Kabardins | 4 |
| Central Asia/Caucasus/Siberia | Kalash | 22 |
| Central Asia/Caucasus/Siberia | Kalmyk | 10 |
| Central Asia/Caucasus/Siberia | Kazakhs | 3 |
| Central Asia/Caucasus/Siberia | Kets | 3 |
| Central Asia/Caucasus/Siberia | Khantys | 3 |
| Central Asia/Caucasus/Siberia | Koryak | 25 |
| Central Asia/Caucasus/Siberia | Kryashen_Tatars | 3 |
| Central Asia/Caucasus/Siberia | Kumyk | 3 |
| Central Asia/Caucasus/Siberia | Kyrgyz | 18 |
| Central Asia/Caucasus/Siberia | Lezgin | 6 |
| Central Asia/Caucasus/Siberia | Makrani | 25 |
| Central Asia/Caucasus/Siberia | Mansi | 12 |
| Central Asia/Caucasus/Siberia | Mishar_Tatars | 1 |
| Central Asia/Caucasus/Siberia | Nganasan | 13 |
| Central Asia/Caucasus/Siberia | North | 4 |
| Central Asia/Caucasus/Siberia | Pathan | 24 |
| Central Asia/Caucasus/Siberia | Rushan_Vanch | 2 |
| Central Asia/Caucasus/Siberia | Saami | 5 |
| Central Asia/Caucasus/Siberia | Sakha | 7 |
| Central Asia/Caucasus/Siberia | Selkup | 13 |
| Central Asia/Caucasus/Siberia | Shor | 2 |
| Central Asia/Caucasus/Siberia | Shugnan | 1 |
| Central Asia/Caucasus/Siberia | Tabasarans | 3 |
| Central Asia/Caucasus/Siberia | Tajik | 11 |
| Central Asia/Caucasus/Siberia | Tatars | 3 |
| Central Asia/Caucasus/Siberia | Tubalar | 22 |
| Central Asia/Caucasus/Siberia | Tundra_Nenets | 3 |
| Central Asia/Caucasus/Siberia | Turkmen | 10 |
| Central Asia/Caucasus/Siberia | Tuvinian | 13 |
| Central Asia/Caucasus/Siberia | Udmurds | 4 |
| Central Asia/Caucasus/Siberia | Ulchi | 25 |
| Central Asia/Caucasus/Siberia | Uygur | 11 |
| Central Asia/Caucasus/Siberia | Uzbek | 13 |
| Central Asia/Caucasus/Siberia | Vepsas | 4 |
| Central Asia/Caucasus/Siberia | Yaghnobi | 1 |
| Central Asia/Caucasus/Siberia | Yakut | 26 |
| Central Asia/Caucasus/Siberia | Yukagir | 19 |
| East Asia | Ami | 2 |
| East Asia | Atayal | 1 |
| East Asia | Dai | 101 |
| East Asia | Daur | 10 |
| East Asia | Han | 242 |
| East Asia | Hezhen | 9 |
| East Asia | Japanese | 133 |
| East Asia | Korean | 2 |
| East Asia | Lahu | 8 |
| East Asia | Miao | 10 |
| East Asia | Mongolian | 16 |
| East Asia | Naxi | 8 |
| East Asia | NorthernHan | 10 |
| East Asia | Oroqen | 9 |
| East Asia | She | 10 |
| East Asia | Tamang | 1 |
| East Asia | Tu | 10 |
| East Asia | Tujia | 10 |
| East Asia | Xibo | 9 |
| East Asia | Yi | 10 |
| North Africa | Mozabite | 27 |

| | | |
|---|---|---|
| North Africa | Saharawi | 2 |
| Oceania/North Philippine Negrito (Aeta, Agta, Batak) | Aeta | 3 |
| Oceania/North Philippine Negrito (Aeta, Agta, Batak) | Agta | 3 |
| Oceania/North Philippine Negrito (Aeta, Agta, Batak) | Australian | 2 |
| Oceania/North Philippine Negrito (Aeta, Agta, Batak) | Batak | 3 |
| Oceania/North Philippine Negrito (Aeta, Agta, Batak) | Bougainville | 10 |
| Oceania/North Philippine Negrito (Aeta, Agta, Batak) | Hawaiian | 1 |
| Oceania/North Philippine Negrito (Aeta, Agta, Batak) | Koinanbe | 3 |
| Oceania/North Philippine Negrito (Aeta, Agta, Batak) | Kosipe | 3 |
| Oceania/North Philippine Negrito (Aeta, Agta, Batak) | Maori | 1 |
| Oceania/North Philippine Negrito (Aeta, Agta, Batak) | PapuanHighlands | 9 |
| Oceania/North Philippine Negrito (Aeta, Agta, Batak) | PapuanSepik | 8 |
| South Asia | Asur | 1 |
| South Asia | Balija | 1 |
| South Asia | Bengali | 103 |
| South Asia | Brahmin | 5 |
| South Asia | Dhaka_mixed_popul | 3 |
| South Asia | Gond | 1 |
| South Asia | Gujarati | 101 |
| South Asia | Gupta | 1 |
| South Asia | Ho | 1 |
| South Asia | Irula | 2 |
| South Asia | Kapu | 3 |
| South Asia | Khonda_Dora | 1 |
| South Asia | Kol | 1 |
| South Asia | Kshatriya | 1 |
| South Asia | Kurmi | 1 |
| South Asia | Kusunda | 2 |
| South Asia | Madhya_Pradesh | 1 |
| South Asia | Madiga | 2 |
| South Asia | Mala | 2 |
| South Asia | Malayan | 1 |
| South Asia | Marwadi | 1 |
| South Asia | Orissa | 1 |
| South Asia | Punjab | 1 |
| South Asia | Punjabi | 104 |
| South Asia | Relli | 2 |
| South Asia | Santhal | 1 |
| South Asia | Sindhi | 24 |
| South Asia | SriLankan_Tamil | 101 |
| South Asia | Telugu | 104 |
| South Asia | Thakur | 1 |
| South Asia | Yadava | 2 |
| SoutheastAsia | Bajo | 4 |
| SoutheastAsia | Burmese | 10 |
| SoutheastAsia | Cambodian | 9 |
| SoutheastAsia | Dusun | 10 |
| SoutheastAsia | Igorot | 10 |
| SoutheastAsia | Kinh | 101 |
| SoutheastAsia | Lebbo | 4 |
| SoutheastAsia | Luzon | 2 |
| SoutheastAsia | Murut | 8 |
| SoutheastAsia | Thai | 2 |
| SoutheastAsia | Vietnamese | 10 |
| SoutheastAsia | Vizayan | 2 |
| West Eurasia | Albanian | 4 |
| West Eurasia | Arabs_Israel | 5 |
| West Eurasia | Assyrian | 3 |
| West Eurasia | Basque | 24 |
| West Eurasia | Bedouin | 46 |
| West Eurasia | Belarusian | 4 |
| West Eurasia | BergamoItalian | 12 |
| West Eurasia | British | 91 |
| West Eurasia | Bulgarian | 2 |
| West Eurasia | Cossacks | 4 |
| West Eurasia | Crete | 2 |
| West Eurasia | Croats | 4 |
| West Eurasia | Czech | 1 |
| West Eurasia | Druze | 45 |
| West Eurasia | Estonian | 8 |
| West Eurasia | Finnish | 102 |
| West Eurasia | French | 27 |
| West Eurasia | Germans | 3 |
| West Eurasia | Greek | 2 |
| West Eurasia | Hungarian | 4 |
| West Eurasia | Iberian | 107 |
| West Eurasia | Icelandic | 2 |
| West Eurasia | Ingrian | 3 |
| West Eurasia | Iranian | 6 |
| West Eurasia | Iraqi | 2 |
| West Eurasia | Jordanian | 5 |
| West Eurasia | Karelian | 3 |
| West Eurasia | Komis | 2 |
| West Eurasia | Latvian | 3 |
| West Eurasia | Lebanese | 1 |
| West Eurasia | Lithuanian | 3 |
| West Eurasia | Maris | 4 |
| West Eurasia | Moldavian | 2 |
| West Eurasia | Mordvins | 3 |
| West Eurasia | Northwest_European | 121 |
| West Eurasia | Norwegian | 1 |
| West Eurasia | Orcadian | 15 |
| West Eurasia | Palestinian | 45 |
| West Eurasia | Poles | 4 |
| West Eurasia | Polish | 1 |
| West Eurasia | Roma | 3 |
| West Eurasia | Russian | 32 |
| West Eurasia | Samaritan | 1 |
| West Eurasia | Sardinian | 27 |
| West Eurasia | Saudi | 2 |
| West Eurasia | Swedes | 2 |
| West Eurasia | Turkish | 2 |
| West Eurasia | Tuscan | 115 |
| West Eurasia | Ukrainian | 7 |
| West Eurasia | Yemenite | 2 |

**Table S15. Default model, inference, decoding, and annotation parameters used in ArchaicSeeker 3.0.**

| Parameter | Value used in this study | Module / stage | Description | Source / note |
|---|---|---|---|---|
| Input dimension | 7 | Base model (Mamba) | Six reference-compressed match channels (AFR/DEN/NEAN max/mean) plus one relative distance channel | Matches the final Methods description and code implementation |
| Output dimension | 3 | Base model / smoother | Three site-level ancestry states used for inference | Code-defined architecture |
| Model dimension | 192 | Base model (Mamba) | Hidden channel dimension of the Mamba backbone | Code-defined architecture |
| Number of Mamba layers | 8 | Base model (Mamba) | Depth of the long-context backbone | Code-defined architecture |
| Expansion factor | 2 | Base model (Mamba) | Channel expansion used within Mamba blocks | Code-defined architecture |
| State dimension (d_state) | 64 | Base model (Mamba) | State-space dimension in the Mamba backbone | Code-defined architecture |
| Convolution kernel (d_conv) | 4 | Base model (Mamba) | Local convolution width inside Mamba blocks | Code-defined architecture |
| Head dimension | 48 | Base model (Mamba) | Head dimension used in the Mamba-2 implementation | Code-defined architecture |
| Window length (W) | 4,096 SNPs | Base model / windowed inference | Context length processed by the long-context backbone | Main framework setting |
| Inference stride (S) | 512 SNPs | Overlap-aware reassembly | Step size for overlapping sliding-window inference | Main framework setting |
| Smoother input dimension | 4 | Boundary smoother | Three reassembled logit channels plus one position-derived channel | Matches the final Methods description and code implementation |
| Smoother kernel size (k) | 8,192 | Boundary smoother | Receptive field spans two adjacent 4,096-SNP windows for cross-window refinement | Main framework setting |
| Maximum gap (G) | 1 Mb | Decoding | Splits candidate tracts when consecutive archaic-labeled SNPs are separated by a large physical interval | Decoding rule |
| Pre-merge minimum length | 5 kb | Decoding / pre-merge filtering | Removes short fragmented calls before merging | Guided by Fig. S3.7 |
| Pre-merge minimum score | None | Decoding / pre-merge filtering | No additional score threshold applied before merging | Guided by Fig. S3.7 |
| Merge distance (\delta) | 10 kb | Decoding / merging | Merges nearby archaic sub-blocks into a tract | Guided by Fig. S3.7 |
| Post-merge minimum score | 0.85 | Decoding / post-merge filtering | Default score threshold for retaining merged segments | Guided by Fig. S3.8 |
| Post-merge minimum length | 15 kb | Decoding / post-merge filtering | Default length threshold for retaining merged segments | Guided by Fig. S3.8 |
| Mosaic threshold ( $\tau$ ) | 0.2 | Segment annotation | Segment-level annotation threshold used after tract construction to label merged segments with sufficient mixed-source support as mosaic | Post hoc annotation rule; not a training output state |

### Table S16. High-frequency AS3-specific introgressed candidate regions.

| Source | Chr | Start | End | Length (bp) | Population | Super-population/region | Frequency | Gene annotation |
|---|---|---|---|---|---|---|---|---|
| Neanderthal | chr2 | 54,519,229 | 54,559,478 | 40,249 | CEU | EUR | 36.4% | SPTBN1,SPTBN1-AS1,RPL23AP32,ENSG00000234943 |
| Neanderthal | chr2 | 54,519,229 | 54,559,478 | 40,249 | BEB | SAS | 38.4% | SPTBN1,SPTBN1-AS1,RPL23AP32,ENSG00000234943 |
| Neanderthal | chr2 | 54,519,229 | 54,559,478 | 40,249 | ITU | SAS | 41.2% | SPTBN1,SPTBN1-AS1,RPL23AP32,ENSG00000234943 |
| Neanderthal | chr2 | 227,704,474 | 227,720,640 | 16,166 | ITU | SAS | 40.2% | SLC19A3,SCYGR6,SCYGR7 |
| Neanderthal | chr2 | 227,704,474 | 227,720,640 | 16,166 | STU | SAS | 34.3% | SLC19A3,SCYGR6,SCYGR7 |
| Neanderthal | chr3 | 1,810,748 | 1,863,073 | 52,325 | GBR | EUR | 34.1% | NA |
| Neanderthal | chr3 | 143,494,899 | 143,529,739 | 34,840 | CEU | EUR | 35.4% | SLC9A9,GAPDHP47,ST13P15,ENSG00000303585 |
| Neanderthal | chr3 | 143,494,899 | 143,529,605 | 34,706 | FIN | EUR | 30.8% | SLC9A9,GAPDHP47,ST13P15,ENSG00000303585 |
| Neanderthal | chr3 | 143,494,899 | 143,529,605 | 34,706 | GBR | EUR | 34.6% | SLC9A9,GAPDHP47,ST13P15,ENSG00000303585 |
| Neanderthal | chr3 | 143,504,304 | 143,529,739 | 25,435 | TSI | EUR | 30.8% | SLC9A9,GAPDHP47,ST13P15,ENSG00000303585 |
| Neanderthal | chr6 | 65,600,024 | 65,617,277 | 17,253 | CLM | AMR | 41.5% | EYS |
| Neanderthal | chr16 | 15,852,921 | 15,856,149 | 3,228 | STU | SAS | 35.8% | MYH11 |
| Neanderthal | chr16 | 15,852,921 | 15,897,592 | 44,671 | CDX | EAS | 53.8% | MYH11,CEP20,RNU6-213P,ENSG00000262171 |
| Neanderthal | chr16 | 15,852,921 | 15,897,592 | 44,671 | CHS | EAS | 44.3% | MYH11,CEP20,RNU6-213P,ENSG00000262171 |
| Neanderthal | chr16 | 15,852,921 | 15,856,149 | 3,228 | JPT | EAS | 50.0% | MYH11 |
| Neanderthal | chr16 | 15,852,921 | 15,897,592 | 44,671 | KHV | EAS | 48.5% | MYH11,CEP20,RNU6-213P,ENSG00000262171 |
| Neanderthal | chr17 | 9,844,362 | 9,895,845 | 51,483 | CEU | EUR | 45.5% | GSG1L2,GLP2R,NPM1P45,RCVRN |
| Neanderthal | chr18 | 1,956,911 | 1,965,582 | 8,671 | BEB | SAS | 30.2% | ENSG00000266602,ENSG00000263745 |
| Neanderthal | chr18 | 1,952,175 | 1,965,582 | 13,407 | CHB | EAS | 33.5% | ENSG00000266602,ENSG00000263745 |
| Neanderthal | chr18 | 1,952,175 | 1,965,582 | 13,407 | JPT | EAS | 32.2% | ENSG00000266602,ENSG00000263745 |
| Neanderthal | chr18 | 74,152,857 | 74,194,084 | 41,227 | CEU | EUR | 47.0% | TIMM21,ENSG00000303326 |
| Neanderthal | chr18 | 74,173,279 | 74,194,084 | 20,805 | IBS | EUR | 47.7% | ENSG00000303326 |
| Neanderthal | chr18 | 74,178,005 | 74,194,084 | 16,079 | PUR | AMR | 41.8% | ENSG00000303326 |
| Neanderthal | chr18 | 77,411,928 | 77,497,322 | 85,394 | FIN | EUR | 41.4% | ENSG00000305981,ENSG00000264015,ENSG00000199392,BDP1P |
| Neanderthal | chr18 | 77,411,928 | 77,497,322 | 85,394 | IBS | EUR | 35.5% | ENSG00000305981,ENSG00000264015,ENSG00000199392,BDP1P |
| Neanderthal | chr20 | 63,697,389 | 63,769,790 | 72,401 | CDX | EAS | 67.2% | RTEL1-TNFRSF6B,TNFRSF6B,ARFRP1,ZGPAT,ENSG00000273154,ENSG00000274501,LIME1,ENSG00000273047,SLC2A4RG,ZBTB46,ZBTB46-AS2,ENSG00000298747,ZBTB46-AS1 |
| Neanderthal | chr20 | 63,697,389 | 63,769,790 | 72,401 | CHB | EAS | 68.0% | RTEL1-TNFRSF6B,TNFRSF6B,ARFRP1,ZGPAT,ENSG00000273154,ENSG00000274501,LIME1,ENSG00000273047,SLC2A4RG,ZBTB46,ZBTB46-AS2 |
| Neanderthal | chr20 | 63,697,389 | 63,769,790 | 72,401 | CHS | EAS | 67.1% | RTEL1-TNFRSF6B,TNFRSF6B,ARFRP1,ZGPAT,ENSG00000273154,ENSG00000274501,LIME1,ENSG00000273047,SLC2A4RG,ZBTB46,ZBTB46-AS2,ENSG00000298747,ZBTB46-AS1 |
| Neanderthal | chr20 | 63,697,389 | 63,769,790 | 72,401 | JPT | EAS | 62.0% | RTEL1-TNFRSF6B,TNFRSF6B,ARFRP1,ZGPAT,ENSG00000273154,ENSG00000274501,LIME1,ENSG00000273047,SLC2A4RG,ZBTB46,ZBTB46-AS2,ENSG00000298747,ZBTB46-AS1 |
| Neanderthal | chr20 | 63,697,389 | 63,769,790 | 72,401 | KHV | EAS | 61.1% | RTEL1-TNFRSF6B,TNFRSF6B,ARFRP1,ZGPAT,ENSG00000273154,ENSG00000274501,LIME1,ENSG00000273047,SLC2A4RG,ZBTB46,ZBTB46-AS2,ENSG00000298747,ZBTB46-AS1 |
| Neanderthal | chr20 | 63,697,389 | 63,769,790 | 72,401 | PEL | AMR | 52.4% | RTEL1-TNFRSF6B,TNFRSF6B,ARFRP1,ZGPAT,ENSG00000273154,ENSG00000274501,LIME1,ENSG00000273047,SLC2A4RG,ZBTB46,ZBTB46-AS2,ENSG00000298747 |
| Denisovan | chr1 | 59,414,422 | 59,423,858 | 9,436 | BEB | SAS | 41.3% | FGGY |
| Denisovan | chr1 | 59,414,422 | 59,423,858 | 9,436 | GIH | SAS | 37.4% | FGGY |
| Denisovan | chr1 | 59,414,422 | 59,423,858 | 9,436 | ITU | SAS | 37.3% | FGGY |
| Denisovan | chr1 | 59,414,422 | 59,423,858 | 9,436 | CDX | EAS | 33.9% | FGGY |
| Denisovan | chr1 | 59,414,422 | 59,423,858 | 9,436 | CHB | EAS | 37.9% | FGGY |
| Denisovan | chr1 | 59,414,422 | 59,423,858 | 9,436 | JPT | EAS | 33.2% | FGGY |
| Denisovan | chr1 | 59,414,422 | 59,423,858 | 9,436 | KHV | EAS | 37.9% | FGGY |
| Denisovan | chr4 | 58,286,198 | 58,309,838 | 23,640 | MXL | AMR | 35.2% | NA |
| Denisovan | chr4 | 58,286,198 | 58,347,233 | 61,035 | FIN | EUR | 36.4% | NA |
| Denisovan | chr4 | 58,286,198 | 58,309,838 | 23,640 | PEL | AMR | 35.9% | NA |
| Denisovan | chr5 | 24,564,746 | 24,584,002 | 19,256 | CHB | EAS | 61.7% | CDH10,, |
| Denisovan | chr5 | 24,564,746 | 24,584,002 | 19,256 | PJL | SAS | 51.0% | CDH10,, |
| Denisovan | chr5 | 24,564,746 | 24,584,002 | 19,256 | CDX | EAS | 54.3% | CDH10,, |
| Denisovan | chr5 | 24,564,746 | 24,584,002 | 19,256 | KHV | EAS | 56.6% | CDH10,, |
| Denisovan | chr5 | 24,564,746 | 24,584,002 | 19,256 | CHS | EAS | 62.4% | CDH10,, |
| Denisovan | chr5 | 24,564,746 | 24,584,002 | 19,256 | CLM | AMR | 38.8% | CDH10,, |
| Denisovan | chr5 | 24,564,746 | 24,584,002 | 19,256 | MXL | AMR | 39.8% | CDH10,, |
| Denisovan | chr5 | 24,564,746 | 24,584,002 | 19,256 | GIH | SAS | 49.0% | CDH10,, |
| Denisovan | chr5 | 24,564,746 | 24,584,002 | 19,256 | STU | SAS | 45.6% | CDH10,, |
| Denisovan | chr5 | 24,564,746 | 24,584,002 | 19,256 | GBR | EUR | 57.1% | CDH10,, |
| Denisovan | chr5 | 24,564,746 | 24,584,002 | 19,256 | IBS | EUR | 54.2% | CDH10,, |
| Denisovan | chr5 | 24,564,746 | 24,584,002 | 19,256 | BEB | SAS | 53.5% | CDH10,, |
| Denisovan | chr5 | 24,564,746 | 24,584,002 | 19,256 | ITU | SAS | 45.6% | CDH10,, |
| Denisovan | chr5 | 24,564,746 | 24,584,002 | 19,256 | CEU | EUR | 48.5% | CDH10,, |
| Denisovan | chr5 | 24,564,746 | 24,584,002 | 19,256 | FIN | EUR | 58.6% | CDH10,, |
| Denisovan | chr5 | 24,564,746 | 24,584,002 | 19,256 | TSI | EUR | 52.8% | CDH10,, |
| Denisovan | chr5 | 24,564,746 | 24,584,002 | 19,256 | PUR | AMR | 40.4% | CDH10,, |
| Denisovan | chr5 | 88,504,904 | 88,554,704 | 49,800 | JPT | EAS | 46.2% | MIR9-2HG |
| Denisovan | chr5 | 88,504,904 | 88,554,704 | 49,800 | CHS | EAS | 39.5% | MIR9-2HG |
| Denisovan | chr5 | 88,504,904 | 88,554,704 | 49,800 | CHB | EAS | 46.6% | MIR9-2HG |
| Denisovan | chr5 | 88,504,904 | 88,554,704 | 49,800 | CDX | EAS | 42.5% | MIR9-2HG |
| Denisovan | chr5 | 129,344,410 | 129,357,949 | 13,539 | CLM | AMR | 38.3% | ,ADAMTS19-AS1 |
| Denisovan | chr5 | 129,344,410 | 129,357,949 | 13,539 | IBS | EUR | 40.2% | ,ADAMTS19-AS1 |
| Denisovan | chr5 | 129,344,410 | 129,357,949 | 13,539 | PUR | AMR | 33.7% | ,ADAMTS19-AS1 |
| Denisovan | chr5 | 129,345,042 | 129,357,949 | 12,907 | TSI | EUR | 38.3% | ,ADAMTS19-AS1 |
| Denisovan | chr5 | 137,360,488 | 137,376,387 | 15,899 | CDX | EAS | 34.4% | SPOCK1 |
| Denisovan | chr6 | 701,631 | 709,753 | 8,122 | PEL | AMR | 33.5% | , |
| Denisovan | chr6 | 6,098,939 | 6,122,140 | 23,201 | FIN | EUR | 36.4% | NA |
| Denisovan | chr6 | 6,098,939 | 6,122,140 | 23,201 | STU | SAS | 36.3% | NA |
| Denisovan | chr6 | 147,642,544 | 147,661,994 | 19,450 | CHS | EAS | 31.4% | SAMD5, |
| Denisovan | chr7 | 36,087,020 | 36,137,669 | 50,649 | GBR | EUR | 44.0% | MARK2P13,,,,,EEPD1 |
| Denisovan | chr7 | 105,506,862 | 105,530,709 | 23,847 | ITU | SAS | 33.8% | PUS7,, |
| Denisovan | chr8 | 32,558,164 | 32,584,421 | 26,257 | PEL | AMR | 42.9% | NRG1 |
| Denisovan | chr15 | 48,555,536 | 48,603,494 | 47,958 | JPT | EAS | 37.5% | FBN1 |
| Denisovan | chr16 | 25,320,369 | 25,344,175 | 23,806 | BEB | SAS | 30.8% | ZKSCAN2-DT |
| Denisovan | chr22 | 38,884,941 | 38,906,169 | 21,228 | PJL | SAS | 37.0% | NA |
| Denisovan | chr22 | 38,884,941 | 38,906,169 | 21,228 | CLM | AMR | 48.4% | NA |
| Denisovan | chr22 | 38,884,941 | 38,906,169 | 21,228 | CEU | EUR | 40.4% | NA |
| Denisovan | chr22 | 38,884,941 | 38,906,169 | 21,228 | FIN | EUR | 32.8% | NA |
| Denisovan | chr22 | 38,884,941 | 38,906,079 | 21,138 | GBR | EUR | 34.6% | NA |
| Denisovan | chr22 | 38,884,941 | 38,906,169 | 21,228 | IBS | EUR | 39.7% | NA |
| Denisovan | chr22 | 38,884,941 | 38,906,169 | 21,228 | TSI | EUR | 37.9% | NA |

| Source | Chr | Start | End | Length (bp) | Population | Super-population/region | Frequency | Gene annotation |
|---|---|---|---|---|---|---|---|---|
| Denisovan | chr22 | 38,884,941 | 38,905,958 | 21,017 | BEB | SAS | 31.4% | NA |
| Denisovan | chr22 | 38,884,941 | 38,906,169 | 21,228 | ITU | SAS | 34.8% | NA |
| Denisovan | chr22 | 38,884,941 | 38,906,169 | 21,228 | STU | SAS | 38.2% | NA |
| Denisovan | chr22 | 38,884,941 | 38,906,169 | 21,228 | MXL | AMR | 60.2% | NA |
| Denisovan | chr22 | 38,884,941 | 38,906,169 | 21,228 | PEL | AMR | 81.8% | NA |
| Denisovan | chr22 | 38,884,941 | 38,906,079 | 21,138 | PUR | AMR | 38.0% | NA |
| Denisovan | chr22 | 45,309,502 | 45,389,055 | 79,553 | IBS | EUR | 57.0% | FAM118A,SMC1B,RIBC2 |
| Denisovan | chr22 | 45,309,502 | 45,389,055 | 79,553 | GBR | EUR | 67.0% | FAM118A,SMC1B |
| Denisovan | chr22 | 45,309,502 | 45,389,055 | 79,553 | TSI | EUR | 55.6% | FAM118A,SMC1B,RIBC2 |
| Denisovan | chr22 | 45,309,502 | 45,389,055 | 79,553 | CHS | EAS | 39.5% | FAM118A,SMC1B,RIBC2 |
| Denisovan | chr22 | 45,309,502 | 45,389,055 | 79,553 | JPT | EAS | 51.9% | FAM118A,SMC1B,RIBC2 |
| Denisovan | chr22 | 45,309,502 | 45,389,055 | 79,553 | PEL | AMR | 58.2% | FAM118A,SMC1B,RIBC2 |
| Denisovan | chr22 | 45,309,502 | 45,389,055 | 79,553 | PUR | AMR | 41.8% | FAM118A,SMC1B,RIBC2 |
| Denisovan | chr22 | 45,312,742 | 45,389,055 | 76,313 | CEU | EUR | 61.1% | FAM118A,SMC1B |
| Denisovan | chr22 | 45,320,006 | 45,389,055 | 69,049 | FIN | EUR | 59.1% | FAM118A,SMC1B |
| Denisovan | chr22 | 45,322,746 | 45,389,055 | 66,309 | MXL | AMR | 56.2% | FAM118A,SMC1B |
| Denisovan | chr6 | 701,536 | 709,753 | 8,217 | PEL | AMR | 33.5% | , |
| Denisovan | chr12 | 52,419,765 | 52,486,587 | 66,822 | TSI | EUR | 38.8% | ,KRT90P,KRT75,KRT6B,KRT6C |
| Denisovan | chr12 | 52,453,131 | 52,487,215 | 34,084 | CHS | EAS | 48.1% | KRT6B,KRT6C,KRT6A |
| Denisovan | chr12 | 52,453,131 | 52,475,232 | 22,101 | JPT | EAS | 66.8% | KRT6B,KRT6C |
| Denisovan | chr12 | 52,460,074 | 52,487,215 | 27,141 | KHV | EAS | 42.9% | KRT6C,KRT6A |
| Denisovan | chr22 | 45,320,006 | 45,389,055 | 69,049 | GBR | EUR | 65.9% | FAM118A,SMC1B,RIBC2 |
| Denisovan | chr22 | 45,320,006 | 45,389,055 | 69,049 | PEL | AMR | 58.2% | FAM118A,SMC1B,RIBC2 |
| Denisovan | chr3 | 128,699,301 | 128,835,265 | 135,964 | Papuans | Oceania | 33.3% | RPN1,Metazoa_SRP,POU5F1P6,RAB7A,,,,MARK2P8,FTH1P4,RN7SL698P,RPS15AP16 |
| Denisovan | chr3 | 194,522,489 | 194,541,469 | 18,980 | Papuans | Oceania | 50.0% | ,RNU6-1101P |
| Denisovan | chr7 | 7,003,357 | 7,042,260 | 38,903 | Papuans | Oceania | 37.5% | NA |
| Denisovan | chr11 | 30,668,800 | 30,706,896 | 38,096 | Papuans | Oceania | 50.0% | NA |
